# TNFSF13 insufficiency disrupts human colonic epithelial cell-mediated B cell differentiation

**DOI:** 10.1101/2024.09.23.614260

**Authors:** Xianghui Ma, Noor Dawany, Ayano Kondo, Kelly Maurer, Tatiana Karakasheva, Rawan Shraim, Patrick A. Williams, Louis R. Parham, Lauren A. Simon, Charles H. Danan, Maire A. Conrad, David A. Piccoli, Marcella Devoto, Kathleen E. Sullivan, Klaus H. Kaestner, Judith R. Kelsen, Kathryn E. Hamilton

**Affiliations:** Division of Gastroenterology, Hepatology, and Nutrition; Department of Pediatrics; Children’s Hospital of Philadelphia; Philadelphia, PA, 19104, USA; Department of Biomedical and Health Informatics; Children’s Hospital of Philadelphia; Philadelphia, PA, 19104, USA; Department of Genetics and Center for Molecular Studies in Digestive and Liver Diseases, Perelman School of Medicine, University of Pennsylvania, Philadelphia, Pennsylvania, Philadelphia, PA, 19104, USA; Division of Allergy Immunology, Children’s Hospital of Philadelphia, Philadelphia, PA, 19104, USA; Institute for Research in Genetics and Biomedicine, CNR, Cagliari, Italy, and Department of Translational and Precision Medicine, University Sapienza, Rome, Italy; Institute for Regenerative Medicine, University of Pennsylvania, Philadelphia, PA, 19104, USA

## Abstract

Cytokines mediating epithelial and immune cell interactions modulate mucosal healing-a process that goes awry with chronic inflammation as in inflammatory bowel disease. TNFSF13 is a cytokine important for B cell maturation and function, but roles for epithelial TNFSF13 and putative contribution to inflammatory bowel disease are poorly understood. We evaluated functional consequences of a novel monoallelic *TNFSF13* variant using biopsies, tissue-derived colonoids and induced pluripotent stem cell (iPSC)-derived colon organoids. *TNFSF13* variant colonoids exhibited a >50% reduction in secreted TNFSF13, increased epithelial proliferation, and reduced apoptosis, which was confirmed in iPSC-derived colon organoids. Single cell RNA-sequencing, flow cytometry, and co-immunoprecipitation identified FAS as the predominant colonic epithelial receptor for TNFSF13. Imaging mass cytometry revealed an increase in epithelial-associated B cells in *TNFSF13* variant colon tissue sections. Finally, *TNFSF13* variant colonoids co-cultured with memory B cells demonstrated a reduction in the production of IgA+ plasma cells compared to control colonoid co-cultures. Our findings support a role for epithelial TNFSF13 as a regulator of colonic epithelial growth and epithelial crosstalk with B cells.

**SUMMARY:** Epithelial TNFSF13 regulates colonic epithelial growth and epithelial-B cell interactions.

**Graphical Abstract:** 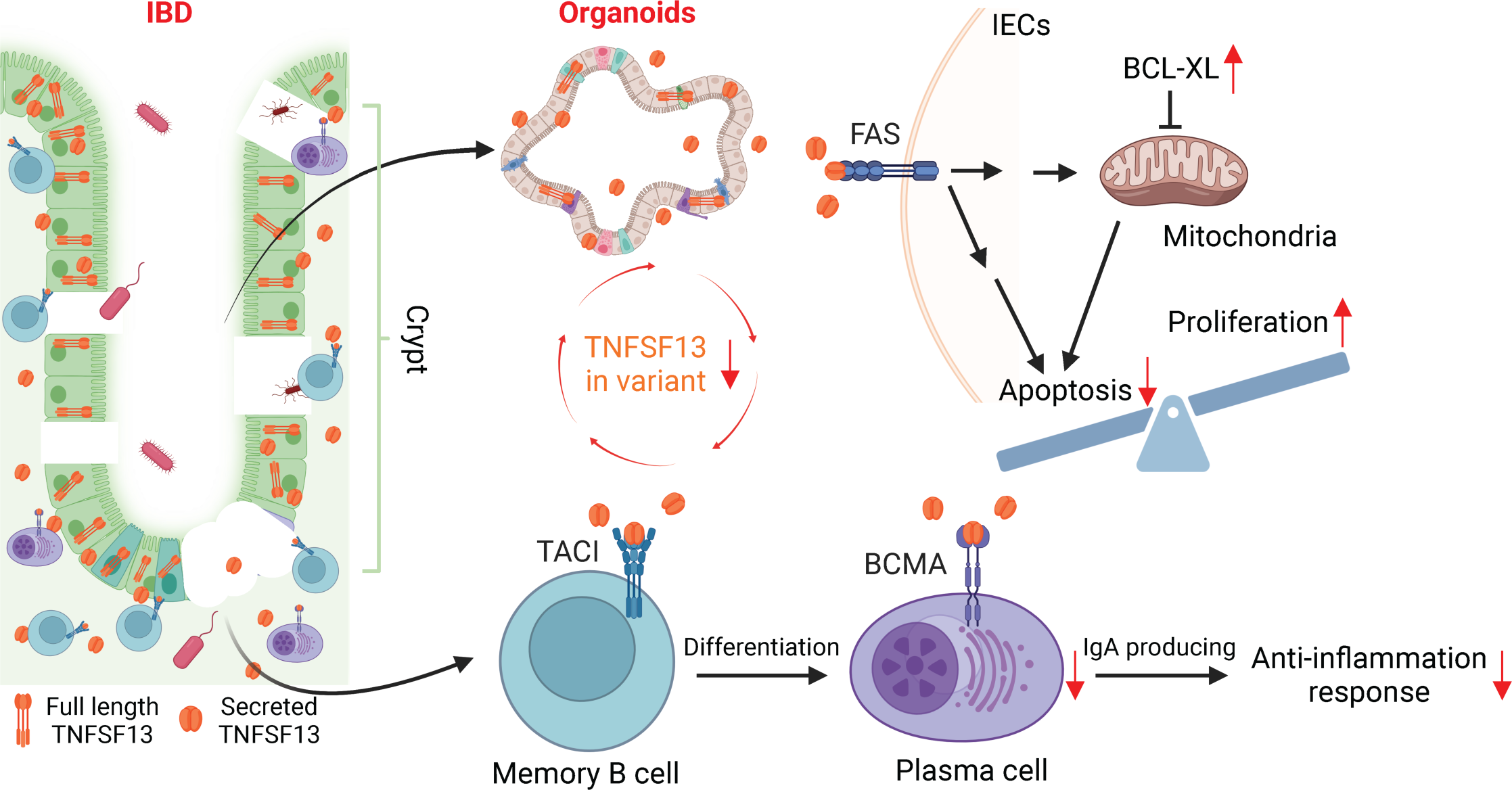

## INTRODUCTION

Inflammatory bowel disease (IBD) is attributed to a combination of factors including environment, diet, microbiota, and genetics(1). Very early onset (VEO)-IBD is a classification of pediatric IBD diagnosed in children who present with symptoms before age 6 (2, 3). Patients with VEO-IBD may exhibit more severe clinical symptoms, higher failure rates to conventional therapies, and strong family history as well as different genetic contributions to disease onset compared with older children or adults with IBD (4). The discovery and characterization of new gene variants in patients with VEO-IBD have improved our understanding not only of IBD pathogenesis, but also of fundamental intestinal biology. Reported genetic variants in patients with VEO-IBD are broadly characterized as immune, epithelial, or combined epithelial and immune in nature (5-7). The current mainstay of treatment for IBD, immunosuppressive therapy, is directed towards immune-mediated pathways, leaving an untapped opportunity for epithelial-targeted therapies (reviewed in (7)). The development of human organoid technology from affected patients’ epithelial stem and progenitor cells provides a translational model to study the physiology of intestinal epithelial cells in IBD (8). In this study, we used tissue-derived colonoids and induced pluripotent stem cell (iPSC)-derived colon organoids to investigate the function, mechanisms, and functional roles of epithelial Tumor Necrosis Factor Ligand Superfamily Member 13 (TNFSF13/APRIL), a cytokine typically attributed to B cell maturation and function.

TNFSF13 is secreted from myeloid cells and is best characterized for its effects on immune cells (9-11). Upon binding to its receptors TACI or BCMA, TNFSF13 promotes B cell activation, proliferation, maturation, plasma cell survival, and subsequent immunoglobin production, leading to activation of anti-inflammatory pathways (12-14). A recent study described a patient with common variable immunodeficiency harboring a homozygous frameshift mutation in *TNFSF13*, which resulted in the absence of plasmablasts and increased marginal zone B cells with a normal number of B cells in blood (15). Moreover, TNFSF13 deficiency in dendritic cells impairs differentiation from memory B cells to plasma cells *in vitro* (15, 16). Studies in mice suggest that TNFSF13 may have roles in intestinal epithelial cell-immune cell crosstalk. Epithelial cell-secreted TNFSF13 can promote immunoglobulin A2 (IgA2) class switching triggered by bacterial sensing via Toll-like receptors (9). A different mouse study found that overexpression of epithelial TNFSF13 resulted in enhanced anti-inflammatory B cell differentiation *in vitro* (17). Anti-inflammatory roles of TNFSF13 have also been reported in other tissues (18-20); however, the functional roles of TNFSF13 in human intestinal epithelial cells, and putative contribution to mucosal damage or healing, are not known. The goal of the present study was to evaluate the functional consequences of a novel *TNFSF13* gene variant using organoid models and tissue analyses to understand new, fundamental epithelial biology that may elucidate previously unknown mechanisms of disease pathogenesis.

## RESULTS

### A novel *TNFSF13* variant reduces TNFSF13 expression and alters epithelial proliferation

The current study emerged from a patient with severe colonic infantile onset IBD diagnosed at age 4 months (Supplementary Figure 1A, Table 1), with clinical history described in Methods. Whole exome sequencing (WES) was performed on the patient and his parents and identified a *de novo* heterozygous frameshift mutation (an inserted T in exon 3) in *TNFSF13* gene (NM_003808: c.372_373insT, pAla125_Thr126fs) in the patient (Supplementary Figure 1B). Repeat immune work up was performed and while his initial immune evaluation was unrevealing, due to refractory disease, repeat studies demonstrated increased transitional B cells consistent with impaired class switching. While other, predominantly homozygous TNFSF13 variants have been reported (https://mastermind.genomenon.com /detail?mutation=NC_000017.11:g.7559652A%3EG), our variant was not found in 1000 Genomes, ESP, ExAC or gnomAD sequence databases and no predictions were available from PolyPhen or SIFT.

Sanger sequencing confirmed the presence of the *TNFSF13* variant strand in peripheral blood mononuclear cells (PBMCs) and colonoids from the patient (Supplementary Figure 1B). qPCR with 3 different probes around variant *TNFSF13* indicated a significant decrease in *TNFSF13* mRNA compared with healthy controls and patients with VEO-IBD without an identified monogenic defect, defined hereon in as *TNFSF13* wild type VEO-IBD (just shown as VEO-IBD below) (Supplementary Figure 1C). This frameshift mutation caused a premature stop codon, leading to a predicted truncation in the protein via SWISS-MODEL (Supplementary Figure 1D). Typically, functional TNFSF13 is assembled into a homo- or hetero-trimer (21). Although it retained the intact transmembrane region and furin cleavage site, the truncated variant protein is predicted to lack most of its soluble region (Supplementary Figure 1D-F). ELISA confirmed a significant decrease in secreted TNFSF13 in variant colonoid media compared to healthy control and VEO-IBD colonoids (Figure 1A). RNAscope for *TNFSF13* in variant colonoids and colon biopsies demonstrated decrease in epithelial *TNFSF13* transcript levels (indicated by individual red dots) compared to controls (Figure1B & C, technical controls in Supplementary Figure 2A and B-C). Taken together, these data demonstrate a significant decrease of TNFSF13 on mRNA and protein levels in variant tissue.

**Figure 1.**
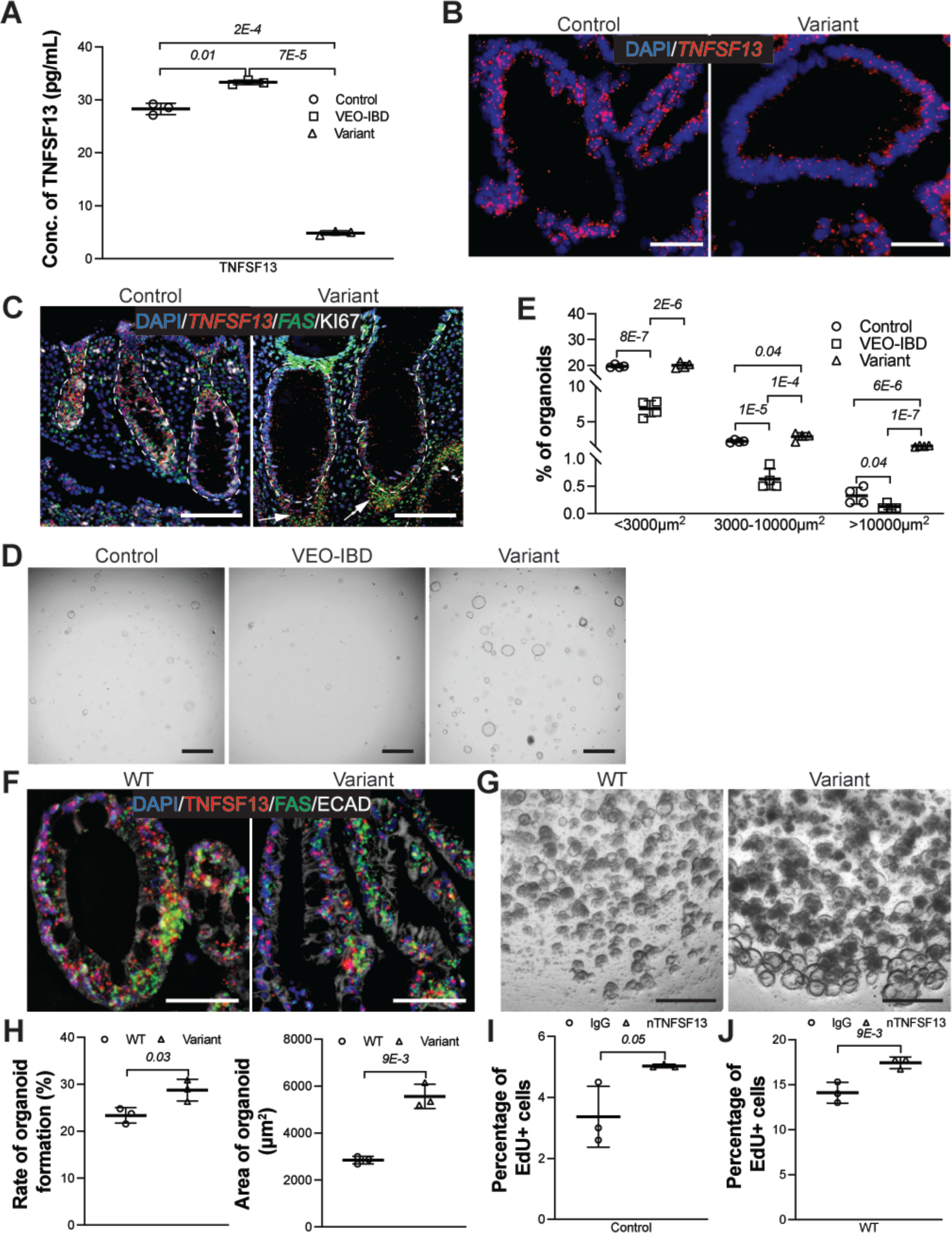
TNFSF13 variant colonoids/organoids exhibit enhanced colonoid formation efficiency and proliferation. **(A)** ELISA for secreted TNFSF13 in colonoid culture conditioned media (n=3 lines of colonoids from 3 different patients for Control and VEO-IBD, n=3 passages/batches of colonoids for Variant). **(B-C)** Representative immunostaining images for *TNFSF13* RNAscope probe in colonoids **(B)** and co-staining of *TNFSF13* and *FAS* RNAscope probes with Ki67 antibody in colon biopsies from control and variant subjects **(C)**. Arrowheads denote accumulated cells outside of epithelial crypts. Scale bar: 50 μm **(B)** and 100 μm **(C)**. n=3 lines of colonoids from 3 different patients for Control, n=3 passages/batches of colonoids for Variant in **(B)**; n=3 different patients for Control, n=3 slides from different blocks for Variant in **(C)**. **(D)** Representative images (left) of indicated samples for colonoid formation assays on d6 post seeding. Scale bar: 300 μm. **(E)** Quantification of total newly formed colonoids categorized by size on d6 post seeding. n=4 lines of colonoids from 4 patients for Control and VEO-IBD, n=4 passages/batches of colonoids for Variant. Each passage/batache has more than two statistic replicates. **(F)** Representative immunostaining images for co-staining of *TNFSF13* and *FAS* RNAscope probes with E-cadherin antibody in WT and variant iPSC-derived colon organoids on d7 post seeding. Scale bar: 50 μm. n=3 passages/batches of organoids. **(G)** Representative images for organoid formation assay on d9 post seeding in WT and variant iPSC-derived organoids. Scale bar: 400 μm. n=3 passages/batches of organoids. Each passage/batch has more than two statistic replicates. **(H)** Quantification of rate and area of newly formed organoids at 9d post seeding. Two-tailed Student’s *t*-test was used for statistical analysis. **(I-J)** Percentage of EdU^+^ cells after IgG or TNFSF13 neutralizing antibody (nTNFSF13) treatment for control tissue-derived colonoids **(I)** or WT iPSC-organoids **(J)** at d7 post seeding. Colonoid size was calculated by the maximum of vertical projection area. n=3 lines of colonoids from 3 different patients for **(I)**. n=3 passages/batches of organoids for **(J)**. *P* values are shown on bar graphs unless *P*>0.05. Two-way ANOVA (with multiple comparisons) was used for statistical analysis in **(A)** and **(E)**. Two-tailed Student *t*-test was used in **(H-J)**.

Upon visual inspection, we noticed an increase in colonoid number and size in *TNFSF13* variant versus control patient colonoids (healthy subjects and *TNFSF13* wild type VEO-IBD) at day 6 post seeding (Figure 1D-E, Supplementary Figure 2D). We directly assessed organoid formation efficiency via single cell plating and measured colonoid size as a proxy for proliferative capacity. Colonoids were significantly more numerous and larger in *TNFSF13* variant versus controls (Figure 1D-E). To confirm whether the observed colonoid formation efficiency and size phenotypes were driven by variant *TNFSF13* and not a consequence of the tissue state at the time of biopsy, we generated a human induced pluripotent stem cell (iPSC) line with the same variant and compared it to a wildtype (WT) *TNFSF13* isogenic control line. After directed differentiation into colon organoids(22), RNAscope, qPCR, and ELISA demonstrated the variant line had decreased *TNFSF13* compared to WT (Figure 1F and Supplementary Figure 2E-G). Furthermore, single cell-seeded organoid formation assays showed higher organoid formation efficiency and size in *TNFSF13* variant versus WT organoids (Figure 1G-H, Supplementary Figure 2H). Since our data demonstrated that epithelial TNFSF13 may have anti-proliferative roles in non-variant cells, we used a TNFSF13 neutralizing antibody on control colonoids and WT iPSC-derived colon organoids to evaluate proliferation directly using EdU incorporation. We confirmed the ability of the antibody to neutralize TNFSF13 using dose curves in mouse splenic B cell proliferation assays as published previously(23) (Supplementary Figure 3A-C). We observed an increase in EdU^+^ proliferative cells in both control patient colonoids and WT colon organoids treated with TNFSF13 neutralizing antibody compared to IgG control (Figure 1I-J). Taken together, TNFSF13 neutralization data are consistent with our observation that decreased TNFSF13 expression promotes increased organoid size as a result of increased proliferation.

### TNFSF13 binds to FAS receptor in colonic epithelial cells

TNFSF13 can bind to multiple surface receptors in different cell types(12-14). We investigated expression of these receptors in tissue-derived colonoids and iPSC-derived colon organoids. Flow cytometry (FACS) analysis revealed that TACI and BCMA, which are abundant in B cells and plasma cells, are not detected by FACS in colonic epithelial cells (Supplementary Figure 4A-B). Instead, FACS implicated the lesser known TNFSF13 receptors FAS and HVEM (24) were detected in colonoids and iPSC colon organoids (Figure 2A-B). Since FAS has been associated previously with proliferation and apoptosis (25), we next tested whether TNFSF13 can interact with the FAS receptor in colonic epithelial cells via co-immunoprecipitation (co-IP). We observed that FAS is only detected when the capture antibody for TNFSF13 is present, in contrast to IgG and input controls (Figure 2C). Furthermore, RNAscope data indicate expression of TNFSF13 and FAS in human colonoids, providing spatial evidence of their co-expression in the same and neighboring cells, including in Ki67^+^ proliferating cells and FABP2^+^ enterocytes (Figure 2D-F, white arrowheads). To confirm a functional role for FAS in control colonoids and WT iPSC colon organoids, we treated cultures with a FAS neutralizing antibody and observed a modest yet significant increase in EdU^+^ cells in response to FAS neutralization by FACS (Figure 2G). FAS is a TNF superfamily receptor commonly described as a pro-apoptotic factor; however, some studies demonstrate additional roles such as NFκB activation, among other roles (26). Our results demonstrate that FAS neutralization has a similar effect to TNFSF13 neutralization: to increase the proportion of proliferative cells in culture. Taken together, these data suggest that a TNFSF13-FAS axis can modulate epithelial proliferation *in vitro*.

**Figure 2.**
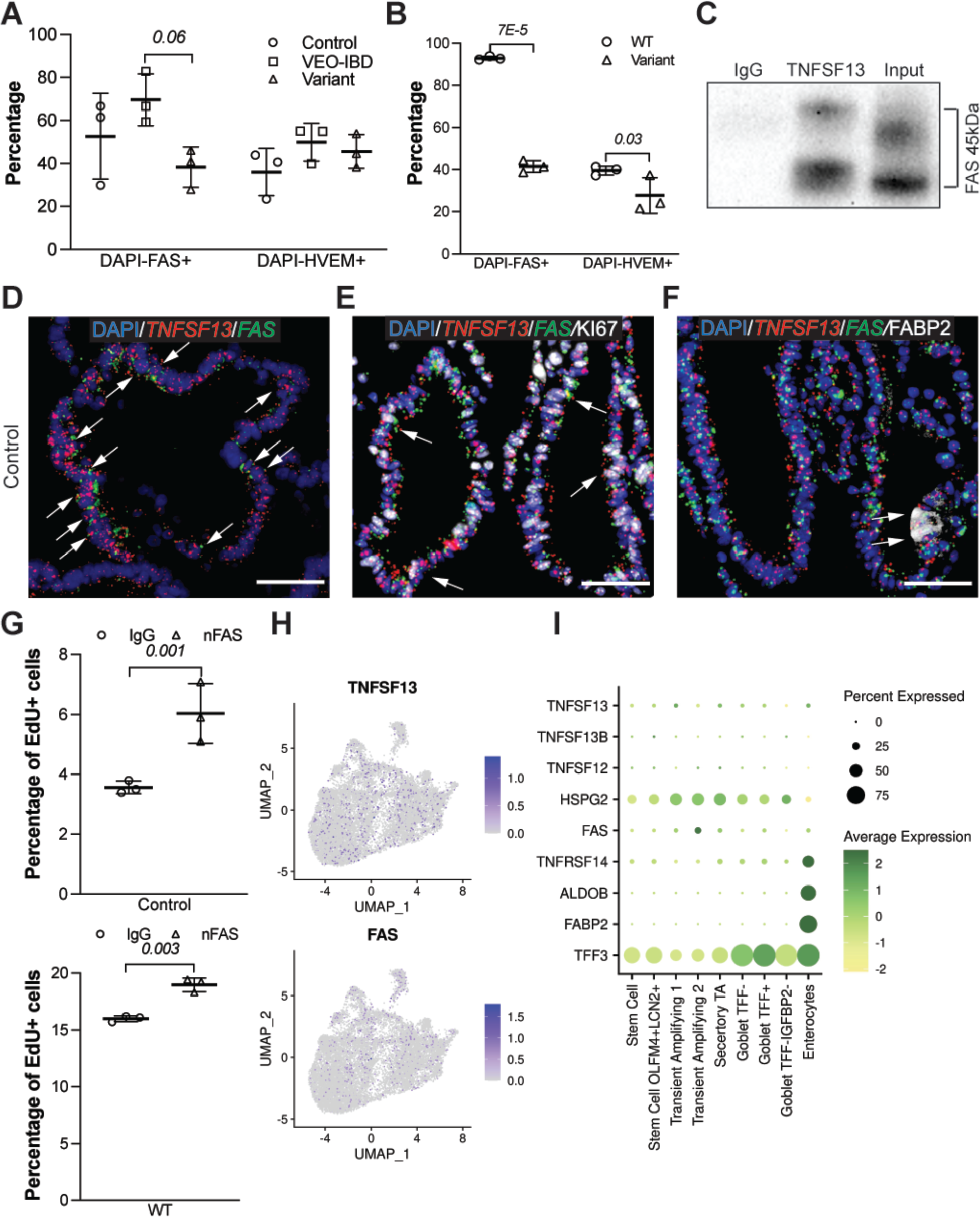
FAS is a receptor for TNFSF13 in colonic epithelial cells. **(A-B)** Percentage of FAS^+^ and HVEM^+^ cells in colonoids **(A)** or iPSC-organoids **(B)** on d7 post seeding by FACS. **(C)** Western blotting for FAS with co-IP supernatant from control colonoids d7 post seeding. TNFSF13 was used as capture antibody for co-IP. **(D)** Representative immunostaining images for co-staining of *TNFSF13* and *FAS* RNAscope probes in control colonoids on d7 post seeding. White arrow heads denote co-expression of *TNFSF13* and *FAS*. Scale bar: 100 μm. **(E)** Representative immunostaining images for co-staining of *TNFSF13* and *FAS* RNAscope probes with Ki67 antibody in control colonoids on d7 post seeding. White arrow heads denote co-expression of *TNFSF13, FAS,* and Ki67. Scale bar: 100 μm. **(F)** Representative immunostaining images for co-staining of *TNFSF13* and *FAS* RNAscope probes with FABP2 antibodies in control colonoids on d7 post seeding. White arrow heads denote co-expression of *TNFSF13*, *FAS,* and FABP2. Scale bar: 100 μm. **(G)** Percentage of EdU^+^ cells in IgG or FAS neutralizing antibody (nFAS)-treated control colonoids (upper) or WT iPSC-organoids (lower) on d7 post seeding. Two-tailed Student’s *t*-test was used for statistical analysis. **(H)** UMAP plots showing the expression pattern of *TNFSF13* and *FAS* in scRNA-seq data from human tissue-derived colonoids. n=2 lines of colonoids from 2 different patients for Control and VEO-IBD, n=2 passages/batches of colonoids for Variant. **(I)** Dot plot indicating the relative expression pattern of selected genes of *TNFSF13* family and related receptors and enterocyte markers among annotated clusters for human colonoids scRNA-seq data. *P* values are shown on bar graphs unless *P*>0.05. Two-way ANOVA (with multiple comparisons) was used for statistical analysis in **(A-B)**. Two-tailed Student *t*-test was used in **(G)**.

### Transcriptome analysis reveals altered apoptosis pathways in *TNFSF13* variant epithelium

We next evaluated transcriptional differences between control, *TNFSF13* wild type VEO-IBD and *TNFSF13* variant colonoids via single cell RNA sequencing (scRNA-Seq). Based on previously annotated marker genes (27), we identified and assigned colonic epithelial cells to 9 clusters: 2 stem cell clusters, 3 transit-amplifying (TA) cell clusters, 3 goblet cell clusters and 1 enterocyte cluster which were visualized by uniform manifold approximation and project (UMAP) (Figure 3A-B; cell counts in Supplementary Table 3). UMAP and dot plot of combined samples demonstrated broad expression of TNFSF13 and FAS in human colonoids, especially in TA cells and colonocytes (dark purple dots on UMAP, Figure 2H-I).

**Figure 3.**
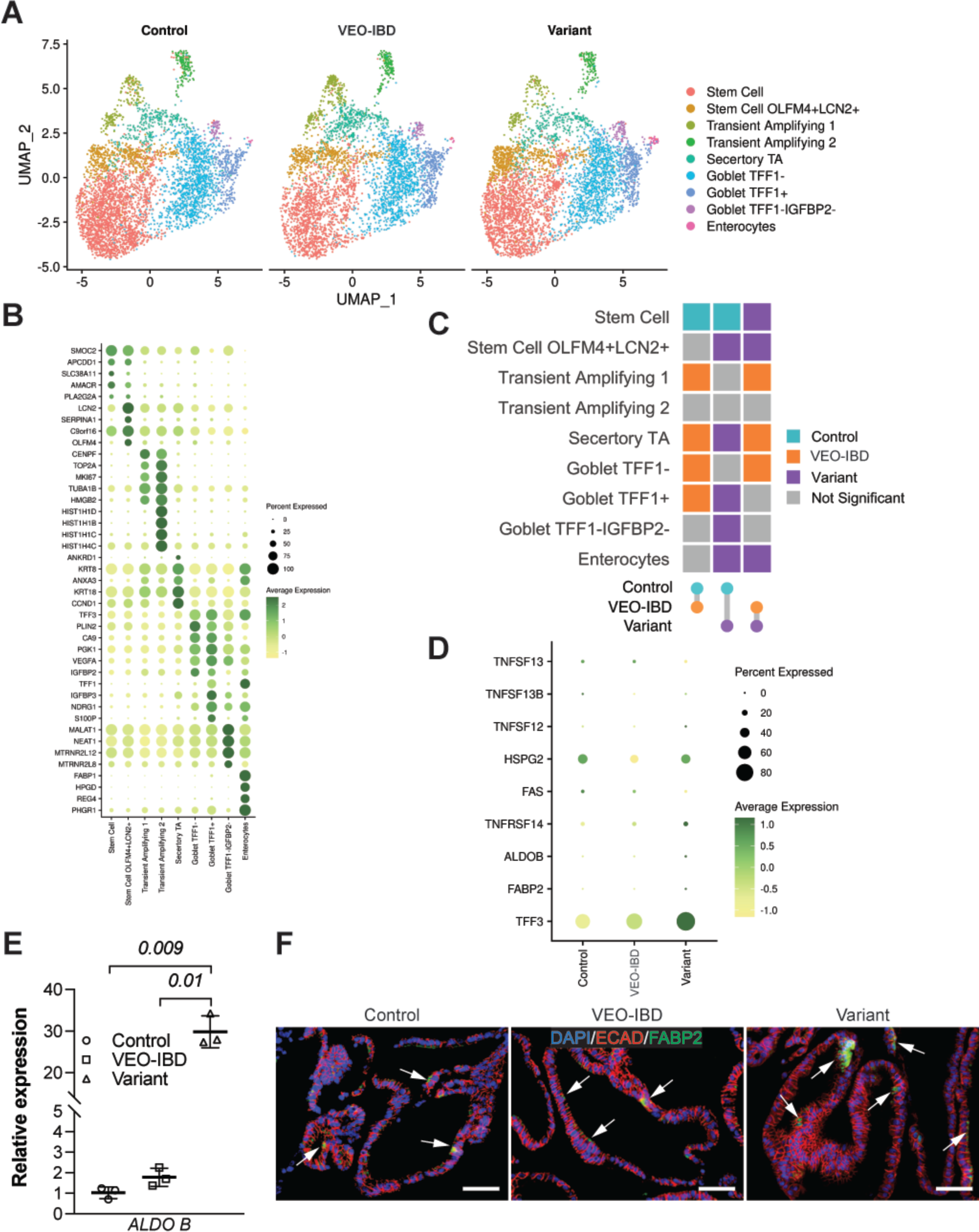
Transcriptomic profiling in human colonoids. **(A)** UMAP visualizations of scRNA-seq data for human colonoids. n=2 lines of colonoids from 2 different patients for Control and VEO-IBD, n=2 passages/batches of colonoids for Variant. **(B)** Dot plot with relative expression of top 5 changed genes for each annotated cluster for scRNAseq datasets in human colonoids. **(C)** Color scale indicates group with higher percentage of cells within a given cluster in each comparison. The color indicates the condition with higher percentage of a cluster in each pairwise comparison. **(D)** Dot plot with relative expression of selected genes of TNFSF13 family and related receptors and enterocyte markers among control, VEO-IBD and variant in human colonoids. **(E)** qPCR for ALDOB in colonoids on d7 post seeding. One-way ANOVA (with multiple comparisons) was used for statistical analysis. n=3 lines of colonoids from 3 different patients for Control and VEO-IBD, n=3 passages/batches of colonoids for Variant. **(F)** Representative IF images for FABP2 and E-cadherin in human colonoids. White arrow heads denoted FABP2^+^ cells. Scale bar: 50 μm. P values are shown on bar graphs unless P>0.05.

Overall, we observed moderate but likely inconsequential shifts in other cell type proportions when evaluating UMAPs of control, VEO-IBD and variant colonoids separately (Figure 3A-B) but with significantly increased expression of inflammatory marker *LCN2* and enterocytes (Figure 3C) in variant colonoids.

Overall, we observed moderate but likely inconsequential shifts in cell type proportions when evaluating UMAPs of control, TNFSF13 wild type VEO-IBD and TNFSF13 variant colonoids separately (Figure 3A-B). Evaluation of cell type proportions demonstrated that stem cells with expression of inflammatory marker LCN2 are significantly increased in TNFSF13 variant colonoids (denoted as “Stem Cell OLFM4^+^LCN2^+^”), as are goblet cells (denoted as “Goblet TFF1^+^” and “Goblet TFF1^-^IGFBP2^-^”) and colonocytes (Figure 3C). Likewise, gene expression analysis demonstrated increased expression of goblet cell marker *TFF3* and enterocyte markers *ALDOB* (28) and *FABP2* (29) in *TNFSF13* variant colonoids, albeit in a small percentage of cells (Figure 3D). We confirmed by qPCR increased expression of *ALDOB* (Figure 3E) and immunofluorescence (IF) staining for FABP2 (white arrows, Figure 3F) in *TNFSF13* variant colonoids compared with control and VEO-IBD colonoids. We also performed scRNA-seq on fresh biopsies from the same *TNFSF13* variant subject and an additional healthy control subject (Supplementary Table 3) and annotated clusters using published cell markers from human biopsies (30). Analysis of biopsy scRNA-seq data confirmed expression of *TNFSF13* and *FAS* in epithelial cells and a similar lack of robust differences in cell type proportions between *TNFSF13* variant and controls as seen in respective colonoid lines (Supplementary Figure 5A-E). We conclude that phenotypic differences between *TNFSF13* variant and controls are not due to significant changes in lineage allocation between groups.

To explore putative mechanisms of the TNFSF13-FAS axis in colonic epithelial cells, we performed Gene Ontology (GO) enrichment analysis of the combined scRNA-seq data (Supplementary Table 4). Consistent with phenotypic data, we observed an enrichment of pathways involved in epithelial cell proliferation and apoptosis in *TNFSF13* variant versus VEO-IBD or healthy controls (Supplementary Figure 6A, red arrowheads). Colonoid qPCR data confirmed the relative increased expression of proliferation-associated genes, *ID1* and *ECM1* (31, 32) (Figure 4A-B), and mitochondrial anti-apoptotic genes, *ACAA2* (33) and *BCL2L1* (34) (Figure 4C-D) in *TNFSF13* variant versus VEO-IBD or healthy controls. Immunostaining for apoptosis (TUNEL) demonstrated significantly fewer TUNEL^+^ cells and TUNEL^+^ FABP2^+^ cells in *TNFSF13* variant versus control colonoids (Figure 4E-F), which could explain the increase in enterocytes observed in *TNFSF13* variant colonoids in Supplementary Figure 5. Finally, immunoblot indicated increased BCL-XL (anti-apoptotic protein encoded by *BCL2L1*) (34) in both *TNFSF13* variant colonoids and iPSC colon organoids compared to respective controls (Figure 4H). We also observed FAS expression in T cells (Supplementary Figure 8C). Since there are accumulated FAS^+^ cells close to epithelial cells in variant tissue (Figure 1D, arrowhead), our findings provide an additional potential mechanism of action of mucosal TNFSF13 and FAS^+^ T cells that can be pursued in a follow-up study. Taken together, transcriptomics, histological analyses, and immunoblot data support the conclusion that TNFSF13 insufficiency both enhances proliferation and limits apoptosis in colonic epithelial cells, particularly colonocytes.

**Figure 4.**
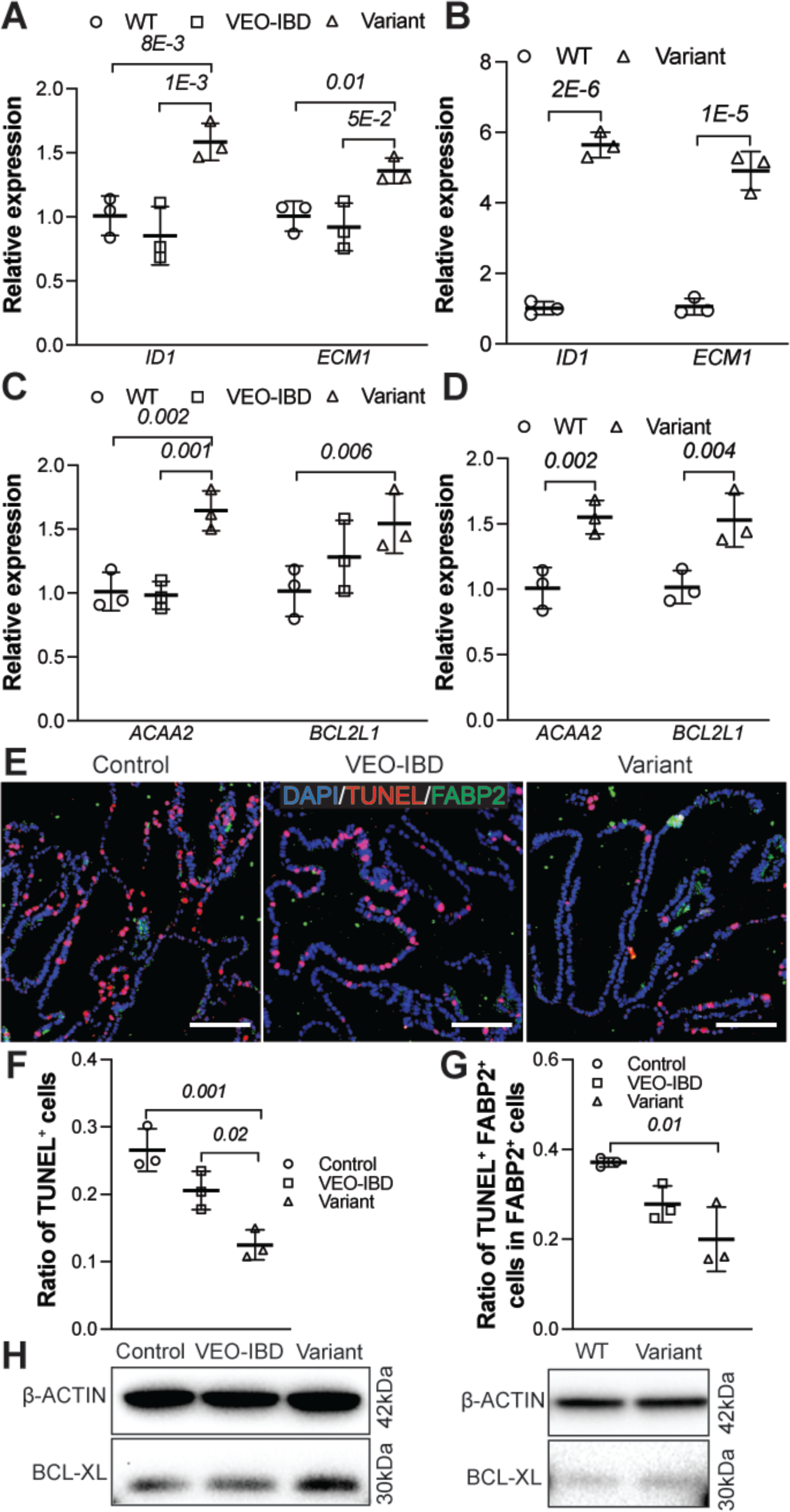
TNFSF13 augments the balance of apoptosis and proliferation through FAS-apoptosis pathway. **(A-B)** qPCR for *ID* and *ECM1* in colonoids **(A)** and iPSC-derived colon organoids **(B)**. **(C-D)** qPCR for *ACAA2* and *BCL2L1* in tissue-derived colonoids from control, VEO-IBD, variant **(C)** and iPSC organoids from WT and variant **(D)**. **(E-G)** Representative immunostaining images **(E)** for TUNEL and FABP2 in colonoids. Scale bar: 100 μm. Quantification for ratio of TUNEL^+^ cells **(F)** per colonoid and TUNEL^+^FABP2^+^ cells in FABP2^+^ cells **(G)**. One-way ANOVA (with multiple comparisons) was used for statistical analysis. **(H)** Western blotting for BCL-XL in colonoids (upper) and iPSC-organoids (lower). β-ACTIN was used as a loading control. *P* value shown in the bar graphs unless *P*>0.05. Two-way ANOVA (with multiple comparisons) was used for statistical analysis in **(A-D)**. n=3 lines of colonoids from 3 different patients for Control and VEO-IBD, n=3 passages/batches of colonoids for Variant. n=3 passages/batches of iPSC-organoids.

### Epithelial *TNFSF13* regulates tissue associated memory B cell differentiation

TNFSF13 is best characterized for its roles in regulating proliferation and differentiation in B cells and plasma cells. We therefore evaluated circulating and tissue immune populations in *TNFSF13* variant and control subjects. We first examined peripheral blood immune changes using FACS of PBMCs (Supplementary Figure 7A). We observed an increase in CD19^+^ B cells in *TNFSF13* variant blood compared with healthy controls, but not as much as compared to *TNFSF13* wild type VEO-IBD (Supplementary Figure 7A-B). CD19^+^CD27^+^CD38^+^ plasmablasts, IgD^+^ and IgM^+^ plasmablasts were relatively lower in *TNFSF13* variant compared with healthy control and VEO-IBD (Supplementary Figure 7B). There were no significant differences in IgD^+^ or IgM^+^ switch, memory, naïve or transitional B cells, or other immune cells (T cell, nature killer cell and monocyte) in *TNFSF13* variant compared with healthy control and VEO-IBD PBMCs (Supplementary Figure 7B-C). Taken together, there were moderate, but non-significant differences in peripheral B cells in *TNFSF13* variant versus control subjects.

The immune cells within the intestinal mucosa play an essential role in the establishment and regulation of intestinal inflammation and injury in IBD (35). We therefore explored immune changes in colonic tissue of *TNFSF13* variant and control subjects. We first evaluated 6,014 variant and 4,755 control cells in the scRNA-seq data of lamina propria layer from the same biopsies as described above, which were sub-clustered into 6 subsets (Supplementary Figure 8A-C). We further sub-clustered B cell (germinal center B cells-GC B cells, memory B cells and naïve B cells) and plasma cell (7 clusters based on Ig types-IgA, IgK, IgL, IgG, and NFKBIA signature) populations (36) (Figure 5A). Cell type abundance analysis indicated fewer germinal center B cells and naïve B cells, but more memory B cells in *TNFSF13* variant compared to control biopsies (Figure 5B-C). These findings are consistent with prior reports of increased memory B cell recruitment and differentiation to plasma cells under inflammatory conditions (37). For plasma cells, although the population of IgA^+^IgK^+^NFκBIA^-^ and IgA^+^IgL^+^NFκBIA^-^ cells are relatively increased, total IgA+ PCs (∼69.3%) decreased in *TNFSF13* variant compared with control (∼74.9%) (Figure 5C). In contrast, we noticed IgG^+^ plasma cells were relatively increased, which had been reported to contribute to IBD (37).

**Figure 5.**
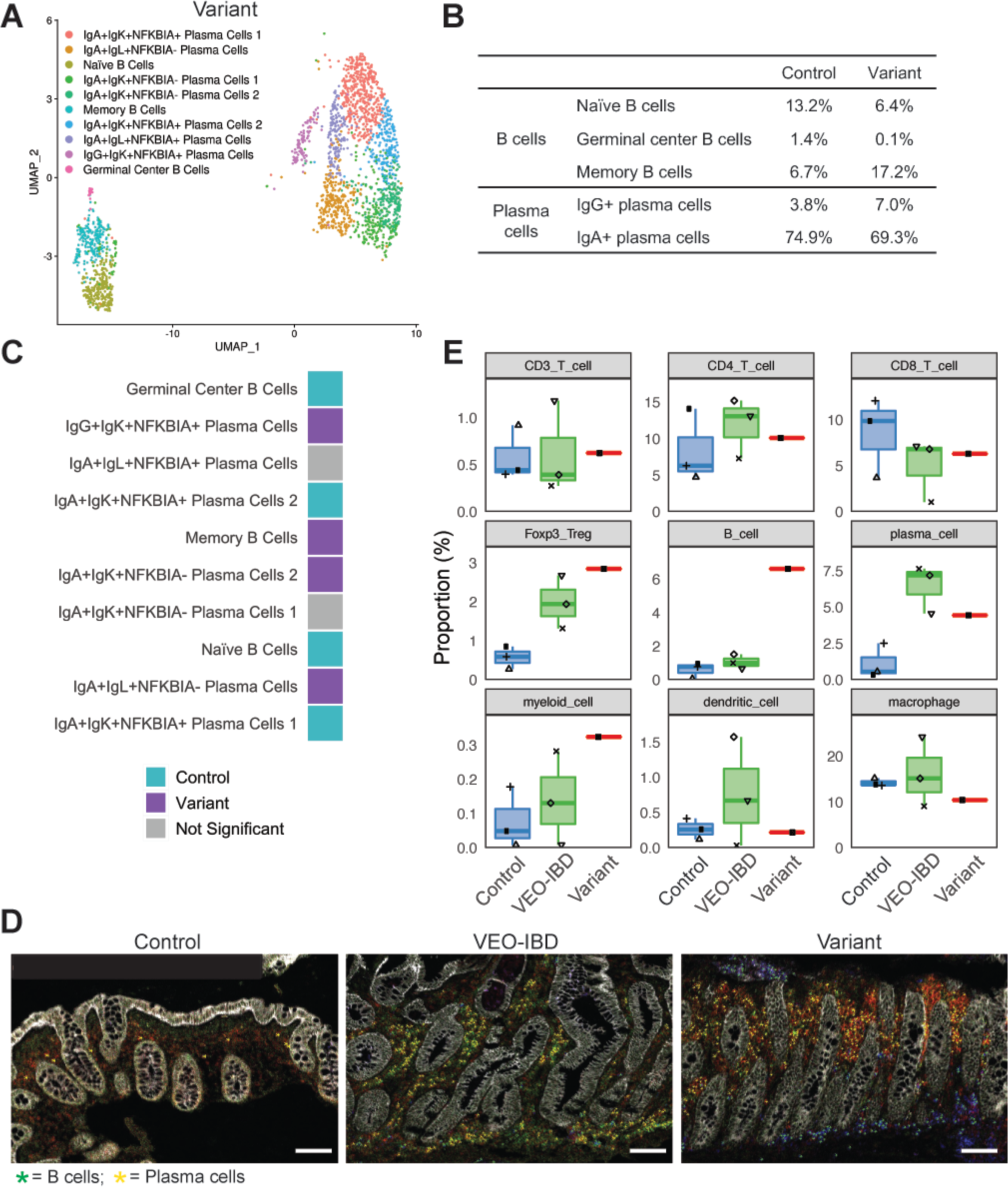
Increased abundance of memory B cells and depletion of IgA^+^ plasma cells observed in *TNFSF13* variant colon. **(A)** UMAP visualizations of scRNA-seq data for B cell and plasma cell clusters among lamina propria cells from variant colon biopsies. n=1 patient for Control and Variant. **(B)** Table indicates abundance (%) of B cell and PC subsets in control and variant samples from scRNAseq data from Variant and Control colon biopsies. **(C)** Comparison of cell type abundance between samples from scRNAseq data from Variant and Control colon biopsies. Color scale indicates which group has a higher percentage of cells within a given cluster. **(D)** Representative IMC overlay images of epithelial, B cell and plasma cell markers in colon from control, VEO-IBD and variant patient. Scale bar: 100 μm. Marker for B cell: CD20^+^; Markers for plasma cell: CD20^-^CD27^+^CD38^+^. n=3 different patients for Control and VEO-IBD, n=3 slides from different blocks for Variant. **(E)** Boxplot showing the rate of immune cell composition quantified by calculating the proportion of specific markers in all cells at the same region (both lamina propria and epithelial cell populations). n=3 different patients for Control and VEO-IBD, n=3 slides from different blocks for Variant.

Imaging mass cytometry (IMC) is a multiplexed imaging platform that utilizes antibodies conjugated to heavy metal isotopes, permitting quantification of different cell types within local tissue niches (38). IMC identified 9 major immune cell populations within colon sections from 7 patients (3 controls, 3 TNFSF13 wild type VEO-IBD, and 1 *TNFSF13* variant with 2 different biopsies)): CD3+ T cells, CD4+ T cells (T helper cells), CD8+ T cells (cytotoxic T cells), FOXP3+ regulatory T cells (Tregs), B cells, PCs, myeloid cells, dendritic cells, and macrophages (Figure 5D, Supplementary Figure 9A-B and Table 5). Because IMC retains the X and Y coordinates of each cell in each image, we were able to assess immune cell composition with spatial resolution. We found increased CD20^+^ total B cells near epithelial crypts in *TNFSF13* variant compared to control and VEO-IBD tissue (Figure 5D-E, green stars). Plasma cell numbers in *TNFSF13* variant were lower than VEO-IBD, but higher than controls. Given that TNFSF13 promotes proliferation and differentiation of B and plasma cells (9), we evaluated cell abundance of proliferative B cell and plasma cell combined with Ki67 staining for proliferation as a putative explanation for aggregation of B cells in *TNFSF13* variant tissue. The percentage of Ki67^+^ total B cells in *TNFSF13* variant sections was lower than control and *TNFSF13* wild type VEO-IBD sections, suggesting that accumulated B cells in *TNFSF13* variant tissue is likely due to enhanced recruitment rather than increased B cell proliferation (Supplementary Figure 9C). Similarly, the percentage of Ki67+ plasma cells in *TNFSF13* variant tissue was lower than in VEO-IBD tissue as well. Taken together, scRNA-seq and IMC data suggest that decreased TNFSF13 in variant tissue might reduce differentiation of memory B cells to IgA producing plasma cells and may contribute to accumulation of B cells in close proximity to epithelial crypts. These newly described phenotypes in *TNFSF13* variant tissue may separately contribute to mucosal damage via (1) reduced beneficial epithelial-IgA+ plasma cell interactions (39), and (2) aberrant B cell accumulation in the epithelial compartment, which hinders stromal contributions to mucosal healing(40).

### Co-culture studies confirm epithelial TNFSF13-mediated B cell modulation

Prior studies reported TNFSF13 defects in dendritic cells or monocyte-derived dendritic cells differentiated from iPSCs impaired memory B cell proliferation and differentiation to plasmablasts, and then to plasma cells (15, 16). To investigate the function of epithelial secreted TNFSF13 on memory B cell differentiation, we developed a series of methods to either directly co-culture of human colonoids with memory B cells, or human colonoid conditioned media with memory B cells (Figure 6A). Co-culture of sorted human memory B cells (DAPI^-^CD19^+^CD20^+^CD27^+^) with equal numbers of human control, VEO-IBD, and variant colonoids at day 8 post seeding indicated the percentage of differentiated plasmablasts significantly decreased in variant conditions (Figure 6B, Supplementary Figure 10A-D). Since mixing colonoid and B cell media at a ratio of 1:1 only permitted short-term co-culture, we collected 2-day conditioned media from equal numbers of human control, VEO-IBD, and variant colonoids and mixed it with human B cell media (ratio of 1:1) for treatment of cultured B cells. Consistent with co-culture studies, we found the percentage of plasmablasts differentiated from sorted human memory B cell significantly decreased in variant at day 8 post seeding with colonoid conditioned media and B cell media mixture (Figure 6c, Supplementary Figure 10a-f). We also found a reduction in the percentage of plasma cell that differentiated from plasamblasts at day 14 post seeding in variant media-treated B cell cultures (Figure 6D). He *et al* showed that epithelial-derived TNFSF13 promotes IgA class switching in mice (9). We therefore examined the IgA^+^ population in total plasma cells and found a decrease in the percentage of IgA^+^ plasma cells in variant media conditions (Fig 6E). ELISA for IgA in media at day 14 post-seeding confirmed that secreted IgA was decreased in B cells with variant conditioned media compared with control and VEO-IBD (Figure 6F).

**Figure 6.**
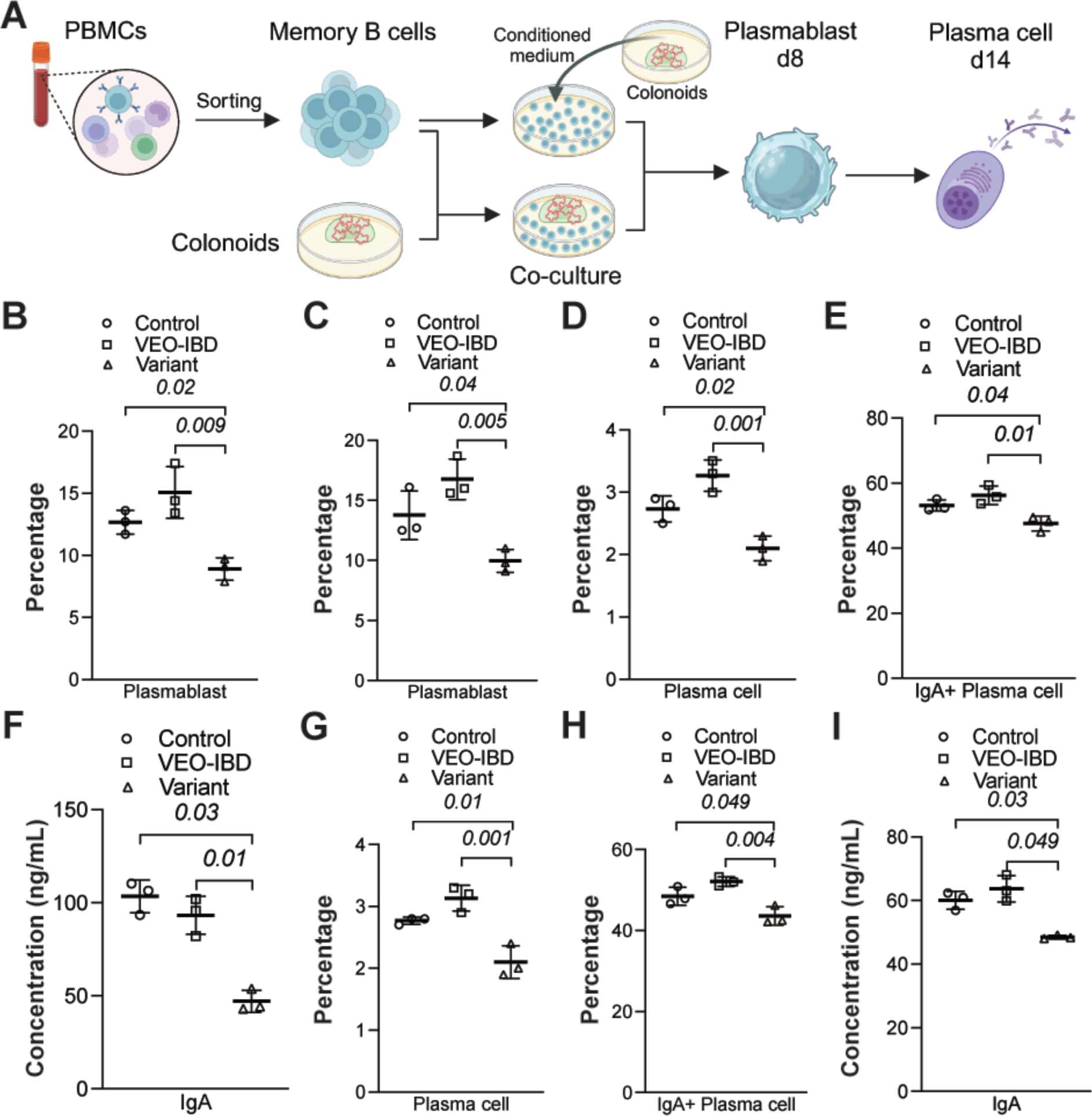
Epithelial-secreted TNFSF13 modulates differentiation of memory B cells to plasmablasts and plasma cells. **(A)** Schematic of co-culture model. **(B)** Percentage of plasmablasts differentiated from sorted human memory B cells at day 8 post-seeding via co-culturing with equal numbers of control, VEO-IBD, and variant colonoids in an IntestiCult media and B cell media mixture (ratio 1:1). **(C)** Percentage of plasmablasts differentiated from sorted human memory B cells at day 8 post-seeding via culturing in conditioned media consisting of B cell media and conditioned media (ratio 1:1). **(D-E)** Percentage of plasma cells and IgA^+^ plasma cells differentiated from sorted human memory B cells at day 14 post-seeding with B cell media-conditioned media (ratio 1:1). **(F)** ELISA for IgA in media differentiated from sorted human memory B cells at day 14 post-seeding. **(G-H)** Percentage of plasma cells and IgA^+^ plasma cells differentiated from sorted human memory B cells at day 14 post-seeding. **(I)** ELISA for IgA in media from sorted human memory B cells at day 14-post seeding by culturing in conditioned media mixture starting at day 6 post-seeding. *P* value shown in the bar graphs unless *P*>0.05. One-way ANOVA (with multiple comparisons) or two-tailed Student’s *t*-test was used for statistical analysis. n=3 lines of colonoids from 3 different patients for Control and VEO-IBD, n=3 passages/batches of colonoids for Variant. n=3 passages/batches of iPSC-organoids. n=7 independent donors to obtain human memory B cells.

To further investigate the role of TNFSF13 on differentiation of plasmablasts to plasma cells, we maintained memory B cells in B cell media for 6 days to get an equal percentage of plasmablasts and then changed to conditioned media mixture from control, VEO-IBD, variant at day 6 post seeding. FACS and ELISA data at day 14 post-seeding indicate the percentage of both plasma cells and IgA^+^ plasma cells decreased in variant media conditions, which is consistent with differentiation directly from memory B cells (Figure 6G-I, Supplementary Figure 10G). Given the importance of IgA^+^ memory B cells and IgA-producing plasma cells for immune homeostasis in the gastrointestinal tract(39), our data suggest that a decrease in epithelial-secreted TNFSF13 may promote reduced antibody secretion and other anti-inflammatory responses, ultimately contributing to IBD pathogenesis(6).

## DISCUSSION

Local cues from intestinal epithelial cells can shape the functional specificity of immune responses, and understanding new mechanisms of epithelial-immune interactions is critical for designing new, epithelial-targeted treatments for IBD. We used reductionist organoid culture systems to dissect novel epithelial cell roles for TNFSF13 in epithelial and immune cells of the human colon. We demonstrated that TNFSF13 secreted by colonic epithelial cells can act upon epithelial cells themselves to increase proliferation and decrease apoptosis *in vitro*. We further identified FAS as the putative TNFSF13 receptor expressed in epithelial cells that can modulate apoptosis signaling either directly through caspase-8-mediated proteolysis of effector caspases (i.e., caspases-3 and caspase-7) or through mitochondria-released apoptosome to propagate the caspase activation cascade (41). These findings are particularly intriguing as dysregulated cell death can be an important feature in patients with IBD.

When broadening our analyses to tissue biopsies, we found dysregulation of local immune responses in the colon, particularly affecting B cell and plasma cell populations in *TNFSF13* variant tissue as reported previously (9, 10, 15). Prior studies demonstrated that TNFSF13 is secreted by intestinal epithelial cells upon Toll-like receptor-mediated bacterial sensing, leading to IgA(2) class switching. These findings support a dual-hit model whereby (1) depleted epithelial TNFSF13-FAS axis signaling promotes an imbalance in proliferation and apoptosis leading to aberrant wound healing and (2) depleted epithelial-derived TNFSF13 leads to insufficient antibody production in response to antigen exposure. While we do not provide data in direct support of the latter, our scRNA-seq and imaging mass cytometry data demonstrate that TNFSF13 deficiency is associated with decreased plasma cells and increased B cell accumulation in the colonic lamina propria. These findings suggest that while the patient’s total quantitative immunoglobulins were normal, that under tissue stress, mucosal changes in TNFSF13 could lead to impaired tissue immunoglobulin levels. This is intriguing, since a recent report in mice demonstrated that B cell depletion is beneficial for mucosal healing in experimental colitis (40). This same study identified a robust expansion of an IFN-induced population of B cells marked by CD274 and Ly6a during the mucosal healing phase following experimental colitis in mice. We evaluated our scRNA-seq data for CD274 and additional reported marker genes from this study but did not find an analogous cell cluster in our human biopsy data (not shown).

Patients with antibody defects can develop IBD, and the contribution of B cells to IBD pathogenesis has been a long-standing area of interest in the field (reviewed in (42)). Recently, aberrant mucosal B cell diversity and maturation were reported via scRNA-seq analysis in patients with ulcerative colitis(37). As such, we decided to pursue epithelial B cell interactions via 3D co-culture. While 2D co-culture systems, including with epithelial cells and B cells, have been reported previously, co-culture in 3D-culture systems in human primary cells are only recently emerging(43-45). Co-culture of human primary B cells with 3D epithelial colonoids has not been reported. In this study, we developed a human primary memory B cell and colonoid co-culture system to explore secreted epithelial TNFSF13 and memory B cell maturation. This system not only provides an accessible method to study signaling between epithelial and immune compartments *in vitro,* but also provides a powerful tool to explore dual tissue compartment changes associated with genetic variants in patients. While TNFSF13 has been described previously as a regulator of B cell maturation, our model permitted this paradigm to be tested directly in primary human cells. Optimization of this co-culture model will allow for future studies to further evaluate epithelial B cell interactions, including reciprocal effects of B cells on epithelial cells or with the addition of other immune of stromal cell types, as has been reported in mice (40).

Our study has limitations. It is tempting to speculate that phenotypes observed in TNFSF13 variant colonoids are due to a generalized disease state rather than variant specific. Furthermore, retention of *in vivo* inflammatory components during colonoid establishment could hinder epithelial cell growth and be a confounding variable. To mitigate this concern, we evaluated all colonoids samples between passages 6 to 15. Additionally, our generation and evaluation of *TNFSF13* variant and isogenic control iPSC-derived colon organoids provided orthogonal data in support of epithelial TNFSF13 functions that have not been described previously. A separate limitation of our study is that TNFSF13 deletion has been reported previously in both human and in mice, where intestinal or IBD-like symptoms are not prominent (15, 46). Our study therefore suggests that TNFSF13 insufficiency may contribute to, rather than cause, IBD. Alternatively, our specific variant could have unknown functions leading to the colonic manifestations observed in the patient. As such, biochemical studies of our variant and analyses in mouse models will be needed to disentangle dose-dependent effects of TNFSF13 and direct links to the complex clinical presentation observed in the present patient.

While data presented herein are based upon a single patient with variant *TNFSF13*, there are broader implications of our findings. A recent study demonstrated that recombinant TNFSF13 can restore plasmablast differentiation *in vitro* in a dendritic cell B cell co-culture experiment using cells from a patient with common variable immunodeficiency harboring a different *TNFSF13* variant (15). It is therefore possible that restoration of TNFSF13 could be a tractable therapy for patients with other *TNFSF13* variants. It is also possible that given the mechanism of disease, patients with variants in *TNFSF13* may not have a sustained response to conventional IBD therapies. In addition, our finding that *TNFSF13* variant colonic epithelial cells exhibit increased proliferation and decreased apoptosis may indicate the need for earlier and more frequent caner surveilance, as this cellular phenotype likely increases risk of colorectal cancer even more than already exists for patients with IBD. In summary, our findings demonstrate novel roles for TNFSF13 to modulate the balance of proliferation and apoptosis in colonic epithelial cells. An imbalance of proliferation and apoptosis, together with aberrant mucosal B cell dynamics observed in the present study, underscores the importance of identifying mechanisms converging upon epithelial and immune compartments that could serve as future therapeutic targets.

## MATERIALS AND METHODS

### Sex as a biological variable

Both male and female patients were included in the study.

### Subject enrollment and demographics

This study was conducted with the approval of the Institutional Review Board (IRB): IRB # 14-010826. All parents of patients provided written informed consent. Biopsy specimens, human peripheral blood mononuclear cells (PBMCs), and histological samples were obtained from de-identified patient. The patient with the TNSF13 variant presented with colonic IBD at 4 months of age with diarrhea and poor growth. Due to medically refractory disease, the patient underwent diverting ileostomy at 21 months of age and over time developed progressive stricturing disease requiring subsequent hemi-colectomy with sparing of the right colon. The patient developed sacroilitis post-operatively and ultimately achieved remission of intestinal and joint disease with infliximab and rapamycin. Immunologic workup was performed at the patient’s initial presentation that was unrevealing, including lymphocyte subsets, immunoglobulins, DHR and FOXP3 analysis. As part of the research protocol, biosamples were obtained from patients with VEO-IBD and healthy controls. Patients with VEO-IBD were diagnosed at ≤6 years old of age using standard methods of endoscopic, radiologic, laboratory and clinical evaluation. Indications for colonoscopy in patients with VEO-IBD included diagnosis, change in disease status, and surveillance of disease. All patients with VEO-IBD underwent immunologic and genetic evaluation. Control VEO-IBD subjects denoted as *TNFSF13* wild type do not have a known or candidate monogenic disorder. Genetic studies were carried out through whole exome sequencing (WES) and included trio analyses. Healthy control samples were selected from subjects undergoing colonoscopy for the following reasons: abdominal pain, poor growth, rectal bleeding, or diarrhea, and had normal endoscopic and histologic findings. Individuals with a previous diagnosis of other intestinal or systemic inflammatory disease, including chronic allergic or inflammatory diseases, were excluded. Detailed patient information and the purpose for each sample used in this study can be found in Supplementary Table 1.

### Whole exome sequencing, variant calling, and annotation

Whole exome sequencing was performed on the variant patient and his parents (Supplementary Table 1). Library preparation and exome capture were performed using the Agilent SureSelect v4 capture kit with DNA samples isolated from PBMCs. Sequence reads were aligned to the reference human genome (GRCh37) using the Burrows–Wheeler alignment (BWA) algorithm (v.0.7.15)(47) and variants were called using GATK’s best practices. Variants were functionally annotated with information from multiple databases including dbSNP(48), dbNSFP(49), 1000 Genomes Project(50), and the Genome Aggregation Database (gnomAD v2.1.1)(51) using SnpEff (http://snpeff.sourceforge.net) then filtered to retain only moderate- and high-effect, rare (minor allele frequency < 1%) variants. Trio analysis for the patient included identifying variants that are *de novo*, compound heterozygous, homozygous, and X-linked and results were limited to variants within genes known to be associated with VEO-IBD or immunodeficiency.

### Generation of colonoids from patient biopsies

Mucosal biopsies were obtained from endoscopically affected and unaffected areas of the terminal ileum and/or left/right colon during colonoscopy procedures conducted for disease surveillance or diagnosis (Supplementary Table1). The collected samples were promptly transported in cold DMEM (Corning, New York, USA) on ice to the laboratory for subsequent crypt isolation. To generate patient-derived colonoids, biopsies were rinsed once in cold DPBS (Corning, New York, USA) and then in chelation buffer.

Subsequently, they were incubated in a cold, fresh 0.5M EDTA chelation buffer comprising 2% sorbitol (Fisher Scientific, Massachusetts, USA), 1% sucrose (Sigma-Aldrich, Massachusetts, USA), 1% BSA (Fisher Scientific, Massachusetts, USA), and 1x Gentamicin (Thermo Fisher Scientific, Massachusetts, USA) in DPBS for a duration of 30 minutes. Following incubation, the biopsy samples were gently scraped off using forceps to release crypts. The isolated crypts were then resuspended in 30-50 µL of Matrigel (Corning, New York, USA) after filtrating with 70μm nylon strainer, with the exact volume adjusted based on the number of crypts and plated to achieve an optimal density. A successful isolation was defined by the presence of at least 50 crypt units per biopsy region, and plating volumes of Matrigel were meticulously adjusted to ensure uniform crypt density and minimize potential density-related growth bias. Colonoids were fed every other day in human IntestiCult Organoid Growth Media (Stem Cell Technologies, British Columbia, Canada) and collected or split roughly 7 days after plating depending on the specific experiment. When colonoids reached an appropriate density to avoid overgrowth, mechanical passaging was initiated as outlined below: colonoids were suspended in 3mL cold advanced DMEM/F12 (Thermo Fisher Scientific, Massachusetts, USA) and then pipetted up and down for 5 times in 15 mL conical tube using a p1000 µL pipette tip fitted with p200 µL pipette tip. Subsequently, the collected colonoids were spun down, resuspended, and re-plated in the Matrigel at an appropriate volume to maintain uniform density. 10 μM Y-27632 (LC Laboratories, Massachusetts, USA) was added for the first 2 days. Media was changed every other day. All colonoid lines in this study were utilized between Passage 6 and Passage 15 for consistency and reliability.

### Colonoids/organoids formation assay

For colonoids/organoids formation assay, cells were collected at day 7 after plating and digested with 0.05% trypsin (ThermoFisher Scientific, Massachusetts, USA) for 10 min at 37°C in a bead bath (final ratio as 10% FBS was added to de-activate trypsin). Single cells were further dissociated by pipetting up and down several times. 5,000 live cells (or 1,000 live cells for tissue-derived colonoids) were quantified by a Countess™ 3 FL Automated Cell Counter (ThermoFisher Scientific, Massachusetts, USA) and plated in 10 µL Matrigel per well in a 96-well plate with 100 µL of media. 10 μM Y-27632 was added for the first 2 (tissue-derived colonoids) or all (iPSC-derived colon organoids) days. Imaging and quantification of live colonoids/colon organoids were performed on days 1-7 to monitor plating efficiency and growth using the Keyence BZ-X 700 all-in-one microscope with accompanying analysis software.

### Generation of human induced pluripotent stem cell (iPSC)-derived organoids

Human iPSC lines carrying the same heterozygous frameshift mutation in TNFSF13 as human variant patient (NM_003808: c.372_373insT, pAla125_Thr126fs), and wildtype iPSC line were generated and validated by the Children’s Hospital of Philadelphia Human Pluripotent Stem Cell Core using the parental line CHOPWT14 described previously (52). Feeder-independent culture, expansion, and differentiation of both the wild type (WT) and variant iPSC lines were performed at the University of Pennsylvania (UPenn) iPSC Core. To prepare cells for differentiation, iPSCs were seeded on 0.12 mg/ml Geltrex™ LDEV-free reduced growth factor basement membrane matrix (ThermoFisher Scientific, Massachusetts, USA, 1:100 diluted in cold DMEM/F12 from the same vendor) pre-coated plates (1 hour at RT) and maintained with mTeSR™1 complete medium at 37°C with 5% O_2_ and CO_2_ incubator. Human iPSC-derived organoids were differentiated and maintained as previously described protocol with the following adjustments(22).

Approximately 5 million live single cells (per well of a 6-well plate) were seeded on Corning Matrigel (diluted 1:30 in cold DMEM/F12) coated plate after digesting with Gentle Cell Dissociation Reagent for 10min at 37°C (with the addition of 10 μM Y-27632 for the first 10 days). The STEMdiff™ Definitive Endoderm Kit (Stem Cell Technologies, British Columbia, Canada) was used to differentiate monolayer cultures to definitive endoderm (DE) once cells reached around 90-100% confluency on Day 1 or Day 2 after plating. Following a 4-day DE differentiation with sequential administration of supplements MR and CJ according to the manufacturer’s instructions, the cells was transitioned to hindgut endoderm (HE) differentiation by treating with HE media (containing 3 μM CHIR99021-Cayman Chemical Company, 1x GlutaMAX-Thermo Fisher Scientific, 1x Pen/Strep-Thermo Fisher Scientific, 0.5 μg/mL FGF4-PeproTech, and 1x B27 supplement-Thermo Fisher Scientific in RPMI 1640-Corning) for another 4 days (fresh HE media was changed daily). At the end of HE differentiation, cells were primed for colonic differentiation over a 12-day period in colonic medium comprised of advanced DMEM/F12 with 3 μM CHIR99021, 1x GlutaMAX, 1x Pen/Strep, 0.3 μM LDN193189-Cayman Chemical Company, 1× B27 supplement and 0.1 μg/mL EGF-R&D Systems. Colonic media was refreshed daily, and detached spheroids were collected and re-seeded in Corning Matrigel simultaneously with colonic media. Differentiated colonic cells were disassociated into single cells by Accutase for 10 min at 37°C and then seeded in Matrigel with colonic media (with the addition of 10 μM Y-27632 for the first 2 days). iPSC-derived organoids were obtained from both detached spheroids and seeded colonic cells. Organoids were fed every other day and split every 7 days. To remove other types of cells, single cell suspension from organoids, digested by 0.05% trypsin at 37°C for 10 min (10% FBS-Peak Serum as final ratio to de-activated trypsin), was subjected to flow cytometry (FACS) with MoFlo Astrios sorter (Beckman Coulter, Pennsylvania, USA) or FACSAria Fusion Sorter (BD Biosciences, New Jersey, USA) in the CHOP Flow Core after incubating with PE anti-CD326 (EpCAM) Monoclonal Antibody (G8.8) (ThermoFisher Scientific, Massachusetts, USA) and DAPI for 30min on ice in dark. Sorted DAPI^-^EpCAM^+^ intestinal cells were seeded and expanded in Matrigel (∼50,000 live cells per 30uL Matrigel; 10 μM Y-27632 was added for the first 10 days and then split as normal) to generate organoids for subsequent analysis and experiments.

### Mouse B cell isolation/maintenance, treatment, and resazurin assay

To isolate mouse B cells, normal mouse spleen was harvested from adult WT mice and processed into single cell suspension by mincing it through a 70 µm cell strainer placed in a 6-well plate containing 5 mL of 1x DPBS, using the flat end of a plunger from a 3-cc syringe. Mouse experiments were approved under IACUC protocol 001278 at the Children’s Hospital of Philadelphia. The pellet was collected by centrifugation at 300x g at 4°C and incubated in 5 mL of 1x RBC lysis buffer (Thermo Fisher Scientific, Massachusetts, USA) per spleen for 5 min at room temperature with occasional shaking to remove blood cells. The reaction was stopped by adding 20 mL of 1x DPBS. Subsequently, the cells were collected and incubated with FITC anti-mouse CD19 [6D5] (Biolegend, California, USA) diluted at 1:50 in FACS buffer for 30 min on ice in the dark. DAPI at a final concentration of 0.1 µg/mL was add for an additional 10 min. After washing with 3 mL FACS buffer and resuspending in 1mL FACS buffer, the cells were sorted with a MoFlo Astrios sorter (Beckman Coulter, Pennsylvania, USA) or FACSAria Fusion Sorter (BD Biosciences, New Jersey, USA) in CHOP Flow Core. Specifically, 100,000 sorted DAPI-CD19+ B cells per well were plated in a 96-well plate with 100 μL RPMI 1640 with L-Glu media (Corning, 10%FBS+1xAnti-Anti+10mM HEPES--Thermo Fisher Scientific + 50uM β-Mercaptoethanol--Thermo Fisher Scientific) at 37°C/5%CO_2_. To stimulate B cells, 10µg/mL of F(ab’)2-Goat anti-Mouse IgM (mu) Secondary Antibody (Cat# 16-5092-85, Thermo Scientific, Massachusetts, USA) was added to the media upon plating. To evaluate the function of TNFSF13 in B cells, recombinant Human TNFSF13 (HEK293-expressed) protein (Cat #5860-AP-010, R&D Systems, Minnesota, USA) at a varying concentration gradient (0, 0.0005, 0.001, 0.005, 0.01, 0.05, 0.1, 0.2, 0.5, 1, 1.5, 2 μg/mL) was added at 4h post plating. Meanwhile, to assess the efficiency of neutralizing antibody, nTNFSF13 (Cat #MAB5860, R&D Systems) with a concentration gradient (0, 0.005, 0.01, 0.05, 0.1, 0.5, 1, 5, 10, 20, 30, 50 μg/mL) was added along with 500 ng/mL recombinant Human TNFSF13 protein for each well at 4h post plating. For measuring proliferation in B cells, Resazurin (Cat# AR002, R&D Systems) was added to all wells at a volume equal to 10% of the cell culture volume (10 μL) and incubated for another 4 hours in incubator. Fluorescence was measured using a TECAN Infinite® 200Pro microplate reader (TECAN, Zürich, Switzerland) with a wavelength of 544 nm excitation and 590 nm emission. Negative control wells containing only media were set up in parallel to account for background effects. The process and analysis of data was conducted using an online tool (https://www.aatbio.com/tools/four-parameter-logistic-4pl-curve-regression-online-calculator) and the equation form was created as shown in Supplementary Figure 2B-C.

### Human memory B cell isolation/differentiation, treatment, and analysis by flow cytometry

Human PBMCs from 7 independent donors were purchased from UPenn Human immunology Core (supported by NIH P30 AI045008 and P30 CA016520). To isolate memory B cells, human PBMC cell pellets were collected by centrifuging at 300xg for 5min at 4°C, and then incubated in FACS buffer (2% FBS in DPBS) with indicated antibodies in the dark on ice for 30 min: anti-CD19 Mouse Monoclonal Antibody (PE) [clone: SJ25C1] (BioLegend), anti-CD27 Mouse Monoclonal Antibody (FITC) [clone: M-T271] (BioLegend), BD Pharmingen™ APC-H7 Mouse Anti-Human CD20 (BD Biosciences), Anti-CD20 Mouse Monoclonal Antibody (PE/Cy7®) [clone: 2H7] (BioLegend), anti-human IgA Antibody (APC) (Miltenyi Biotec, Cologne, Germany). DAPI (Sigma-Aldrich, Massachusetts, USA) was added at a final concentration of 0.1µg/mL for an additional 10min. After washing with 3 mL FACS buffer, cells were resuspended and subjected to flow cytometry using a MoFlo Astrios (Beckman Coulter) or FACSAria Fusion Sorter (BD Biosciences) in the CHOP Flow Core. Following sorting, 37,000-100,000 sorted DAPI^-^CD19^+^CD20^+^CD27^+^ memory B cells were seeded equally in a 96-well plate with 150μL B cell medium, consisting of RPMI 1640 with L-Glu media +10%FBS+1xAnti-Anti+10mM HEPES + 1µg/mL R848 (Cat# tlrl-r848, Invivogen, California, USA) + 50uM β-Mercaptoethanol at 37°C/5%CO2, IntestiCult/B cell media mixture, or conditioned media mixture, respectively. Media was changed every other day.

For co-culture of memory B cell with human colonoids, 3,000 clumps of control, VEO-IBD, variant colonnoids were seeded in 45μL Matrigel with 500μL human IntestiCult Organoid Growth Media for the first 2 days (considered colonoid seeding day as d-2). After 2 days of growth, an equal number of sorted DAPI^-^ CD19^+^CD20^+^CD27^+^ memory B cells were seeded in colonoid wells with 500μL mixture media (IntestiCult media : B cell media = 1:1) (considered B cell seeding day as d0). Mixture media was changed every other day. At d8, differentiated memory B cells were collected without disturbing the Matrigel dome and subjected to do FACS to examine the percentage of plasmablasts. Differentiated memory B cells were collected by centrifugation at 300xg for 5min at 4°C, and then incubated in FACS buffer (2% FBS in DPBS) with various antibodies for 30 min in the dark on ice: anti-CD27 Mouse Monoclonal Antibody (FITC) [clone: M-T271] (BioLegend), BD Pharmingen™ APC Mouse Anti-Human CD38 (BD Biosciences), anti-CD138 Mouse Monoclonal Antibody (PE) [clone: MI15] (BioLegend). DAPI (Sigma-Aldrich, Massachusetts, USA) added at a final concentration of 0.1µg/mL for an additional 10min. After washing with 3 mL FACS buffer, cells were resuspended and analyzed with an LSR Fortessa analyzer (BD Biosciences).

For culturing memory B cells with conditioned media mixture, conditioned media was initially obtained by seeding 3,000 clumps of control, VEO-IBD, and variant colonnoids in 45μL Matrigel with 500μL human IntestiCult Organoid Growth Media. Fresh conditioned media was then collected from the 2-day conditioned human IntestiCult Organoid Growth Media in human colonoids wells every other day (d2, d4, d6, d8, d10), respectively. Moreover, the remaining conditioned media collected at d4-post seeding was centrifuged and stored for ELISA to test the expression level of secreted TNFSF13. To create conditioned media mixture, collected fresh conditioned media was centrifuged at 300xg for 5 min at 4°C to remove debris and then mixed with an equal volume of fresh B cell media. An equal number of sorted DAPI^-^ CD19^+^CD20^+^CD27^+^ memory B cells were seeded in the fresh conditioned mixture media (considered d0) and were fed every other day. To examine the percentage of plasma cells and IgA^+^ plasma cells, differentiated memory B cells were collected at d14 by centrifugation at 300xg for 5min at 4°C, and then incubated in FACS buffer (2% FBS in DPBS) with various antibodies for 30 min in the dark on ice: anti-CD19 Mouse Monoclonal Antibody (PE) [clone: SJ25C1] (BioLegend), anti-CD27 Mouse Monoclonal Antibody (FITC) [clone: M-T271] (BioLegend), Anti-CD138 Mouse Monoclonal Antibody (Brilliant Violet® 605) [clone: MI15] (BioLegend), anti-human IgA Antibody (APC) (Miltenyi Biotec). DAPI (Sigma-Aldrich) at a final concentration of 0.1µg/mL for an additional 10min. After washing with 3 mL FACS buffer, cells were resuspended and analyzed with an LSR Fortessa analyzer (BD Biosciences). Before FACS (d14 post-seeding), media in each well was collected and stored at -80°C after centrifugation for testing the expression level of IgA with ELISA.

## Data availability

Single cell RNA sequencing data will be deposited on a publicly available database. All other data are available from the corresponding author upon request.

The transcript profiling data is deposited in U.S. National Library of Medicine Gene Expression Omnibus (GEO) with accession number GSE243445.

Values for all data points in graphs are reported in the Supporting Data Values file.

**Additional materials and methods are provided in the online Supplementary Materials and Methods.**

## Supporting information

supplemental Files combined with figures and tables

## ACKNOWLEDGMENTS

We thank Drs. Ning Li, Yuhua Tian, Xin Wang, Christopher Lengner for protocols and reagents. We thank Drs. Katrina Estep, Wenli Yang, Feikun Yang, Jean Ann Maguire, Chintan Jobaliya, Youngjun Choi, Ricardo Cruz-Acuña for technical advice regarding iPSC generation, maintenance, and differentiation. We thank Dr. Kai Tan and lab members Anusha Thadi, Chia-hui Chen, Qin Zhu, Changya Chen, and Dr. Melanie Ruffner and lab member Jarad Beers for sharing equipment, reagents, and resources for performing of 10x library preparation. We gratefully thank Drs. Hui Wang, Rui Li, Zhen Lu for assistance with human and mouse B cell isolation and maintenance. We acknowledge support of the following cores: Penn NIH/NIDDK P30 Molecular Pathology and Imaging Core founded by Center for Molecular Studies in Digestive and Liver Diseases (P30DK050306), UPenn Pluripotent Stem Cell Core, UPenn Human Immunology Core, CHOP Flow Core, CHOP Human Pluripotent Stem Cell Core, CHOP Pathology Core, CHOP Center for Applied Genomics. This work was supported by the following: NIH R01 DK124369 (KEH), Lisa Dean Moseley Stem Cell Grant (KEH), Children’s Hospital of Philadelphia (CHOP) Institutional Development Funds and Gastrointestinal Epithelium Modeling Program (KEH), CHOP Roberts Collaborative Pilot Grant (ND), and NIH R01DK127044 (JRK).

## AUTHOR CONTRIBUTIONS

Dr. Kathryn Hamilton had full access to all of the data in the study and takes responsibility for the integrity of the data and the accuracy of the data analysis. K.E.H., J.R.K. conceptualized and supervised the study. K.E.H., J.R.K., X.M. designed and conducted all the experiments and manuscript drafting. J.R.K., K.E.S., K.H.K., M.D. and D.A.P. provided critical review of the manuscript for important intellectual content. Bioinformation analysis was performed by N.D. and X.M.. J.R.K., R.S., M.C., D.A.P., M.D. were involved in patient inclusion and sample acquisition. A.K. and X.M. were performed IMC staining and data analysis. T.K., P.A.W., L.R.P., L.A.S., C.H.D. were helped with protocol optimizing and material support. All authors were contributed to data acquisition, analysis and/or interpretation.

## ABBREVIATIONS

BCMA: B cell maturation antigen
CD: Crohn’s disease
co-IP: co-immunoprecipitation
DEGs: differentiated expression genes
DSS: dextran sulfate sodium
EdU: 5-ethynyl-2′-deoxyuridine
FACS: Flow cytometry
FAS: Fas cell surface death receptor
hiPSC: human induced pluripotent stem cell
HVEM: herpes virus entry mediator
IBD: inflammatory bowel disease
IEC: intestinal epithelial cell
IgA: immunoglobulin A
IMC: imaging mass cytometry
PBMC: peripheral blood mononuclear cell
PC: plasma cell
IF: immunofluorescence
IRB: Institutional Review Board
PBMCs: peripheral blood mononuclear cells
scRNAseq: Single cell RNA sequencing
TA: transient amplifying
TACI: transmembrane activator and calcium modulator and cyclophilin ligand interactor
TNFSF13: tumor necrosis factor ligand superfamily member 13
UC: ulcerative colitis
UMAP: uniform manifold approximation and project
VEO-IBD: very early onset inflammatory bowel disease
WES: whole exome sequencing
WT: wild type

