## supplemental Files combined with figures and tables for "TNFSF13 insufficiency disrupts human colonic epithelial cell-mediated B cell differentiation"

#### Supplementary Materials and Methods

1. Supplementary Figures

2. Supplementary Methods

3. Supplementary Methods References

4. Supplementary Figures and Figure legends

5. Supplementary Tables

#### SUPPLEMENTARY MATERIALS AND METHODS

##### <u>*SUPPLEMENTARY METHODS*</u>

*TOPO TA clone and sanger sequencing.* TOPO® TA Cloning® Kits for Sequencing (Thermo Fisher Scientific) was employed to validate the haplotype of both variant and control sequences according to the manufacture's protocol. Briefly, mRNA isolated from human PBMCs, colonoids for variant and control, as well as iPSC-derived organoids from variant and WT lines, was reverse transcribed to generate cDNA to serve as the template for TA cloning, following the protocol in above kit. PCR product was generated using a 50 µL PCR reaction mixture comprised of DNA Template 10-100 ng, 10X PCR Buffer 5 µL, 50 mM dNTPs 0.5 µL, and water to a final volume of 49 µL, Taq Polymerase (1 unit/µL) 1 µL (Cat# M0273S, New England Biolabs, Massachusetts, USA), specific primers (~200 ng each, Genewiz, New Jersey, USA) 1 µM for each with standard cycling parameters in thermal cycler ProFlex™ 3 x 32-well PCR System (95 ºC 5min, 95 ºC 30s, 56 ºC 30s, 72 ºC 40s, 35 cycles, 72 ºC 10min, 4 ºC) (Supplementary Table 2). The resulting PCR product was assessed using agarose gel electrophoresis.

For the TOPO® Cloning reaction, a mixture containing fresh PCR product (4 µL), Salt Solution (1 µL), water (to a final volume of 5 µL), and TOPO® vector (1 µL) was prepared and incubated at room temperature for 5 min. Subsequently, 2 µL of the TOPO® Cloning reaction and 1 vial of One Shot® chemically competent E. coli was gently mixed and incubated on ice for 5 min. This mixture was then subjected to a 30-second heat-shock at 42 ºC, followed by immediate transfer to ice. Pre-warmed LB Broth (Miller) medium (Sigma-Aldrich) was added, and the mixture was horizontally shaken at 37 ºC for 1 hour. Following the incubation, 50 µL of the mixture was spread onto a pre-warmed selective plate (containing LB Agar Broth with 50 µg/mL ampicillin-- Thermo Fisher Scientific) and the plate was incubated overnight at 37 ºC. 10 colonies for each line were picked and cultured overnight in 5 mL of LB Broth containing 50 µg/mL ampicillin. The plasmid DNA was extracted using QIAprep Spin Miniprep Kit (Qiagen, Hilden, Germany) according to the manufacture's protocol and submitted to GENEWIZ for Sanger Sequencing with the universal primer M13 forward and M13 reverse. The sequencing data was analyzed using the Benchling online tool (https://www.benchling.com/) to determine and confirm the haplotype of the variant and control sequences.

*Histological analyses and immunostaining.* Human colonoids or organoids were collected and stored at 70% ethanol after being fixed in 4% paraformaldehyde (PFA, VWR) in 1x PBS for 24 hours at 4°C and then submitted to the Molecular Pathology and Imaging Core (MPIC) of UPenn to create histological samples/unstained slides. To simultaneously detect the expression of RNAscope probes and antibodies, RNAscope® Multiplex Fluorescent v2 Assay (Cat# 323100) was employed, combined with Immunofluorescence Integrated Co-Detection kit (Advanced Cell Diagnostics, California, USA) according to the manufacture's protocol. Briefly, after baking for 1 hour at 60°C, the air-dried fresh sections were deparaffinized (2x 5min fresh xylene, 2x 2min 100% ethanol at room temperature, in turn) and thoroughly dried in an oven for 5 min at 60°C. The slides were then applied hydrogen peroxide for 10 min at room temperature. Following this, the slides were washed twice with distilled water before slowly immersing the slide rack into mild-boiled 1x co-detection target retrieval solution for 15 minutes

(98-102°C). At the end of target retrieval, the hot slide rack was promptly transferred to distilled water to wash twice, followed by once with 1x PBST (0.1% Tween-20). The slides were then incubated overnight at 4°C with primary antibody diluted in co-detection antibody diluent: anti-Ki67 antibody (SP6) (1:100, Abcam), anti-E-Cadherin antibody (1:50, BD Biosciences), anti-FABP2/I-FABP antibody (1:100, R&D systems). Following primary antibody incubation and washing 3x 2min with PBST, the slides were applied post-primary fixation by submerge in 10% Neutral Buffered Formalin (NBF, VWR, Pennsylvania, USA) for 30 min at room temperature in a fume hood. Whereat, protease plus was applied for 30 min at 40°C in a HybEZ™ Oven, followed by two 2-minute washes with distilled water. The slides were then subjected to hybridization with 1:50 diluted *TNFSF13* (Cat# 406981-C2) in *FAS* (Cat# 427031) (or diluted in probe diluent if using only the *TNFSF13* probe), positive control (Cat# 320881) and negative control (Cat# 320871) probes for 2 hours at 40°C in a HybEZ™ Oven, respectively. After probe hybridization, the slides could be either stored in 5x SSC (Promega) overnight at room temperature or for next step after being rinsed with 1x wash buffer for 2 x 2 min. Subsequently, the slides were hybridized with a series of steps: AMP1 for 30 min, AMP2 for 30 min, AMP3 for 15 min, HRP-C1 signal for 15 min (for *FAS* probe), 1:750 diluted Opal™ 570 (Cat# FP1488001KT, Akoya Biosciences, Massachusetts, USA) fluorophore in RNAscope® Multiplex TSA buffer for 30 min, HRP blocker for 15 min, HRP-C2 signal for 15 min (for *TNFSF13* probe), 1:750 diluted Opal™ 690 (Cat# FP1497001KT, Akoya Biosciences) fluorophore in RNAscope® Multiplex TSA buffer for 30 min, HRP blocker for 15 min in a HybEZ™ Oven at 40°C, successively. Washes with 1x wash buffer twice for 2 min at room temperature were performed between hybridization steps. After the final wash step, the slides were applied for specific fluorophore-conjugated secondary antibodies (AffiniPure IgG, Jackson ImmunoResearch, Pennsylvania, USA) from the same host as primary antibodies, diluted at 1:500 in co-detection antibody diluent, for 1 hour at room temperature. Following this, the slides were washed with 1x PBST 3x 2min at room temperature and then counterstained with DAPI for 30s without washing and mounted with Prolong Gold antifade mounting medium (Thermo Fisher Scientific). The slides could be stored at 4°C in the dark for up to two weeks.

Click-iT™ Plus TUNEL Assay for In Situ Apoptosis Detection, Alexa Fluor™ 594 dye (Cat# 10618, Thermo Fisher Scientific) was employed to identify apoptosis on slides according to the manufacture's protocol. Briefly, after baking for 1 hour at 60°C, tissue sections were deparaffinized/hydrated firstly (2x 5min fresh xylene, 2x 5min 100% ethanol, 5 min 95% ethanol, 5 min 80% ethanol, 5 min 70% ethanol, 5min di $H_2O$ at room temperature, in turn) and then applied for antigen retrieval for 15min in 10 mM citric acid in di $H_2O$ (pH 6.0) using a 2100 antigen retriever (Aptum Biologics, Southampton, UK). After rinsing once for 5 min with di$H_2O$ and 1xPBS respectively, the slides were fixed with 4% paraformaldehyde for 15 minutes at 37°C, permeabilized with Proteinase K solution at room temperature and fixed with another 4% paraformaldehyde for 5 minutes at 37°C, in turn. Between these steps, the slides underwent two 5-minute washes in PBS. In the final stages of the procedure, the slides were rinsed with deionized water, followed by a pretreatment step with 100 µL of TdT Reaction Buffer for 10 minutes at 37°C. Subsequently, the slides were incubated in 50 µL TdT reaction mixture (47 µL TdT Reaction Buffer, 1 µL EdUTP, 2 µL TdT enzyme) for 60 minutes at

37°C. Following being washed with deionized water, 3% BSA and 0.1% Triton™ X-100 (Sigma-Aldrich) in PBS, as well as 1x PBS for 5 min each, the slides were exposed to 50 μL of the Click-iT™ Plus TUNEL reaction cocktail (comprising with 45 μL Click-iT™ Plus TUNEL Supermix, 5 μL 10X Click-iT™ Plus TUNEL Reaction Buffer Additive) for 30 minutes at 37°C, protected from light. Following this, the slides were washed with 3% BSA in PBS and then 1x PBS for 5 minutes each. Moving forward, the slides were blocked with 3% BSA in PBS for 1 hour at room temperature protected from light before be subjected to an overnight incubation at 4°C with primary antibody (FABP2/I-FABP) diluted in blocking buffer. After being washed twice with 3% BSA in PBS, the slides were treated with fluorophore-conjugated secondary antibody diluted at 1:500 in blocking buffer for 1 hour and then counterstained with DAPI for 8 min at room temperature, protected from light. The slides finally were mounted with Prolong Gold antifade mounting medium and stored at 4°C in the dark.

All images were acquired eight hours later with Keyence BZ-X 800 all-in-one microscope and were further analyzed using software from the Keyence microscope, Adobe Photoshop, or Image J software. For quantification of copies of *TNFSF13* and *FAS* per cell in colonoids/orgnoids and colon tissue, the total number of red/green dots (representing positive signal) in each view were divided by total numbers of cells in the same view. At least 10 colonoids/organoids/crypts from 4-6 images for each sample were analyzed. For quantification of TUNEL$^+$ cells, more than 10 colonoids/views from 4-6 images were assessed for each sample. Detailed antibodies information was listed in Supplementary Table 2.

*5-ethynyl-2'-deoxyuridine (EdU) assay.* For EdU assay, appropriate colonoids, organoids and monolayers were pre-treated with 10 μM EdU in the respective media at day 7 (day 2 for monolayer) after plating. Following 2-hour EdU treatment, cells were collected and enzymatically digested to single cells, as described above. After DAPI staining and subsequently washing with 3 mL of 1% BSA in PBS, cell pellet was collected and processed for EdU staining using Click-iT™ Plus EdU Alexa Fluor™ 647 Flow Cytometry Assay Kit (Thermo Fisher Scientific) according to the manufacture's protocol. Briefly, the pellet was fixed with 100 μL of Click-iT™ fixative (Component D) for 15 minutes at room temperature, protected from light. After being washed with 3 mL of 1% BSA in PBS at the end of fixation, cells were then permeabilized in 100 μL of 1X Click-iT™ saponin-based permeabilization and wash reagent for additional 15 min in darkness on ice. Next, 500 μL of Click-iT™ Plus reaction cocktail (438 μL DPBS, 10 μL Copper protectant, 2.5 μL Fluorescent dye picolyl azide, 50 μL Reaction buffer additive for one sample) was added in cells without washing and incubated for 30 minutes at room temperature, protected from light. After being washed once with 3 mL of 1X Click-iT™ saponin-based permeabilization and wash reagent and resuspended in FACS buffer, the cells are ready to be analyzed with an LSR Fortessa analyzer (BD Biosciences) in the CHOP Flow Core.

*Co-immunoprecipitation (co-IP) and Western blotting.* Human colonoids cells at day 7 after plating were collected as shown above and lysed in 1 mL of NP-40 lysis buffer (Thermo Fisher Scientific) for co-IP, or RIPA lysis buffer (Thermo Fisher Scientific) for regular protein samples, along with protease and phosphatase inhibitors for 30 minutes on ice with shaking at 10-minute intervals. The resultant supernatant was then transferred to a fresh tube placed on ice, following centrifugation at 10,000 × g for 10 minutes at 4°C or store at -80°C for subsequent use. Next, for co-IP, 500μL of the lysate, containing antigen sample, was mixed with 10 μg of antibody of Human TNFSF13 Antibody (Cat# MAB8844, R&D Systems) or Mouse IgG2B Isotype Control (Cat# MAB004, R&D Systems) and incubated overnight at 4ºC with constant shaking. 25μL (0.25mg) per sample of Sera-Mag™ SpeedBeads Magnetic Protein A/G Particles (Cytiva) were prewashed in 175 μL and 1 mL of wash buffer (25 mM Tris, 0.65 M NaCl, 0.05% Tween-20 detergent- Promega, pH 7.5) with gently vortex to mix, in turn. The particles were subsequently collected using a magnetic stand. These pre-washed magnetic beads were introduced into the antigen sample/antibody mixture and incubated at room temperature for 1 hour with mixing. Following this incubation, the beads were collected with a magnetic stand and washed for three times, each with 500μL of wash buffer. After the final wash, the beads were resuspended in 100μL of SDS-PAGE reducing sample buffer (GenScript, New Jersey, USA) and heated either at 96-100ºC in a heating block for 10 minutes or at room temperature for 10 minutes with mixing if the samples were intended for Western blot with a rabbit antibody (primary or secondary). Subsequently, the beads were separated magnetically, and the supernatant containing the target antigen was carefully collected and stored at -20°C.

NuPAGE™ 4 to 12%, Bis-Tris, 1.0–1.5 mm, Mini Protein Gels (Thermo Fisher Scientific) was employed for detecting FAS protein in co-IP supernatant containing target antigen or other soluble proteins in colonoids/organoids according to the manufacture’s protocol. Briefly, appropriate samples and pre-stained protein marker

(ApexBio, Texas, USA) were loaded into a NuPAGE mini Protein Gel and then was carried out electrophoresis with 1X NuPAGE® SDS Running Buffer (200 mL for the Upper Buffer Chamber and 600 mL for the Lower Buffer Chamber, Thermo Fisher Scientific) at a constant voltage of 200V for 50 min within a XCell SureLock™ Mini-Cell system (Thermo Fisher Scientific). Subsequently, proteins were transferred and assembled to a pre-soaked PVDF membrane (pre-soaking firstly with methanol for 30s and washing with water, and then immersing in 2 x NuPAGE® Transfer Buffer for 5 min, Thermo Fisher Scientific) using Trans-Blot Turbo Transfer System (Bio-Rad, California, USA) for 30-60 min under standard conditions. After transfer, the membrane was blocked with 5% dry milk (Lab Scientific, New Jersey, USA) in TBST (0.1% Tween-20 in 1x TBS buffer) for 1-2 hours at room temperature on a shaker, and then incubated with diluted Recombinant anti-FAS antibody (EPR5700) (Cat# ab133619, Abcam, Cambridge, UK), anti-BCL-XL (Cat# 2762S, Cell Signaling Technology, Massachusetts, USA), anti-β-ACTIN (Cat# A5316-.2ML, Sigma-Aldrich) in blocking buffer overnight at 4°C on a shaker. After being washed three times for 10 min with TBST, the membrane was incubated with diluted Peroxidase (HRP) Anti-Rabbit IgG Goat Secondary Antibody (Cat# 7074S, Cell Signaling Technology), or Rabbit anti-Mouse IgG (H+L) Secondary Antibody [HRP] (Abcam or Novus Biologicals, Colorado, USA), or Anti-mouse IgG VeriBlot for IP secondary antibody (Cat# ab131368, Abcam) in blocking buffer for 1 hour at room temperature. Protein detection was facilitated by a luminol-based detection reagent (Santa Cruz Biotechnology, Texas, USA). Subsequently, the membrane was imaged using a Gel Doc XR+ Gel Documentation System (Bio-Rad). Detailed antibodies information was listed in Supplementary Table 2.

*RNA isolation and qRT-PCR.* Total RNA was isolated with Quick-RNA™ Miniprep Kit (ZYMO RESEARCH) from human PBMCs, patient-derived colonoids and iPSC-derived organoids and then subjected to synthesize cDNA using random hexamers with TaqMan™ Reverse Transcription Reagents (Thermo Fisher Scientific) according to the manufacturer's instructions. Real-time quantitative PCR (qPCR) was performed either with the Power SYBR Green Master Mix or TAQMAN Fast Gene Expression Universal PCR Master Mix 2X (for Taqman probes) (Applied Biosystems) on QuantStudio 3 and/or 5 Real-Time PCR Systems (Thermo Fisher Scientific) with specific primers (Supplementary Table 5) according to the manufacture's protocol. GAPDH was used as an internal control for normalization. The relative gene expression levels were calculated using the formula $2^{-\Delta\Delta Cp}$ method. Microsoft excel and GraphPad Prism9 software were used to process the qPCR data.

*Neutralization experiments.* Experiments used 1 μg/mL TNFSF13 (Cat #MAB5860, R&D Systems), 10 μg/mL recombinant Human TNFSF13 (HEK293-expressed) protein (Cat #5860-AP-010, R&D Systems), Mouse IgG1 Isotype Control (Cat # MAB002, R&D Systems), or 5 μg/mL FAS (clone ZB4, Cat #05-338, Millipore-Sigma, Massachusetts, USA) and Mouse IgG1 Negative Control (Millipore-Sigma) antibodies in media. Media was changed every other day. For patient-derived colonoids and iPSC-derived organoids, neutralizing antibodies were introduced from day 0 and continued until the time of sample collection. Detailed antibodies information is listed in Supplementary Table 2.

*ELISA.* To evaluate secreted TNFSF13 protein, 300 clusters derived from all human colonoids, or 2000 clusters derived from iPSC-organoids lines were plated in 30 µL Matrigel and feed with 500 µL media in a 24-well plate with standard passage protocol. 3 wells with 500 µL media, devoid of organoids, were designated as negative control on the same plate, simultaneously. Cells, were harvested at d4 post-seeding with the aforementioned protocol. To obtain both cytosolic and solubilized membrane and membrane-associated proteins from the same sample, the Mem-PERTM Plus Membrane Protein Extraction Kit (Thermo Fisher Scientific) was employed according to the manufacture's protocol. Briefly, cell pellet was resuspended and incubated in 0.75mL of Permeabilization buffer with protease and phosphatase inhibitors (Thermo Fisher Scientific) for 10 minutes at 4°C with constant mixing after being washed with 3mL and 1.5mL of Cell Wash Solution successively, supplemented with protease and phosphatase inhibitors, followed by centrifugation at 300 × g for 5 minutes at 4°C. The supernatant, containing cytosolic proteins, was collected and stored at -80°C after being centrifuged at 16,000 × g for 15 minutes at 4°C. Meanwhile, the resulting pellet was reconstituted in 0.5mL of Solubilization Buffer along with protease and phosphatase inhibitors and incubated at 4°C for 30 minutes with constant mixing. After a centrifugation step 16,000 ×g for 15 minutes at 4°C, the supernatant, containing solubilized membrane and membrane-associated proteins, was transferred and stored at -80°C. Furthermore, media that contained secreted proteins were collected and stored at -80°C following centrifugation at 300 × g for 5 minutes at 4°C. For B cell differentiation assays, 3000 cell clusters derived from human colonoid lines were plated in 45 µL Matrigel and fed with 500 µL media in a 24-well plate. Conditioned media containing secreted proteins were collected at day 4 post-seeding or day 9 post-seeding and stored at -80°C following centrifugation at 300 × g for 5 minutes at 4°C, respectively.

The Human APRIL/TNFSF13 DuoSet® ELISA Development System (Cat# DY884B) was employed to measure the expression level of TNFSF13 in colonoids/iPSC-organoids or media according to the manufacture's protocol. Briefly, the sealed plate was incubated with 100 µL per well of the diluted Capture Antibody overnight at room temperature and washed three times with 400 µL per well of Wash Buffer. After blocking with 300 µL of Reagent Diluent per well for a minimum of 1 hour at room temperature, the washed plate was treated with 100 µL per well of either sample or standards in Reagent Diluent and incubated for 2 hours at room temperature with an adhesive strip covering. The standard was reconstituted in Reagent Diluent with a concentration gradient (2000 pg/mL, 1000 pg/mL, 500 pg/mL, 250 pg/mL, 125 pg/mL, 62.5 pg/mL, 31.3 pg/mL, 0 pg/mL). Then, the plate was incubated with 100 µL of the Detection Antibody for 2 hours, followed by 100 µL of the working dilution of Streptavidin-HRP for 20 min, and 100 µL of Substrate Solution for 20 min, in turn. The plate was washed 3 times with 400 µL per well of Wash Buffer between these steps. At the end of the final wash, 50 µL of Stop Solution was added to each well and mixed thoroughly by gently tapping the plate. The optical density (O.D.) of each well was determined promptly under 450 nm with wavelength correction setting at 570nm, using a GloMax®-Multi Detection System (Promega, Wisconsin, USA) or Varioskan LUX multimode microplate reader. The corrected O.D. is the reading at 450 nm subtract readings at 570 nm. The process and analysis of data was used an four parameter curve fit (4PL) online tool (https://www.aatbio.com/tools/four-parameter-logistic-4pl-curve-regression-online-calculator) and the equation form was created as follows based on the standard curve (Y=absorbance O.D., X=concentration of TNFSF13 in pg/mL):

$$Y = Min + \frac{Max - Min}{1 + \left(\frac{X}{inflection\ Point}\right)^{Hill\ coefficient}}$$

To evaluate secreted IgA protein, media was collected at day 14 post-seeding from differentiated memory B cells and stored at -80°C following centrifugation at 2000 × g for 10 minutes at 4°C. In parallel, 3 wells containing 150 µL media, devoid of cells, were designated as negative control on the same plate. The Human IgA ELISA Kit (Cat# ab196263) was used according to the manufacture's protocol. Briefly, 50 µL of diluted samples (1:1 into Sample Diluent NS) and/or standard were added to appropriate wells of SimpleStep Pre-Coated 96-Well Microplate. Two blank wells were used as the zero control. Each sample was assayed with two technical replicates. After adding another 50 µL of the Antibody Cocktail to each well, the plate was sealed and incubated for 1h at room temperature on a plate shaker set to 400 rpm. Following a wash step with 3 x 350 µL 1X Wash Buffer PT, 100 µL of TMB Development Solution was added to the washed plate and incubated for 10 minutes in the dark on a plate shaker set to 400 rpm. At the end of the incubation, 100 µL of Stop Solution was added, and absorbance at 450 nm wavelength measured using a GloMax®-Multi Detection System (Promega) or Varioskan LUX multimode microplate reader. The concentration was determined as described above.

Due to the variation in the percentage of B cells and memory B cells from each donor, we seeded 37,000 -100,000 human memory cells per well in our assays. As such, the absolute values of ELISA between two assays were not directly comparable. However, we ensured equal cell numbers between control and case in each assay for meaningful comparisons.

*Single cell RNA sequencing (scRNA-seq) preprocessing and analysis.*

*Sample preparation.* Human colonoids (2 control, 2 VEO-IBD, 1 variant with 2 replicates from different passage and batch) were collected and digested into single cells as shown above. The cell pellet was resuspended and incubated in 45 µL of FACS buffer and 5 µL of Human TruStain FcX™ Fc Blocking reagent (BioLegend) for 10 min at 4°C. And then 1µg of TotalSeqtrade-B0251 anti-human Hashtag 1 Antibody (BioLegend) was introduced for untreated samples (TotalSeqtrade-B0252 anti-human Hashtag 2 Antibody for TNF$\alpha$-treated sample, TotalSeqtrade-B0253 anti-human Hashtag 3 Antibody for IFN-$\gamma$ treated sample and TotalSeqtrade-B0254 anti-human Hashtag 4 Antibody for Trail treated sample) in FACS buffer (up to 50 µL) and incubated for another 30 min at 4°C after centrifuging the antibody pool at 14,000 x g for 10 minutes at 4°C. Afterward, the cells were washed three times with 3 mL FACS buffer, centrifuged at 4°C for 5 minutes at 300 x g, and then incubated with DAPI before being sorted in 1xDPBS+0.04% BSA with a MoFlo Astrios sorter (Beckman Coulter) or FACSAria Fusion Sorter (BD Biosciences) in CHOP Flow Core, as shown above, to isolate live single cells. Equivalent cell number from sorted untreated and treated samples were combined to create a sample pool for library construction. Only data from untreated samples was used in this study.

For fresh tissue from human colon biopsy, colonic biopsies were collected during colonoscopy and placed into Eppendorf tubes containing 1mL of collection medium (comprising 1% Pen/Strep and 1x HEPES and 1x GlutaMAX in Advanced DMEM/F12). These samples were promptly transported to the lab immediately for processing or cryopreservation. For isolation of crypts and lamina propria, the biopsies were washed twice with 1mL of Chelation buffer before incubating in 1mL Chelation buffer along with 2 mM EDTA for 15min at 4°C with rotation. The tissue was then vortexed using a vortex mixer (Scientific Industries) at the maximum speed setting for 5 cycles, with 30 seconds of vortexing followed by a 30-second rest period on ice. After vortexing, the tissue, along with the supernatant, was transferred into a cold 35 mm dish with the luminal side (epithelium) facing upward. The crypts were meticulously scraped off the tissue using two Dumont SS forceps (Angled) under a Stereo microscope (Aven tools). The crypts contained in solution were then collected into a 15 mL conical and spun down at 700x g for 1min, followed by digestion steps in 1.5 mL TrypLE along with 0.5 U/mL DNAse I (Roche, Basel, Switzerland) for 30 min at 37°C with 800 rpm using a ThermoMixer (Thermo Fisher Scientific). After deactivation with 10%FBS and centrifugation at 700x g for 3min, the epithelial cells were reconstituted in 500 µL of 1%BSA in 1x HBSS, supplemented with 0.5 U/mL of DNAse I. Meanwhile, following rinsing twice with 5% FBS in cold DMEM/F12 (the solution was collected in the crypts collection conical), the lamina propria fraction was digested in 500 µL of stroma dissociation enzyme mix, comprising with 0.13 WU/mL of Liberase-TH (Cat #5401135001, Sigma-Aldrich),

0.5U/mL DNAse I in 1x HBSS, for 30 min at 37°C with 800 rpm using a ThermoMixer. The resulting solution was then passed through a 40 μm strainer placed in a 50mL conical to collect supernatant. So, after two more cycles, the remaining fragments was additionally forced through the strainer using a plunger from a 1 mL insulin syringe. A wash with 4 mL of 5% FBS in cold DMEM/F12 was performed to collected any remaining cells from the strainer in the same conical. Following centrifugation at 700x g for 3min, the lamina propria cells was reconstituted in 500 μL of 5% FBS in cold DMEM/F12. Cell number and viability assessments for both epithelial cells and lamina propria cells were determined by Trypan Blue (Gibco, Massachusetts, USA) exclusion using a Countess™ 3 FL Automated Cell Counter (Thermo Fisher Scientific). If cell viability was lower than 70%, the Dead Cell Removal kit (Cat #130-090-101, Miltenyi Biotech) was employed following the manufacturer's instructions. After viability checks, the cells were ready for subsequent library construction steps.

*Library preparation.* Chromium Next GEM Single Cell 3' Reagent Kits v3.1 with Feature Barcoding technology for Cell Surface Protein (10X GENOMIC, California, USA) were employed for library construction according to the manufacture's protocol. Briefly, a total of 43.2 μL of 10,000 cells diluted in nuclease-free water (Thermo Fisher Scientific) (viability ≥90% for colonoid samples and ≥71% for biopsy samples) from above sample pool were combined with 31.8 μL Master Mix (comprising 18.8 μL RT Reagent B, 2.4 μL Template Switch Oligo, 2.0 μL Reducing Agent B, 8.7 μL RT Enzyme C) and transferred to an assembled Chromium Next GEM Chip G, which was filled with Gel Beads or 50% Glycerol (VWR), and the setup was processed in a Chromium Controller (10X GENOMIC) for ~18 min. Upon completion of the run, 100 μL of the Gel

Beads-in-emulsion (GEMs) was carefully transferred to a new tube and incubated in a thermal cycler (53°C 45min, 85°C 5min, 4°C). Following by recovering with 125 µL recovery agent for 2 min, the remaining GEMs were incubated with Dynabeads Cleanup Mix (182 µL Cleanup Buffer, 8 µL Dynabeads MyOne SILANE, 5 µL Reducing Agent B, 5 µL Nuclease-free Water) for 10 min at room temperature and then eluted with Elution Solution I (98 µL Buffer EB, 1 µL 10% Tween 20- Bio-Rad, 1 µL Reducing Agent B) following the standard steps as shown in protocol. 35 µL sample was further processed for cDNA Amplification by mixing with 50 µL Amp mix and 15 µL Feature cDNA Primers 2 in a thermal cycler (98 ºC 3min, 98 ºC 15s, 63 ºC 20s, 72 ºC 1min, 10 cycles, 72 ºC 1min, 4 ºC). The resulting cDNA was then subjected to Pellet Cleanup (for 3' Gene Expression library) and Supernatant Cleanup (for Cell Surface Protein library) with SPRIselect reagent (Beckman Coulter), following standard steps as shown in the protocol. Concentration of the samples obtained from Pellet Cleanup were determined using a Qubit 4 Fluorometer (Thermo Fisher Scientific) and Qubit dsDNA HS Assay Kit (Thermo Fisher Scientific) for subsequent Sample Index PCR step (with concentrations of 25-30ng/µL for Pellet Cleanup samples and 3-10ng/µL for Supernatant Cleanup samples). Next, 10 µl of purified cDNA sample obtained from Pellet Cleanup were subjected to Fragmentation, End Repair & A-tailing by mixing with 25 µL Buffer EB (Qiagen) and 15 µL Fragmentation Mix (comprising with 5 µL Fragmentation Buffer and 10 µL Fragmentation Enzyme) in a thermal cycler (4 ºC, 32 ºC 5min, 65 ºC 30min, 4 ºC) and Post Fragmentation, End Repair & A-tailing Double-Sided Size Selection using SPRIselect reagent with steps as shown in the protocol. Following size selection, 50 µL of resulting samples were subjected to Adaptor Ligation by mixing with 50 µL Adaptor

Ligation Mix (20 µL Ligation Buffer, 10 µL DNA Ligase, 20 µL Adaptor Oligos) in a thermal cycler (20 ºC 15min, 4 ºC) and Post Ligation Cleanup using SPRIselect reagent following the protocol. The samples were then subjected to Sample Index PCR by mixing 60 µL Sample Index PCR Mix (50 µL Amp mix, 10 µL SI Primers) in 30 µL sample and 10 µL of an individual Single Index for each well (recording the well ID that was used) in a thermal cycler (98 ºC 45s, 98 ºC 20s, 54 ºC 30s, 72 ºC 20s, 11 cycles, 72 ºC 1min, 4 ºC) and Post Sample Index PCR Double Sided Size Selection using SPRIselect reagent following the protocol. Finally, 35 µL samples of 3' Gene Expression library can be stored at -20°C for long-term storage or for sequencing. For Cell Surface Protein Library Construction, 5 µL of DNA sample obtained from the Transferred Supernatant Cleanup step was subjected to Sample Index PCR by mixing with Sample Index PCR Mix (50 µL Amp mix, 35 µL Feature SI Primers 2) and 10 µL of an individual Single Index to each well (record the well ID that was used) in a thermal cycler (98 ºC 45s, 98 ºC 20s, 54 ºC 30s, 72 ºC 20s, 9 cycles, 72 ºC 1min, 4 ºC) and Post Sample Index PCR Double Sided Size Selection using SPRIselect reagent, flowing the protocol. At the end, 40 µL samples of 3' Gene Expression library can be stored at -20°C for long-term storage or for sequencing. Both the libraries for 3' Gene Expression and Cell Surface Protein were subsequently submitted to CHOP Center for Applied Genomics for sequencing using NovaSeq platform (Illumina, California, USA).

*Preprocessing and annotation of single-cell RNA-Seq data- colonoids.* Fastq files were generated using 10X Genomics Cell Ranger v6.0.0.0 then aligned to the GRCh38 human reference genome. HTO data was normalized using the centered log ratio transformation then hashtag demultiplexing of the pooled samples was performed using the HTODemux algorithm, with default settings, in Seurat v4.3.0(1). Singlets were then extracted and used for downstream analyses. Seurat was used for quality control, sample normalization, dimensional reduction, data integration, clustering, and visualization. First, cells containing <200 genes or having >25% mitochondrial gene expression were removed. The R package DoubletFinder(2) was then used to remove doublets that had not been previously detected. Data was normalized and variance stabilized using SCTransform v2 prior to integration of the different scRNA-seq datasets. To identify cell clusters, PCA was performed followed by nearest-neighbor graph construction using the first 40 principal component dimensions. Clusters were then identified using the FindClusters algorithm set to a resolution of 0.4. Clusters were annotated using markers identified with the FindAllMarkers function as well as manual visualization of known cell markers from the literature. Clusters with less than 5 DEGs (min.diff.pct = 0.25) were merged (Supplementary Table 3-4).

*Preprocessing and analysis of biopsy single-cell RNA-Seq data.* Epithelial and stromal samples were analyzed separately. Fastq files were generated as described above. Ambient RNA was removed using SoupX v1.6.2(3). The adjusted counts were then converted to Seurat objects and similar QC and processing steps were applied as described above, with the exception for mitochondrial cutoffs, which were set to 85% and 30% for epithelial and stromal samples, respectively. For the epithelial samples, major clusters were identified at a resolution of 0.1. Epithelial cells were then extracted and samples were re-integrated and processed. Initial clustering identified 15 clusters including 1 mitochondrial, 1 ribosomal and 1 stress cluster, which were excluded from further analysis. For the stromal samples, major clusters were identified at a resolution of 1.1. Five B cell clusters were identified within the data and were extracted for further analysis. Samples were re-integrated and processed. Final clustering revealed 10 subclusters including 3 B cell and 7 plasma cell clusters.

*Differential expression analysis.* For colonoid data, we performed a combined differential expression analysis by comparing cells from all clusters across the different phenotypes. DEGs were identified using the FindMarkers function in Seurat (min.diff.pct = 0.1). We used clusterProfiler v4.7.1.1(4) to identify significantly enriched KEGG pathways(5) and biological processes using the Benjamini-Hochberg adjusted p-value for multiple test correction set to <.05. Data are deposited in GEO. The accession number is GSE243445.

We evaluated 4,805 and 4,277 cells, respectively, from 2 independent healthy control and 2 TNFSF13 wild type VEO-IBD colonoid lines, and 4,682 cells from 2 different passages of TNFSF13 variant colonoids.

*Imaging Mass Cytometry (IMC).* Formalin-fixed, paraffin embedded tissue sections from a total of 7 patient samples (3 controls, 3 VEO-IBDs, and 1 TNFSF13 variant with 2 replicates) were subjected to analysis using IMC. Tissue sections were stained with a metal-conjugated antibody cocktail using previously published methods(6) (Supplementary Table 5). We acquired 1-3 regions of interest (ROIs) per tissue section, resulting in a total of 13 images for quantitative analyses. *Cell segmentation*: For cell segmentation, we used the pixel classification feature in Ilastik software(7) and trained the software to distinguish nuclear signals from the background, using the Iridium-191 channel of each IMC image. Probability maps from the nuclear pixel classification step were extracted to segment the nuclei using Cell Profiler software (8). The nuclear segmentation was expanded 6 pixels outward to approximate cell surface. Cell segmentation masks were extracted for performing cell annotation.

*Cell annotation:* Cell annotation was performed using the object classification feature in Ilastik. First, cells were categorized into the following major cell categories: T cells (CD3+), B cells/PCs (CD20+/CD27+/CD38+), dendritic cells (CD11b+), macrophages (CD68+), and monocytes (CD14+). Once these major cell type predictions were exported, we performed object classification again to subcategorize T cells into CD4+ T cells, CD8+ T cells, Foxp3+ regulatory T cells, and CD3+ T cells, and to distinguish between B cells (CD20+) and PCs (CD20-/CD27+/CD38+). We exported csv files containing the probability data for each cell, which was used to annotate the cells into 10 cell populations: CD3+ T cells, CD4+ T cells, CD8+ T cells, regulatory T cells, B cells, PCs, myeloid cells, dendritic cells, macrophages, and “non-immune” cells. A csv file containing the mean intensity of each channel for each cell after passing through a 3x3 pixel median filter was exported from Cell Profiler.

*Data filtering and analysis:* All IMC analyses were performed using R version 3.6.3. The E-cadherin channel was used to create epithelial tissue masks. Using the EBImage package in R, we categorized all cells based on their closest distance to the epithelium. All cells that were annotated as non-immune cells and were 0 μM from the epithelium were annotated to be epithelial cells, and all other non-immune cells were annotated as stromal cells. All cells that were >75um from the epithelium were excluded from the study to focus our analysis on cells that were in close proximity to the epithelium (Figure 5D, Supplementary Figure 5A). Furthermore, we used EBImage to manually remove all lymphoid tissues and submucosal tissues from downstream analysis (Figure 5D-E, Supplementary Figure 5B-C). The number of cells analyzed from each IMC image after filtering is detailed in Supplementary information, Table S5.

IMC identified 9 major immune cell populations within colon sections from 7 patients (3 controls, 3 TNFSF13 wild type VEO-IBD, and 1 TNFSF13 variant with 2 different biopsies): CD3+ T cells, CD4+ T cells (T helper cells), CD8+ T cells (cytotoxic T cells), FOXP3+ regulatory T cells (Tregs), B cells, PCs, myeloid cells, dendritic cells, and macrophages (Figure 4D, Supplementary Figure 10A-B and Table 5). Because IMC retains the X and Y coordinates of each cell in each image, we were able to assess immune cell composition with spatial resolution.

*Flow cytometry.* For human PBMCs, Ficoll-Paque Plus (GE Healthcare, Illinois, USA) was employed to isolate PBMCs from whole blood according to the manufacture's protocol. Zombie Aqua was used to distinguish live cells. B cells were identified by physical characteristics and CD19 expression, and subsets were further identified by additional antibodies: IgM, IgD, CD27, CD38, CD80 and CD21. T cells were taken from physical gate followed by CD3, and subsets were further determined using CD4, CD8, CD45RA, CXCR5, ICOS, CD25, CD161, TCR V alpha 7.2 and TCR V alpha24x18. ILCs were determined using physical lymphocyte gate and alive cells from DAPI negative. Followed by CD45 and lineage negative (CD19, CD3, CD1a, CD11c, CD14, CD34, CD123, BDCA2, and FceRI). Next is CD127 positive. CRTH2 negative is used later to determine ILC1 and ILC3 in combination with cKit and NKp44. CRTH2 positive is ILC2. Monocytes were visualized from first the physical gate followed by CD14. CD16 was analyzed from the lymphocyte, monocyte, and neutrophil physical gates. Analysis was carried out on an LSR Fortessa analyzer (BD Biosciences) in the CHOP Flow Core.

For colonoids and organoids, appropriate cells at day 7 post-plating were collect and recovered from Matrigel. After being digested in 0.05% trypsin (Thermo Fisher Scientific) for 10 min in 37°C bead bath (final concentration of 10% FBS was added to de-activate trypsin), cells were dissociated by pipetting up and down for several times to achieve a single-cell suspension. After centrifugation at 300xg at 4°C, cell pellet was resuspended and incubated in FACS buffer (2% FBS in DPBS) with various antibodies for 30 min in dark on ice: Apc anti-human CD267 (TACI) (1A1) (BioLegend), Brilliant Violet 421™ anti-human CD269 (BCMA) (BioLegend), FITC anti-CD95 Mouse Monoclonal Antibody (clone: DX2) (BioLegend), PE anti-HVEM (TR2) Mouse Monoclonal Antibody (clone: 122) (BioLegend). DAPI (Sigma-Aldrich) was added at a final concentration of 0.1 µg/mL for an additional 10 min. Subsequently, samples were analyzed on an LSR Fortessa analyzer (BD Biosciences) in the CHOP Flow Cytometry Core, following washing with 3 mL FACS buffer. Alternatively, propidium iodide (PI, Thermo Fisher Scientific) was added at a final concentration of 1 µg/mL after washing until analysis.

To validate the efficacy of flow antibodies for TNFSF13 candidate receptors, human PBMCs isolate from whole blood by Ficoll-Paque Plus (GE Healthcare) was used with the same panels. For monolayers, cells were collected around on d8 post-plating and digested with 100 µL TrypLE™ Express Enzyme (Thermo Fisher Scientific) (de-activated using an equal volume of 10% FBS in advanced DMEM/F12) at 37°C for 5 min. Single cell was resuspended in FACS buffer and performed with the same protocol and panels. Detailed antibodies information was listed in Supplementary Table 2.

*Statistical analyses.* Three healthy control and three VEO-IBD human colonoid lines from distinct patients were used as biological replicates. Three replicates with different passages from a single *TNFSF13* variant human colonoid line, and one WT and one variant iPSC-derived colon organoid line were used for statistical analyses. Statistical analysis of qPCR, flow cytometry analyses, organoids formation assay and quantification of positive cells in immunostaining assay were performed using two-tailed Student's t-tests, one-way ANOVA, two-way ANOVA or multi-comparison with Prism GraphPad or Microsoft Excel, and a significance threshold of $P<.05$ was utilized to determine statistical significance. Error bars denote mean ± standard deviation. The schematics were created with BioRender.com.

*Data availability.* Single cell RNA sequencing data will be deposited on a publicly available database. All other data are available from the corresponding author upon request.

#### SUPPLEMENTARY METHODS REFERENCES

### SUPPLEMENTARY FIGURES AND FIGURE LEGENDS

SUPPLEMENTARY FIGURE 1

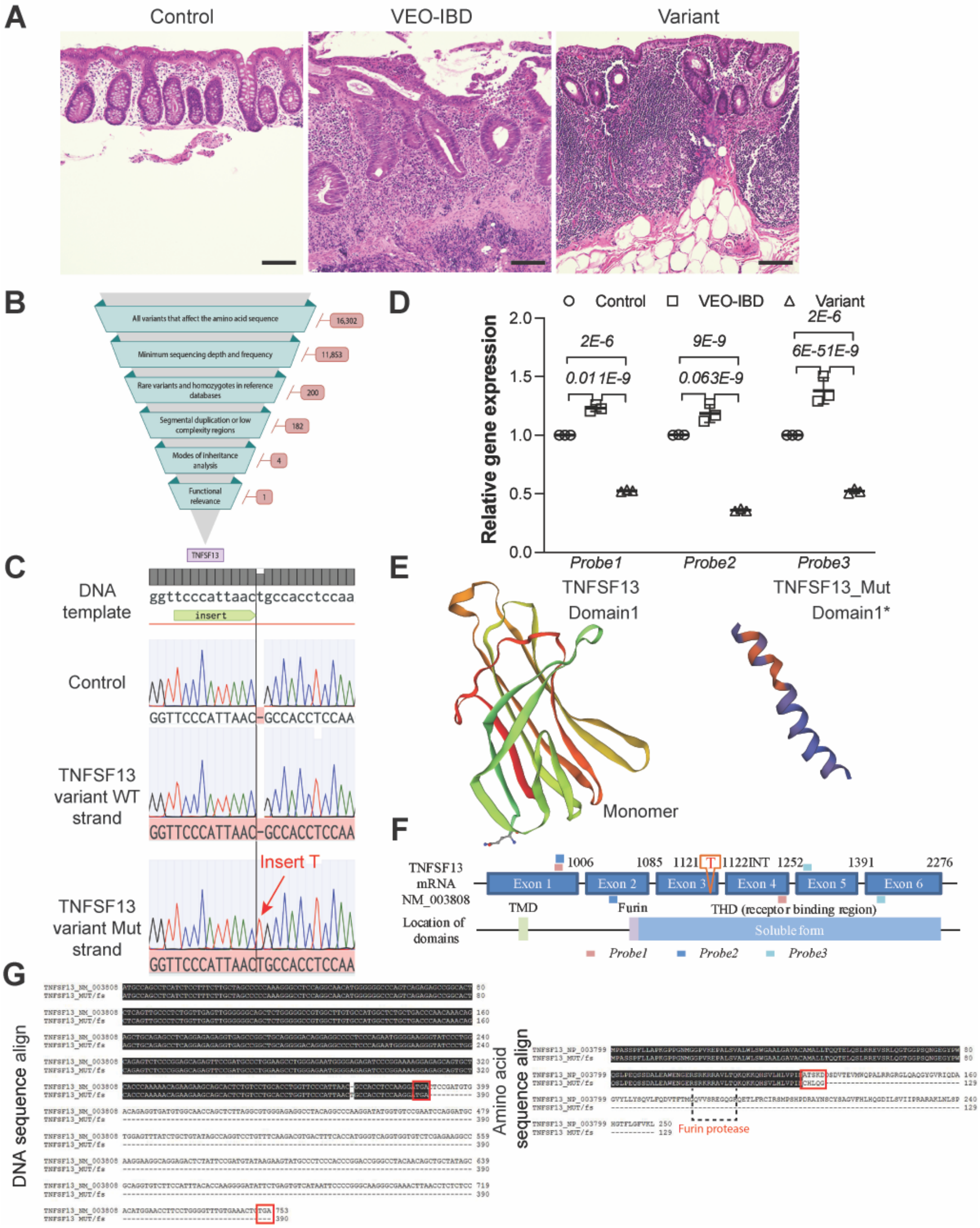

**Supplementary Figure 1 related to Figure 1. Mutation of TNFSF13 in variant both in patient-derived colonoids and iPSC-organoids. (A)** Representative H&E images of colon from control, VEO-IBD and variant. Scale bar: 100 μm. n=3 different patients for Control and VEO-IBD, n=3 slides from different blocks for Variant**.** **(B)** Diagram for filtration strategy of the whole exome sequencing. **(C)** Sanger sequencing for TNFSF13 in variant and healthy control after TOPO TA clone with colonoids, iPSC-organoids and PBMC cDNA. Red arrowhead denoted T insert in the mutant strand. **(D)** qPCR for *TNFSF13* with different probes from different location in *TNFSF13* mRNA in colonoids from control, VEO-IBD and variant. Location of the probes is shown in **(F)**. n=3 lines of colonoids from 3 different patients for Control and VEO-IBD, n=3 passages/batches of colonoids for Variant. **(E)** Images for 3D structure of TNFSF13 and predicted structure of TNFSF13 variant monomer protein. **(F)** Schematic for location of qRT-PCR probes in TNFSF13 mRNA and corresponding domains of TNFSF13 protein. *Probe1* contains the inserted site; *Probe2* is from the upper stream of the inserted site; *Probe3* is from the downstream of the inserted site. Red T denotes the insert of TNFSF13 variant. The size of exons/introns (thin line)/untranslated portions of the exons (UTR) and domains are not shown to scale. **(G)** Alignment of DNA (left) and amino acid sequences (right) of TNFSF13 and TNFSF13 variant sequence (c.372_373 T ins, NM_003808). Red box denotes stop codon (left) or new truncated amino acid sequence for variant (right). Dash line denotes the Furin protease recognizing site in TNFSF13 protein. *P* value as shown in the bar graphs unless *P*>0.05. Two-way ANOVA (with multiple comparisons) was used for statistical analysis in **(D)**.

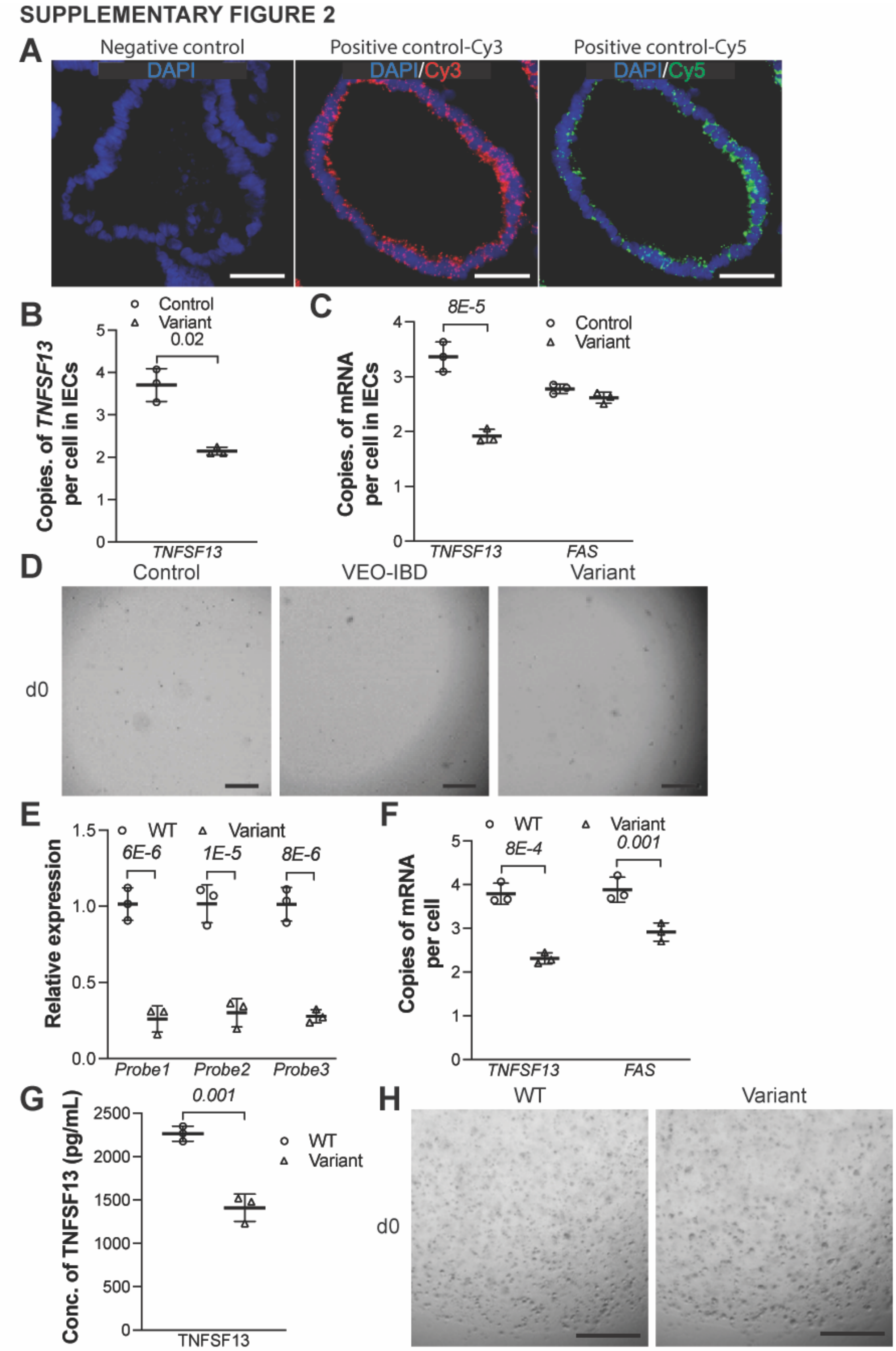

**Supplementary Figure 2 related to Figure 1. Expression of TNFSF13 in human clonoids and iPSC-derived organoids. (A)** Representative IF images for negative control and positive control RNAscope probes for *TNFSF13* (Cy3) and *FAS* (Cy5) probes in human colonoids in main figure. Scale bar: 50 µm. **(B)** Quantification of copies of *TNFSF13* (red dot) in colonnoids in main Figure 1 **(B)**. n=3 lines of colonoids from 3 different patients for Control, n=3 passages/batches of colonoids for Variant. **(C)** Quantification of copies of *TNFSF13* (red dot) and *FAS* (green dot) in IECs in main Figure 1 **(C)**. n=3 different patients for Control and VEO-IBD, n=3 slides from different blocks for Variant**. (D)** Representative images of indicated samples for colonoid formation assays on d0 post-seeding, corresponding with main Figure 1 **(E)**. Scale bar: 300 µm. n=4 lines of colonoids from 4 different patients for Control and VEO-IBD, n=4 passages/batches of colonoids for Variant. **(E)** qPCR for *TNFSF13* with different probes from different location in *TNFSF13* mRNA in WT and variant iPSC-organoids. n=3 passages/batches of iPSC-organoids. **(F)** Quantification of *TNFSF13* (red dot) in orgnoids in main Figure 1 **(F)**. n=3 passages/batches of iPSC-organoids. **(G)** ELISA for secreted TNFSF13 in iPSC-derived organoids culture conditioned media. n=3 passages/batches of iPSC-organoids. **(H)** Representative images for organoid formation assay on d0 post seeding in WT and variant iPSC-derived organoids. Scale bar: 400 µm. n=3 passages/batches of iPSC-organoids. Each passage/batch has more than two statistic replicates. *P* value as shown in the bar graphs unless *P*>0.05. Two-way ANOVA (with multiple comparisons) or two-tailed Student's *t*-test was used for statistical analysis in **(B-C) and (E-G)**.

SUPPLEMENTARY FIGURE 3

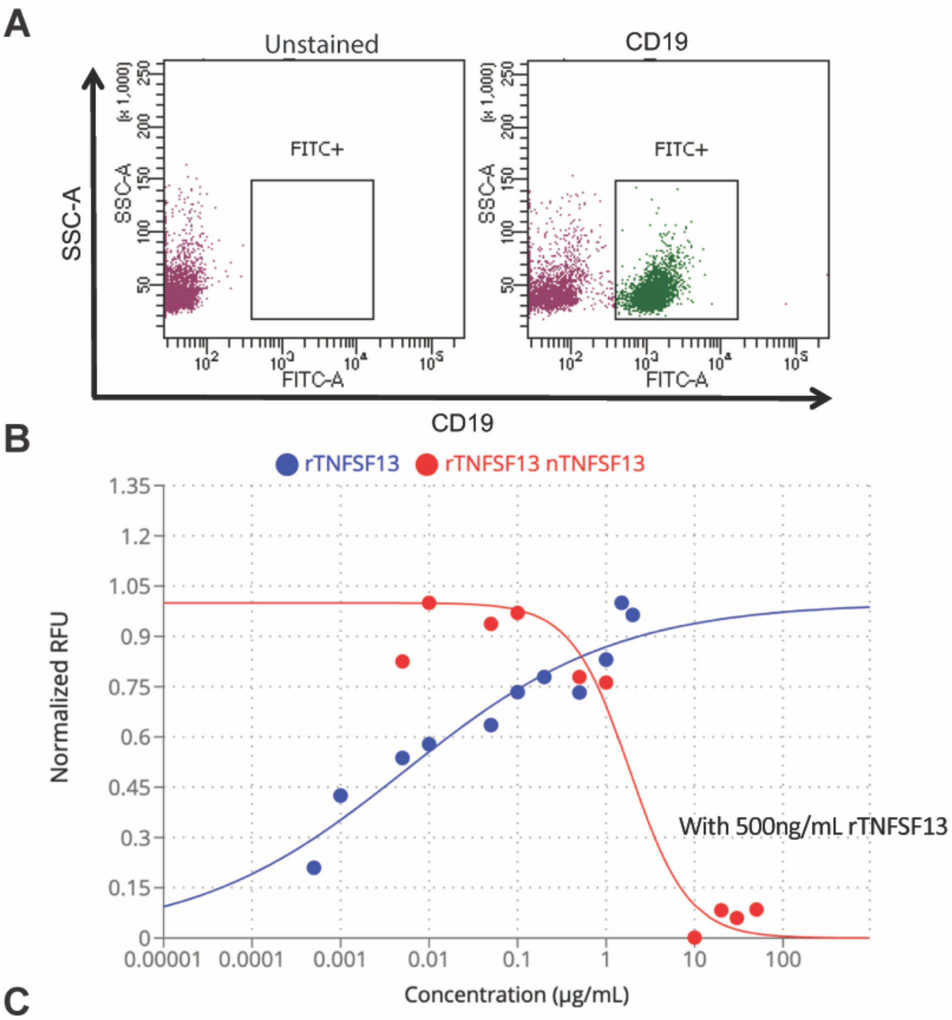

C

$X_{50}$ Regression Results [rTNFSF13]

| Parameter | Value |
|---|---|
| $X_{50}$ | 0.0054 |
| Equations | |
| Equation | $Y = 0 + \frac{1 - 0}{1 + \left(\frac{X}{0.0054}\right)^{-0.3617}}$ |
| Equation Form | $Y = Min + \frac{Max - Min}{1 + \left(\frac{X}{X50}\right)^{\text{Hill coefficient}}}$ |

$X_{50}$ Regression Results [rTNFSF13 nTNFSF13]

| Parameter | Value |
|---|---|
| $X_{50}$ | 1.8537 |
| Equations | |
| Equation | $Y = 0 + \frac{1 - 0}{1 + \left(\frac{X}{1.8537}\right)^{1.3152}}$ |
| Equation Form | $Y = Min + \frac{Max - Min}{1 + \left(\frac{X}{X50}\right)^{\text{Hill coefficient}}}$ |

**Supplementary Figure 3 related to Figure 1. TNFSF13 promote proliferation of mouse splenic B cells with dose-dependent pattern. (A)** FACS for sorting of mouse splenic DAPI$^-$CD19$^+$ B cells. **(B)** Curve chart for cell proliferation assay with resazurin in mouse splenic B cells after treatment with rTNFSF13 and/or nTNFSF13. Red line: cells were treated with gradient concentration of rTNFSF13. Red line: cells were treated with gradient concentration of nTNFSF13 and 500 ng/mL rTNFSF13. n=3. **(C)** Equation for the curve in **(B)**. Left: equation for rTNFSF13 treatment curve (blue line). Right: equation for rTNFSF13+nTNFSF13 treatment curve (red line).

#### SUPPLEMENTARY FIGURE 4

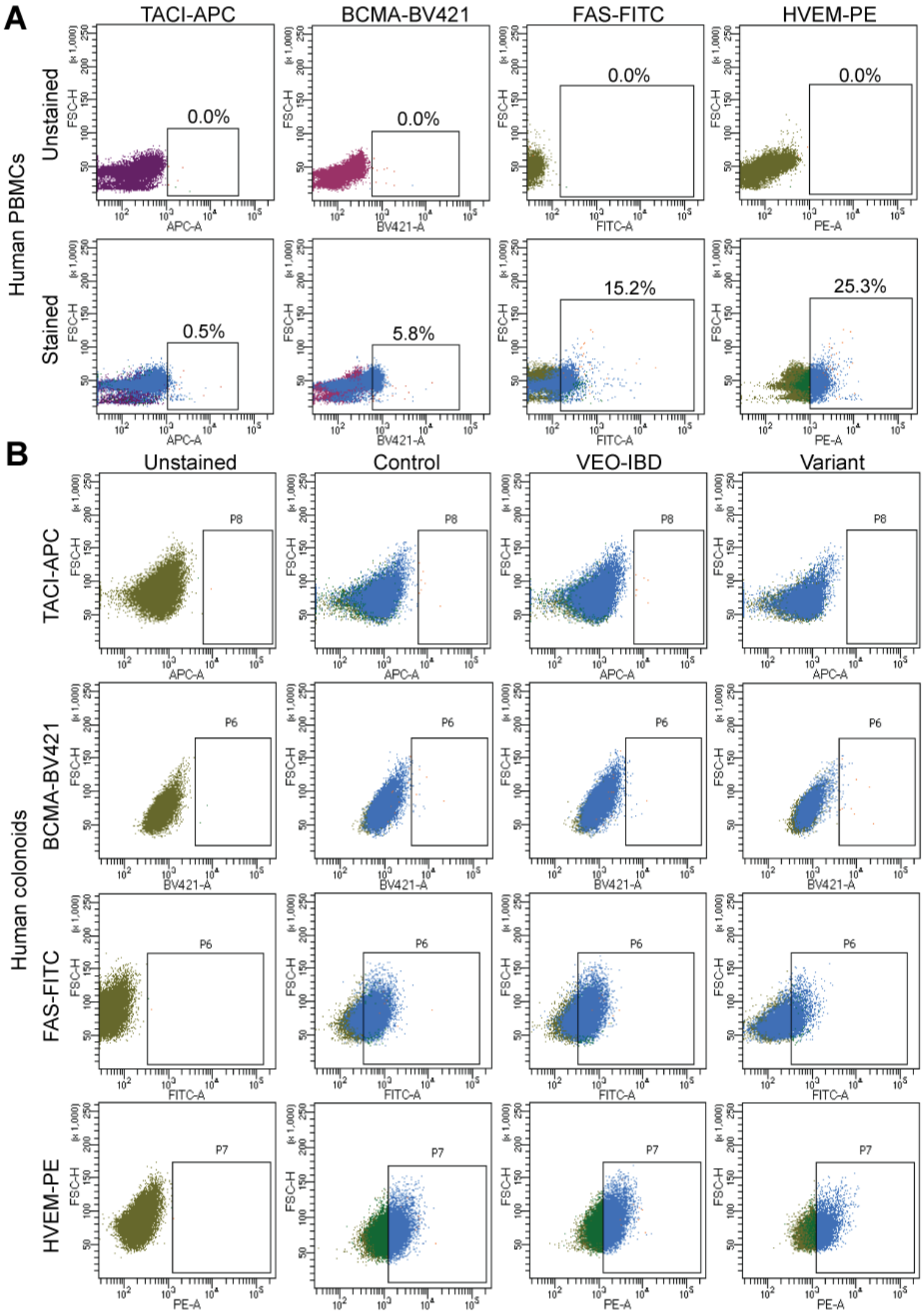

**Supplementary Figure 4 related to Figure 2. FACS strategy for TNFSF13 receptors in human PBMCs and colonoids. (A)** Verification of FACS antibodies of TNFSF13 in human PBMCs. Representative FACS images and percentage of population for TACI$^{+}$, BCMA$^{+}$, FAS$^{+}$ and HVEM$^{+}$ in DAPI$^{-}$ population. **(B)** Representative FACS images for TACI$^{+}$, BCMA$^{+}$, FAS$^{+}$ and HVEM$^{+}$ in DAPI$^{-}$ population in human colonoids of control, VEO-IBD and variant.

#### SUPPLEMENTARY FIGURE 5

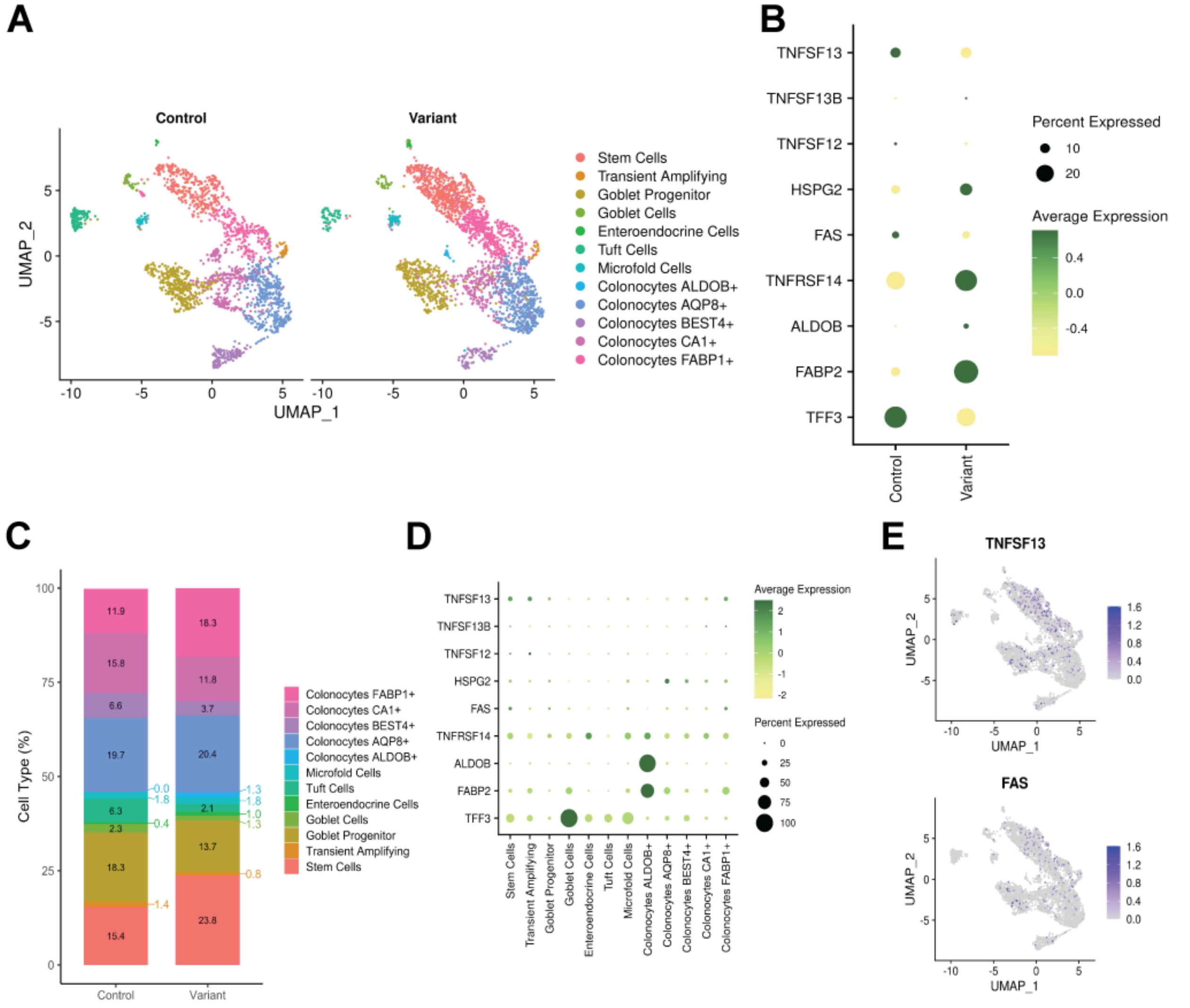

**Supplementary Figure 5 related to Figure 3. scRNAseq analysis of human colonoids and biopsy. (A)** UMAP visualizations of scRNA-seq data for epithelial cells from healthy control and variant colon biopsies. **(B)** Dot plot with relative expression of selected genes of TNFSF13 family and related receptors and enterocyte markers among control and variant in human colon biopsies. **(C)** Barplot indicated relative proportion (%) of epithelial cells in 1 of control and 1 of variant in **(A)**. **(D)** Dot plot indicated the expression pattern of selected genes of TNFSF13 family and related receptors and enterocyte markers among annotated clusters for human biopsy scRNA-seq data with relative expression. **(E)** UMAP plots showing the expression pattern of TNFSF13 and FAS in annotated epithelial cells clusters for human biopsy scRNA-seq data. n=1 patient for Control and Variant.

**SUPPLEMENTARY FIGURE 6**

**A**

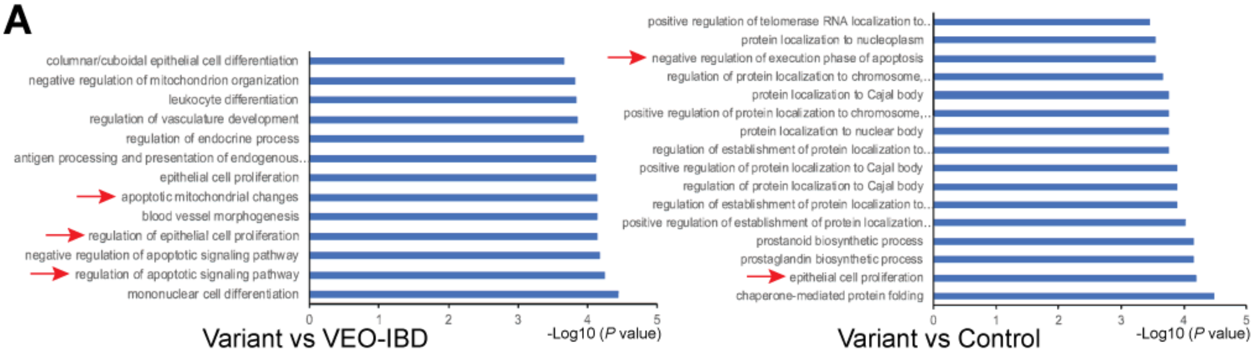

**Supplementary Figure 6 related to Figure 4. GO analysis of biological process for human colonoids scRNAseq data. (A)** GO analysis of biological process on DEGs in human colonoids from scRNA-seq data. Bar graphs showing significant changed biological processes between Variant vs VEO-IBD, Variant vs Control (VEO-IBD vs Control has no significantly changed category). Red arrow heads denote apoptosis and proliferation related categories.

**SUPPLEMENTARY FIGURE 7**

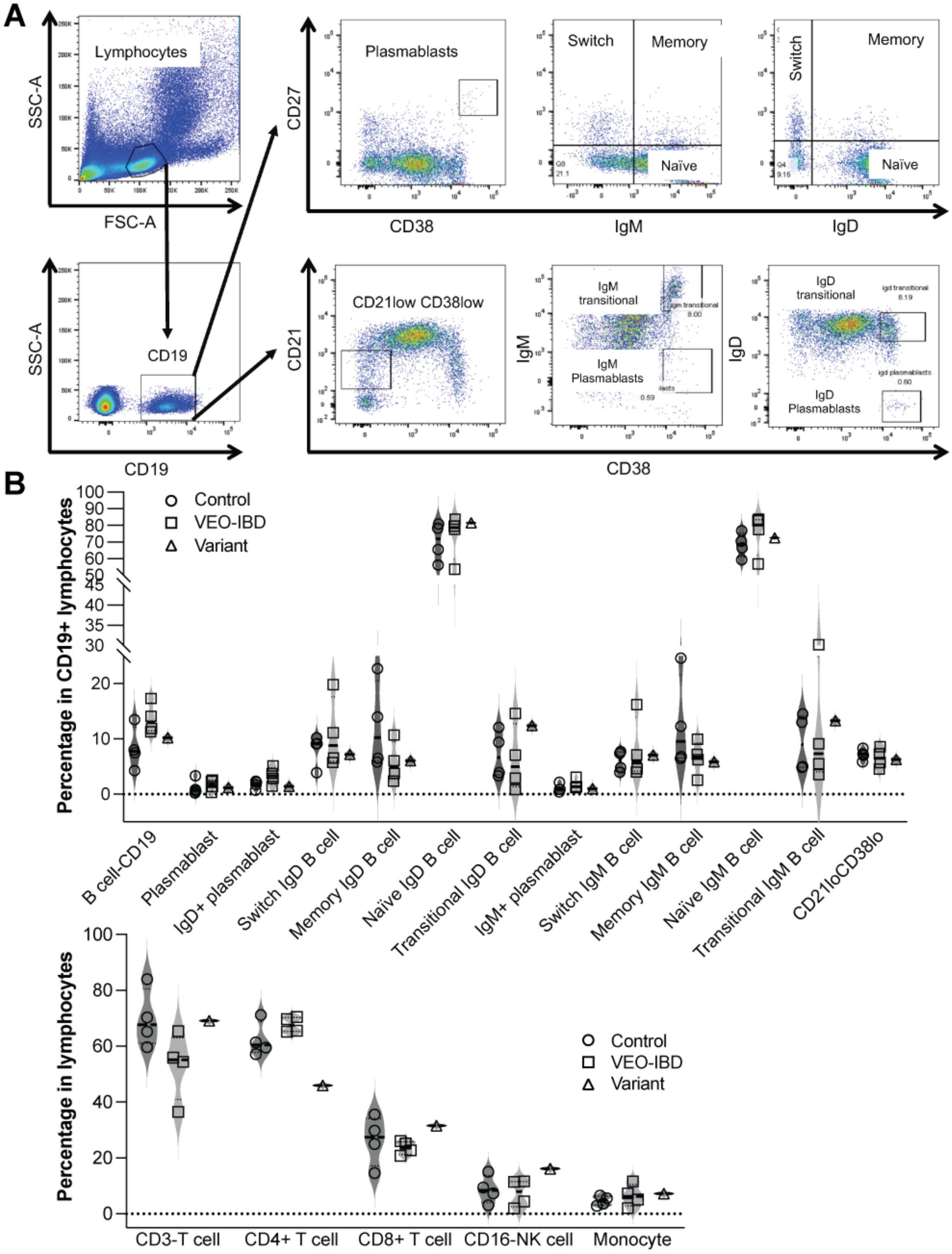

**Supplementary Figure 7 related to Figure 5. No significant differences in immunophenotyping in PBMCs between variant and non-monogenic VEO-IBD**

**subjects. (A)** FACS strategy for identifying subtypes of B cells among human PBMCs from the corresponding patients. **(B)** Percentage of subtypes of B cells in $DAPI^{-}CD19^{+}$ lymphocytes with FACS in 3 of control, 3 of VEO-IBD and 1 of variant patients. **(C)** Percentage of subtypes of other immune cells (T cells, Nature killer cells, Monocytes) in $DAPI^{-}$ lymphocytes with FACS in 3 of control, 3 of VEO-IBD and 1 of variant patients. NK cell: Nature killer cells. n=4 different patients for Control and VEO-IBD, n=1 patient for Variant. Two-way ANOVA was used for statistical analysis only between Control and VEO-IBD. *P* value was not showed if $P>0.05$.

#### SUPPLEMENTARY FIGURE 8

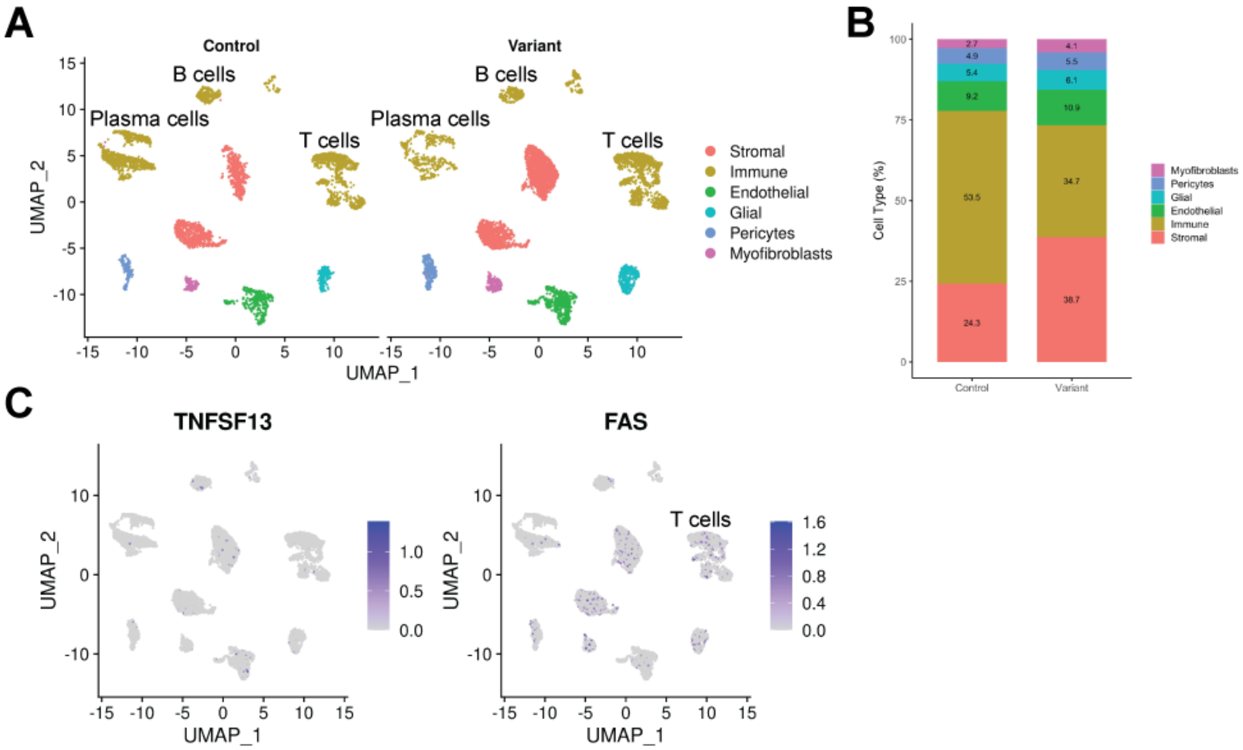

**Supplementary Figure 8 related to Figure 5. scRNAseq analysis of human biopsy. (A)** UMAP visualizations of scRNA-seq data for lamina propria cells from control and variant colon biopsies. **(B)** Barplot indicated cell type abundance (%) of lamina propria cells in 1 of control and 1 of variant. **(C)** UMAP plots showing the expression pattern of *TNFSF13* and *FAS* in annotated lamina propria cells clusters for human biopsy scRNA-seq data. n=1 patient for Control and Variant.

**SUPPLEMENTARY FIGURE 9**

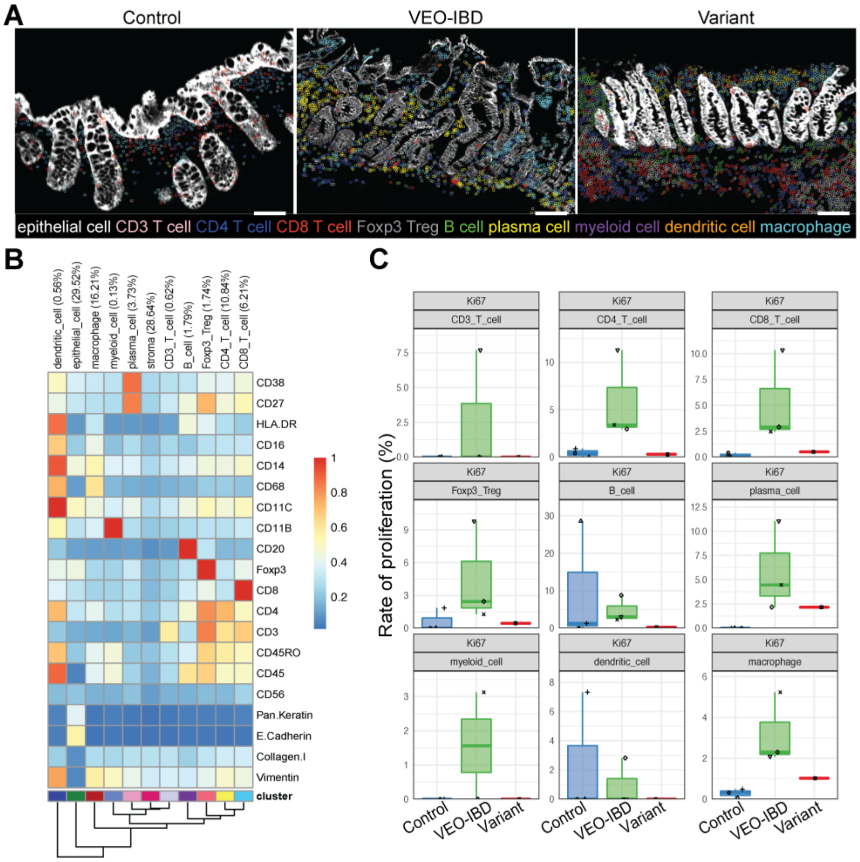

**Supplementary Figure 9 related to Figure 5. Local immune analysis in colon by IMC.** **(A)** Representative IMC overlay images for epithelial and immune cell markers in colon from 3 of control, 3 of VEO-IBD and 2 slides from different effected region of 1 variant patient. Scale bar: 100 µm. **(B)** Heatmap representing the expression profiles of the 11 annotated cell populations. Colors scale represent the average expression of a given marker in each cell population. **(C)** Boxplot showing the rate of proliferation of immune cell composition quantified by calculating the proportion of specific markers in cells that are KI67$^{+}$ in all cells at the same region (stroma + epithelium cell populations). Each point represents a sample/patient. n=3 different patients for Control and VEO-IBD, n=3 slides from different blocks for Variant.

SUPPLEMENTARY FIGURE 10

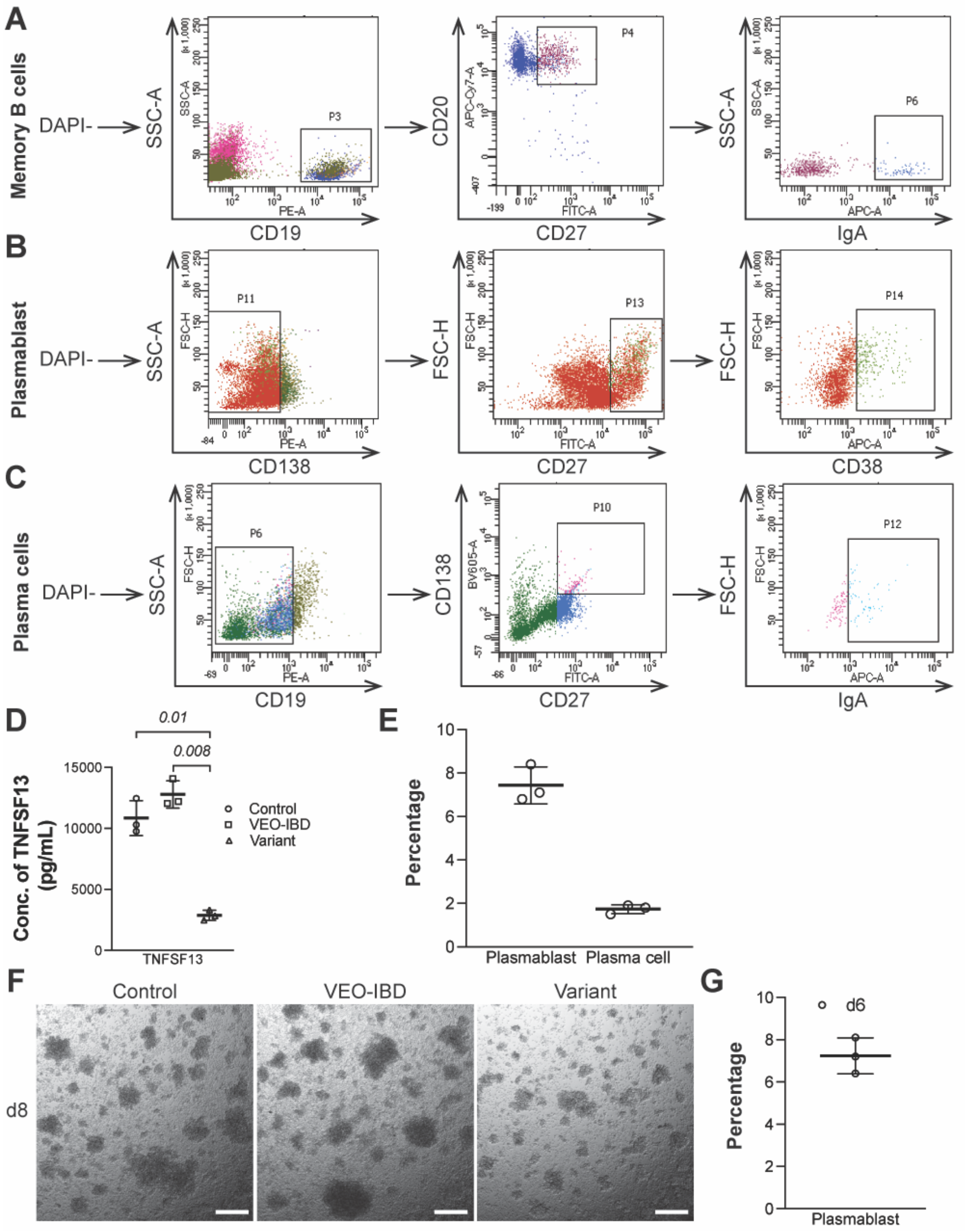

**Supplementary Figure 10 related to Figure 6. TNFSF13 regulate differentiation of memory B cells. (A)** FACS strategy for identifying memory B cells ($CD19^{+}CD27^{+}CD20^{+}$) among human PBMCs. All cells were sorted from $DAPI^{-}$

population. **(B)** FACS strategy for identifying plasmablast (CD138$^{-}$CD27$^{+}$CD38$^{++}$) at d8-post seeding differentiated from sorted human memory B cells. All cells were sorted from DAPI$^{-}$ population. **(C)** FACS strategy for identifying plasma cells (CD19$^{-}$CD27$^{+}$CD138$^{+}$) and IgA$^{+}$ plasma cells (CD19$^{-}$CD27$^{+}$CD138$^{+}$IgA$^{+}$) at d14-post seeding differentiated from sorted human memory B cells. All cells were sorted from DAPI$^{-}$ population. **(D)** ELISA for secreted TNFSF13 at d4-post seeding control, VEO-IBD and variant colonoids (3,000 clumps were seeded at d0) for co-culture and conditioned medium collection. **(E)** Percentage of plasmablast differentiated from sorted human memory B cells at d8-post seeding and plasma cells differentiated from sorted human memory B cells at d14-post seeding by culturing in B cell medium. **(F)** Representative images for differentiated memory B cell cluster at d8 post-seeding by culturing in mixture of B cell medium and conditioned medium. Clusters are growing cells. Scale bar: 300 µm. **(G)** Percentage of plasmablast differentiated from sorted human memory B cells at d6 post-seeding by culturing in B cell medium. *P* value shown in the bar graphs unless $P>0.05$. One-way ANOVA (with multiple comparisons) was used for statistical analysis. n=3 passages/batches of iPSC-organoids. n=7 independent donors to obtain human memory B cells.

**Supplementary Table 1 Patients demographics and speciement**

<table>
<tr><th>Colonoid line</th><th>Age at diagnosis (y)</th><th>Age at collection(y)</th><th>Sex</th><th>Diagnosis</th><th>Disease activity at collection</th><th>Used for</th></tr>
<tr><td>TNFSF13 variant RC-A</td><td>0.58</td><td>7.44</td><td>M</td><td>VEO-IBD</td><td>moderate</td><td>WES, TOPO TA colone, qPCR, OFR, monolayer, Western blotting, flow cytometry, scRNA-seq (colonoids), IMC, ELISA, immunostaining</td></tr>
<tr><td>TNFSF13 variant RC-A</td><td>0.58</td><td>11.09</td><td>M</td><td>VEO-IBD</td><td>moderate</td><td>scRNA-seq (biopsy)</td></tr>
<tr><td colspan="7">IBD/VEO-IBD control</td></tr>
<tr><td>VEO-IBD-1</td><td>4.33</td><td>6.33</td><td>M</td><td>VEO-IBD</td><td>Severe</td><td rowspan="3">qPCR, OFR, monolayer, Western blotting, flow cytometry, scRNA-seq, IMC, ELISA, immunostaining</td></tr>
<tr><td>VEO-IBD-2</td><td>4.08</td><td>4.34</td><td>M</td><td>VEO-IBD</td><td>moderate</td></tr>
<tr><td>VEO-IBD-3</td><td>5.5</td><td>5.5</td><td>F</td><td>VEO-IBD</td><td>Severe</td></tr>
<tr><td>VEO-IBD-4</td><td>1.83</td><td>7.4</td><td>M</td><td>VEO-IBD</td><td>moderate</td><td>FACS with PBMCs for verification of TNFSF13 receptors</td></tr>
<tr><td colspan="7">Healthy control</td></tr>
<tr><td>Control-1</td><td>N/A</td><td>6</td><td>M</td><td>N/A</td><td>N/A</td><td rowspan="3">qPCR, OFR, monolayer, Western blotting, flow cytometry, scRNA-seq, IMC, ELISA, immunostaining</td></tr>
<tr><td>Control-2</td><td>N/A</td><td>3.78</td><td>F</td><td>N/A</td><td>N/A</td></tr>
<tr><td>Control-3</td><td>N/A</td><td>5.45</td><td>F</td><td>N/A</td><td>N/A</td></tr>
<tr><td>Control-4</td><td>N/A</td><td>3.97</td><td>M</td><td>N/A</td><td>N/A</td><td>flow cytometry (PBMCs)</td></tr>
<tr><td>Control-5</td><td>N/A</td><td>10.13</td><td>M</td><td>N/A</td><td>N/A</td><td>TOPO TA colone</td></tr>
<tr><td>Control-6</td><td>N/A</td><td>9.78</td><td>M</td><td>N/A</td><td>N/A</td><td>scRNA-seq (biopsy)</td></tr>
</table>

#### Supplementary Table 2 Antibodies and primers

| **Antibody/probe** | **Catalog number** | **Used for** |
|---|---|---|
| Human APRIL/TNFSF13 Antibody | MAB5860 | B cell and colonoids neutralizing |
| Recombinant Human APRIL/TNFSF13 (HEK293-expressed) Protein | 5860-AP-010 | B cell and colonoids |
| Mouse IgG1 Isotype Control | MAB002 | B cell and colonoids |
| FAS | 05-338 | Neutralizing |
| Mouse IgG1 Negative Control | MABC002 | Neutralizing control for FAS |
| Apc anti-human CD267 (TACI) | #311911 | FACS |
| Brilliant Violet 421™ anti-human CD269 (BCMA) | #357519 | FACS |
| FITC anti-CD95 Mouse Monoclonal Antibody | #305605 | FACS |
| PE anti-HVEM (TR2) Mouse Monoclonal Antibody | #318805 | FACS |
| FITC anti-mouse CD19 | #115505 | FACS |
| PE-Cy7 anti-human CD326 (EpCAM) Antibody | #324222 | FACS |
| FITC anti-human CD326 (EpCAM) Antibody | #324203 | FACS |
| IgA Antibody, anti-human, APC | #130-113-998 | FACS |
| Anti-CD27 Mouse Monoclonal Antibody (FITC) [clone: M-T271] | #356403 | FACS |
| BD Pharmingen™ APC Mouse Anti-Human CD38 | #560980 | FACS |
| Anti-CD19 Mouse Monoclonal Antibody (PE) [clone: SJ25C1] | #363003 | FACS |
| Anti-CD138 Mouse Monoclonal Antibody (PE) [clone: MI15] | #356503 | FACS |
| Anti-CD20 Mouse Monoclonal Antibody (PE/Cy7®) [clone: 2H7] | #302311 | FACS |
| Anti-CD138 Mouse Monoclonal Antibody (Brilliant Violet® 605) [clone: MI15] | #356519 | FACS |
| BD Pharmingen™ APC-H7 Mouse Anti-Human CD20 | #560734 | FACS |
| F(ab')2-Goat anti-Mouse IgM (mu) Antibody | 16-5092-85 | mouse B cell stimulating |
| anti-Ki67 antibody | ab16667 | immunostaining |
| anti-E-Cadherin antibody | #610182 | immunostaining |
| anti-FABP2/I-FABP antibody | AF3078-SP | immunostaining |
| *TNFSF13* probe | #406981-C2 | RNAscope |
| *FAS* probe | # 427031 | RNAscope |
| positive control probe | #320881 | RNAscope |
| negative control probe | #320871 | RNAscope |
| Human TNFSF13 Antibody | MAB8844 | co-IP capture antibody |
| Mouse IgG2B Isotype Control | MAB004 | co-IP |
| Recombinant Anti-Fas antibody | ab133619 | Western blotting |
| BCL-XL Antibody | CST#2762S | Western blotting |
| Monoclonal Anti-β-ACTIN antibody | A5316-.2ML | Western blotting |
| specific fluorophore-conjugated secondary antibodies AffiniPure IgG | Jackson ImmunoResearch | immunostaining |
| Opal™ 690 fluorophore | #FP1497001KT | RNAscope |
| Opal™ 570 fluorophore | #FP1488001KT | RNAscope |
| Peroxidase (HRP) Anti-Rabbit IgG Goat Secondary Antibody | CST#7074S | Western blotting |
| Rabbit Anti-Mouse IgG H&L (HRP) (ab6728) | ab6728 | Western blotting |
| Rabbit anti-Mouse IgG (H+L) Secondary Antibody [HRP] | NBP1-75249 | Western blotting |
| Anti-mouse IgG VeriBlot for IP secondary antibody | ab131368 | Western blotting |

**qPCR-Taqman**

| | |
|---|---|
| TNFSF13 | Hs00601664_g1 |
| TNFSF13B | Hs00198106_m1 |
| TNFSF12-TNFSF13 | Hs01650719_m1 |

**qPCR-SYBR Green** 5' to 3'

| | **Forward** | **Reverse** |
|---|---|---|
| TNFSF13 qF1 | GGCAACCAGCTCTTAGGCG | AAGTCACGTCTTGAAACAGGAC |
| TNFSF13 qF2 | TGCCCTCTGGTTGAGTTGG | CCATTCTCCCAGGCTTCCAG |
| TNFSF13 qF3 | GGGTCAGGTGGTGTCTCG | AAGTTTCGCCCTTGCCCG |
| BCL2L1 | GAGCTGGTGGTTGACTTTCTC | TCCATCTCCGATTCAGTCCCT |
| ACAA2 | AAGTCTCACCTGAAACAGTTGA | CACGCAAACCAACATGCCT |
| ID1 | CTGCTCTACGACATGAACGG | GAAGGTCCCTGATGTAGTCGAT |
| ECM1 | AGCACCCCAATGAACAGAAGG | CTGCATTCCAGGACTCAGGTT |
| AldoB | TGTCTGGTGGCATGAGTGAAG | GGCCCGTCCATAAGAGAAACTT |

**TOPO TA colone** 5' to 3'

| | **Forward** | **Reverse** |
|---|---|---|
| TNFSF13 eF5 | AGCGTGGGGATTGTAAGC | CTGGGGTTACCTGGCTAT |

#### Supplementary Table 3 Effective cell count for scRNAseq

**Human colonoids**

| Sample ID | Pre Filtering | Post Filtering |
|---|---|---|
| Control-1 | 2,915 | 2,487 |
| Control-2 | 3,001 | 2,318 |
| VEO-IBD-2 | 2,811 | 2,394 |
| VEO-IBD-3 | 2,373 | 1,883 |
| Variant-1 | 3,159 | 2,442 |
| Variant-2 | 2,927 | 2,240 |
| in total | 17,186 | 13,764 |

**Human biopsy**

| | Pre-QC | Post-QC | Final-QC* |
|---|---|---|---|
| Variant-LPL | 8226 | 7175 | 6014 |
| Control-6-LPL | 8743 | 6434 | 4755 |
| Variant-epithelial | 6101 | 4302 | 2814 |
| Control-6-epithelial | 6409 | 3307 | 2207 |

*Final-QC: after removal of clusters based on top markers (doublets and markers limited to mitochondrial or ribosomal genes)

#### Supplementary Table 4 DEGs for scRNAseq in human colonoids

**Variant vs VEO-IBD**

| Genes | log2 FC | Pct Variant | Pct VEO | Adj P Val |
|---|---|---|---|---|
| LCN2 | 1.272 | 0.862 | 0.728 | 1.21E-170 |
| PRSS2 | 1.026 | 0.405 | 0.159 | 2.74E-158 |
| HES1 | 0.826 | 0.823 | 0.662 | 5.57E-223 |
| TFF3 | 0.768 | 0.876 | 0.740 | 4.58E-116 |
| BTG2 | 0.702 | 0.752 | 0.514 | 8.82E-204 |
| VMP1 | 0.698 | 0.894 | 0.734 | 2.28E-262 |
| ATF3 | 0.688 | 0.678 | 0.419 | 3.09E-200 |
| CD24 | 0.684 | 0.936 | 0.774 | 0.00E+00 |
| NOTUM | 0.676 | 0.317 | 0.080 | 1.31E-175 |
| KLF6 | 0.648 | 0.804 | 0.618 | 4.66E-184 |
| PRAC1 | 0.625 | 0.592 | 0.176 | 0.00E+00 |
| MMP7 | 0.620 | 0.308 | 0.161 | 3.19E-64 |
| FUT9 | 0.609 | 0.588 | 0.355 | 3.75E-152 |
| SERPINA1 | 0.562 | 0.383 | 0.175 | 1.10E-112 |
| S100A4 | 0.546 | 0.368 | 0.238 | 6.01E-50 |
| ECM1 | 0.541 | 0.423 | 0.257 | 1.72E-75 |
| IL32 | 0.526 | 0.385 | 0.136 | 9.43E-164 |
| DPYSL2 | 0.522 | 0.806 | 0.660 | 3.95E-135 |
| RHOB | 0.501 | 0.715 | 0.489 | 2.82E-157 |
| MUC1 | 0.501 | 0.531 | 0.391 | 1.32E-59 |
| ODAM | 0.499 | 0.335 | 0.114 | 1.16E-141 |
| ITGB8 | 0.431 | 0.493 | 0.267 | 1.13E-125 |
| ZFP36L1 | 0.421 | 0.795 | 0.646 | 2.25E-105 |
| PROX1 | 0.418 | 0.405 | 0.229 | 1.48E-81 |
| IVNS1ABP | 0.418 | 0.728 | 0.571 | 4.36E-97 |
| ACAA2 | 0.416 | 0.471 | 0.352 | 5.30E-50 |
| SLC38A11 | 0.410 | 0.271 | 0.072 | 1.03E-135 |
| ELF3 | 0.409 | 0.781 | 0.634 | 1.67E-89 |
| CDK6 | 0.409 | 0.753 | 0.623 | 5.28E-88 |
| DST | 0.402 | 0.576 | 0.386 | 4.33E-91 |
| NFAT5 | 0.401 | 0.587 | 0.436 | 6.85E-76 |
| NKD1 | 0.394 | 0.491 | 0.336 | 1.76E-66 |
| EGR1 | 0.393 | 0.598 | 0.445 | 1.04E-71 |
| IFI6 | 0.392 | 0.346 | 0.150 | 6.83E-108 |
| MME | 0.391 | 0.274 | 0.088 | 3.96E-114 |
| ID1 | 0.390 | 0.409 | 0.243 | 4.50E-73 |
| RPS4Y1 | 0.386 | 0.946 | 0.546 | 1.73E-128 |
| RARRES2 | 0.385 | 0.425 | 0.273 | 2.36E-60 |
| PLK2 | 0.377 | 0.588 | 0.425 | 3.54E-71 |
| TUBA1A | 0.376 | 0.427 | 0.296 | 1.34E-46 |
| IRF2BP2 | 0.367 | 0.834 | 0.691 | 1.62E-99 |
| CTNNB1 | 0.365 | 0.812 | 0.684 | 7.48E-89 |
| KDM5B | 0.358 | 0.633 | 0.452 | 1.01E-83 |
| DUSP1 | 0.352 | 0.614 | 0.462 | 3.44E-67 |
| MSI2 | 0.350 | 0.722 | 0.569 | 1.76E-82 |
| ABHD2 | 0.347 | 0.557 | 0.383 | 5.25E-78 |
| CLDN4 | 0.344 | 0.830 | 0.697 | 7.73E-84 |
| SEMA6A | 0.342 | 0.329 | 0.117 | 1.79E-126 |
| DEPP1 | 0.334 | 0.281 | 0.101 | 5.45E-102 |
| PBX1 | 0.318 | 0.419 | 0.228 | 4.33E-87 |
| TFDP2 | 0.309 | 0.504 | 0.353 | 8.54E-61 |
| TRBC2 | 0.309 | 0.347 | 0.179 | 2.02E-75 |
| KLF5 | 0.306 | 0.795 | 0.685 | 3.25E-60 |
| TUBB2B | 0.305 | 0.242 | 0.079 | 4.10E-94 |
| TCF4 | 0.303 | 0.427 | 0.270 | 1.31E-57 |
| ACTN1 | 0.302 | 0.769 | 0.655 | 2.76E-63 |
| NFIA | 0.301 | 0.695 | 0.573 | 3.22E-55 |
| RUNX1 | 0.296 | 0.518 | 0.368 | 4.83E-58 |
| CDKN1C | 0.293 | 0.338 | 0.232 | 5.63E-29 |
| SNHG14 | 0.288 | 0.499 | 0.373 | 1.29E-41 |
| TRIB1 | 0.288 | 0.589 | 0.458 | 6.75E-50 |
| GTF2I | 0.287 | 0.763 | 0.662 | 3.98E-53 |
| DUSP4 | 0.285 | 0.428 | 0.285 | 2.00E-52 |
| AL354707.1 | 0.283 | 0.277 | 0.090 | 7.13E-112 |
| LPP | 0.282 | 0.654 | 0.523 | 4.85E-52 |
| APCDD1 | 0.278 | 0.404 | 0.292 | 2.11E-30 |
| ZMYND8 | 0.277 | 0.599 | 0.460 | 2.02E-51 |
| INHBB | 0.277 | 0.267 | 0.126 | 2.98E-62 |
| ZBTB20 | 0.275 | 0.368 | 0.215 | 8.37E-59 |
| ARHGAP5 | 0.275 | 0.660 | 0.525 | 4.31E-52 |
| USP9Y | 0.270 | 0.293 | 0.114 | 3.80E-94 |
| SNHG5 | 0.267 | 0.716 | 0.575 | 1.84E-55 |
| AC022075.1 | 0.266 | 0.273 | 0.166 | 9.37E-35 |
| PTK7 | 0.265 | 0.626 | 0.507 | 9.59E-43 |
| TACSTD2 | 0.265 | 0.345 | 0.216 | 3.84E-42 |
| NUDT4 | 0.264 | 0.499 | 0.371 | 4.02E-44 |
| NET1 | 0.261 | 0.588 | 0.455 | 1.47E-46 |

**Variant vs Control**

| Genes | log2 FC | Pct Variant | Pct Control | Adj P Val |
|---|---|---|---|---|
| LCN2 | 1.914 | 0.862 | 0.512 | 0.00E+00 |
| TFF3 | 1.142 | 0.876 | 0.677 | 1.00E-259 |
| MMP7 | 0.833 | 0.308 | 0.075 | 1.26E-187 |
| MUC1 | 0.773 | 0.531 | 0.253 | 4.82E-205 |
| PRSS2 | 0.713 | 0.405 | 0.296 | 3.70E-40 |
| SERPINA1 | 0.686 | 0.383 | 0.143 | 1.87E-171 |
| KLK7 | 0.684 | 0.686 | 0.470 | 1.81E-166 |
| NOTUM | 0.622 | 0.317 | 0.116 | 1.33E-135 |
| ECM1 | 0.563 | 0.423 | 0.258 | 3.59E-83 |
| RPS4Y1 | 0.554 | 0.946 | 0.497 | 1.97E-238 |
| TCN1 | 0.513 | 0.285 | 0.139 | 2.61E-72 |
| RARRES2 | 0.466 | 0.425 | 0.256 | 3.79E-88 |
| SLC6A8 | 0.459 | 0.195 | 0.095 | 3.85E-45 |
| ODAM | 0.412 | 0.335 | 0.166 | 1.19E-87 |
| IL32 | 0.407 | 0.385 | 0.233 | 2.21E-69 |
| ID1 | 0.371 | 0.409 | 0.255 | 1.06E-68 |
| ABHD2 | 0.368 | 0.557 | 0.389 | 3.16E-87 |
| TACSTD2 | 0.366 | 0.345 | 0.149 | 1.49E-108 |
| PLK2 | 0.357 | 0.588 | 0.442 | 2.26E-67 |
| IFI6 | 0.354 | 0.346 | 0.175 | 3.90E-87 |
| TCF4 | 0.351 | 0.427 | 0.248 | 3.72E-82 |
| BTG2 | 0.351 | 0.752 | 0.644 | 3.65E-59 |
| ANXA1 | 0.347 | 0.487 | 0.347 | 4.64E-47 |
| AC020656.1 | 0.346 | 0.504 | 0.398 | 2.70E-39 |
| ITGB8 | 0.333 | 0.493 | 0.338 | 1.81E-70 |
| IGFBP3 | 0.324 | 0.420 | 0.284 | 5.02E-41 |
| ZFP36L1 | 0.323 | 0.795 | 0.693 | 8.73E-66 |
| TRBC2 | 0.318 | 0.347 | 0.178 | 2.82E-84 |
| AHNAK | 0.304 | 0.634 | 0.528 | 2.57E-41 |
| GABRA2 | 0.296 | 0.254 | 0.068 | 7.01E-131 |
| RHOB | 0.296 | 0.715 | 0.596 | 9.08E-58 |
| CD74 | 0.290 | 0.443 | 0.294 | 6.87E-60 |
| ATF3 | 0.287 | 0.678 | 0.558 | 1.80E-44 |
| ADIRF | 0.281 | 0.318 | 0.215 | 1.05E-28 |
| BCL2L1 | 0.271 | 0.525 | 0.407 | 4.13E-42 |
| USP9Y | 0.271 | 0.293 | 0.117 | 3.60E-99 |
| TSHZ2 | 0.270 | 0.299 | 0.125 | 1.68E-94 |
| IFITM2 | 0.267 | 0.190 | 0.016 | 2.06E-169 |
| GSN | 0.266 | 0.654 | 0.526 | 7.15E-55 |
| KCTD12 | 0.263 | 0.235 | 0.103 | 3.29E-66 |
| DNAJC15 | 0.257 | 0.375 | 0.208 | 2.00E-72 |
| AKR1C3 | 0.256 | 0.408 | 0.296 | 1.38E-36 |
| PDLIM4 | 0.251 | 0.334 | 0.210 | 7.81E-45 |
| MDH2 | -0.250 | 0.511 | 0.638 | 4.43E-52 |
| TXNL4A | -0.251 | 0.382 | 0.541 | 1.26E-60 |
| SNRPD1 | -0.252 | 0.510 | 0.635 | 1.14E-45 |
| TCIM | -0.255 | 0.022 | 0.156 | 1.32E-110 |
| PRDX6 | -0.255 | 0.611 | 0.735 | 2.62E-56 |
| TRAP1 | -0.255 | 0.246 | 0.422 | 6.66E-75 |
| EIF1AX | -0.256 | 0.643 | 0.767 | 3.25E-58 |
| RGMB | -0.257 | 0.333 | 0.474 | 1.74E-44 |
| CCT8 | -0.257 | 0.555 | 0.697 | 2.45E-60 |
| NASP | -0.260 | 0.568 | 0.670 | 3.80E-34 |
| ANP32B | -0.262 | 0.609 | 0.737 | 3.90E-54 |
| CFAP97 | -0.262 | 0.371 | 0.536 | 6.52E-64 |
| CYC1 | -0.264 | 0.328 | 0.484 | 1.49E-60 |
| PARP1 | -0.264 | 0.513 | 0.671 | 1.88E-59 |
| LSM4 | -0.265 | 0.519 | 0.675 | 1.87E-61 |
| TOMM5 | -0.266 | 0.429 | 0.580 | 1.99E-56 |
| MRPS34 | -0.268 | 0.443 | 0.604 | 1.31E-62 |
| ITPR2 | -0.268 | 0.420 | 0.583 | 7.82E-55 |
| CCT6A | -0.270 | 0.630 | 0.752 | 5.87E-56 |
| GSTM3 | -0.271 | 0.437 | 0.602 | 1.35E-64 |
| TOMM40 | -0.272 | 0.212 | 0.389 | 6.46E-80 |
| FERMT1 | -0.274 | 0.319 | 0.499 | 1.53E-74 |
| AURKAIP1 | -0.275 | 0.541 | 0.697 | 7.09E-70 |
| IFI27L2 | -0.276 | 0.165 | 0.357 | 1.48E-99 |
| TRMT112 | -0.276 | 0.517 | 0.670 | 5.21E-69 |
| PRKDC | -0.276 | 0.623 | 0.754 | 6.03E-56 |
| ASCL2 | -0.279 | 0.652 | 0.766 | 1.29E-40 |
| DNMT1 | -0.279 | 0.245 | 0.398 | 1.75E-57 |
| PAICS | -0.279 | 0.370 | 0.551 | 5.20E-75 |
| SNHG17 | -0.280 | 0.122 | 0.298 | 1.42E-98 |
| EBPL | -0.281 | 0.314 | 0.498 | 9.34E-80 |
| FBL | -0.282 | 0.413 | 0.589 | 1.14E-73 |
| AZGP1 | -0.283 | 0.232 | 0.411 | 3.14E-80 |
| HMGA1 | -0.283 | 0.676 | 0.798 | 8.25E-69 |

**VEO-IBD vs Control**

| Genes | log2 FC | Pct VEO | Pct Control | Adj P Val |
|---|---|---|---|---|
| OLFM4 | 1.061 | 0.607 | 0.170 | 0.00E+00 |
| IFI27 | 0.714 | 0.440 | 0.189 | 5.06E-180 |
| PAX8-AS1 | 0.713 | 0.430 | 0.007 | 0.00E+00 |
| LCN2 | 0.642 | 0.728 | 0.512 | 8.63E-113 |
| AC020656.1 | 0.516 | 0.616 | 0.398 | 3.86E-130 |
| HLA-C | 0.516 | 0.932 | 0.827 | 2.93E-184 |
| IGFBP3 | 0.506 | 0.503 | 0.284 | 9.36E-100 |
| PIGR | 0.505 | 0.376 | 0.172 | 2.07E-112 |
| ANXA1 | 0.501 | 0.585 | 0.347 | 5.13E-120 |
| ADIRF | 0.498 | 0.466 | 0.215 | 5.40E-152 |
| TCN1 | 0.417 | 0.299 | 0.139 | 1.13E-76 |
| CDC42EP5 | 0.413 | 0.500 | 0.314 | 1.92E-92 |
| UCA1 | 0.401 | 0.493 | 0.331 | 2.84E-62 |
| PLP2 | 0.400 | 0.406 | 0.163 | 2.09E-154 |
| KLK6 | 0.391 | 0.782 | 0.677 | 4.49E-71 |
| KLK7 | 0.364 | 0.588 | 0.470 | 1.41E-45 |
| SLCO1B3 | 0.331 | 0.430 | 0.245 | 2.81E-83 |
| LGALS1 | 0.327 | 0.268 | 0.166 | 1.54E-31 |
| SLC14A1 | 0.302 | 0.333 | 0.193 | 1.27E-47 |
| MUC1 | 0.272 | 0.391 | 0.253 | 4.18E-45 |
| HLA-E | 0.266 | 0.618 | 0.515 | 7.27E-41 |
| SEMA6A | -0.256 | 0.117 | 0.289 | 8.60E-88 |
| PLCG2 | -0.258 | 0.311 | 0.454 | 2.15E-45 |
| PRSS23 | -0.261 | 0.403 | 0.549 | 1.64E-51 |
| EPHB2 | -0.261 | 0.270 | 0.436 | 1.94E-62 |
| SNHG17 | -0.272 | 0.127 | 0.298 | 7.24E-87 |
| PROX1 | -0.274 | 0.229 | 0.386 | 2.25E-57 |
| SNHG5 | -0.279 | 0.575 | 0.736 | 1.13E-69 |
| MSI2 | -0.281 | 0.569 | 0.691 | 1.92E-50 |
| GABPB1-AS1 | -0.291 | 0.584 | 0.693 | 3.64E-47 |
| GPR155 | -0.292 | 0.083 | 0.259 | 3.38E-106 |
| DACH1 | -0.294 | 0.180 | 0.364 | 2.14E-85 |
| MUC5B | -0.295 | 0.055 | 0.201 | 4.52E-91 |
| RCN1 | -0.295 | 0.525 | 0.681 | 2.49E-64 |
| TUBB2B | -0.296 | 0.079 | 0.253 | 1.07E-103 |
| THBS2 | -0.307 | 0.374 | 0.518 | 1.34E-43 |
| CDK6 | -0.308 | 0.623 | 0.753 | 9.39E-61 |
| PRSS2 | -0.313 | 0.159 | 0.296 | 1.04E-49 |
| PALD1 | -0.322 | 0.208 | 0.421 | 3.66E-106 |
| NKD1 | -0.331 | 0.336 | 0.518 | 4.36E-73 |
| CCND2 | -0.348 | 0.445 | 0.596 | 3.08E-57 |
| BTG2 | -0.351 | 0.514 | 0.644 | 1.16E-49 |
| RETNLB | -0.357 | 0.010 | 0.213 | 5.11E-194 |
| PLCB4 | -0.359 | 0.350 | 0.497 | 7.21E-58 |
| CTNNB1 | -0.364 | 0.684 | 0.805 | 1.96E-85 |
| RGMB | -0.371 | 0.280 | 0.474 | 4.54E-87 |
| ACAA2 | -0.375 | 0.352 | 0.504 | 2.10E-63 |
| SNHG14 | -0.378 | 0.373 | 0.548 | 1.69E-76 |
| FUT9 | -0.385 | 0.355 | 0.558 | 6.16E-98 |
| TUBA1A | -0.390 | 0.296 | 0.501 | 1.53E-91 |
| TGFBI | -0.399 | 0.371 | 0.581 | 2.11E-103 |
| ATF3 | -0.401 | 0.419 | 0.558 | 2.96E-57 |
| AMACR | -0.424 | 0.283 | 0.499 | 2.15E-105 |
| ITPR2 | -0.424 | 0.363 | 0.583 | 3.59E-113 |
| LGR5 | -0.460 | 0.328 | 0.558 | 3.33E-124 |
| GSTM3 | -0.511 | 0.289 | 0.602 | 1.15E-218 |
| SMOC2 | -0.533 | 0.752 | 0.865 | 7.47E-130 |
| SLC38A11 | -0.568 | 0.072 | 0.312 | 3.43E-183 |
| VCAN | -0.663 | 0.098 | 0.366 | 2.32E-194 |
| PRAC1 | -0.722 | 0.176 | 0.606 | 0.00E+00 |
| MTRNR2L8 | -1.720 | 0.332 | 0.762 | 0.00E+00 |
| PLA2G2A | -3.928 | 0.081 | 0.439 | 0.00E+00 |

| | | | | |
|---|---|---|---|---|
| KLHL24 | 0.261 | 0.434 | 0.288 | 2.25E-50 |
| GSN | 0.260 | 0.654 | 0.501 | 9.58E-55 |
| PNRC1 | 0.257 | 0.707 | 0.578 | 2.44E-47 |
| BCL2L1 | 0.256 | 0.525 | 0.397 | 1.30E-39 |
| MYH9 | 0.252 | 0.799 | 0.697 | 4.93E-41 |
| CD74 | 0.252 | 0.443 | 0.310 | 9.05E-43 |
| DYNC1H1 | 0.252 | 0.708 | 0.598 | 1.92E-44 |
| AC119673.2 | 0.251 | 0.201 | 0.031 | 2.55E-132 |
| AURKAIP1 | -0.251 | 0.541 | 0.659 | 1.05E-44 |
| ANP32B | -0.253 | 0.609 | 0.729 | 1.65E-46 |
| MRPL12 | -0.253 | 0.262 | 0.412 | 2.71E-55 |
| MZT2B | -0.256 | 0.602 | 0.735 | 2.25E-54 |
| POMC | -0.259 | 0.016 | 0.194 | 4.55E-168 |
| DCXR | -0.260 | 0.354 | 0.513 | 9.76E-56 |
| FAM162A | -0.260 | 0.566 | 0.690 | 1.18E-37 |
| ARL6IP4 | -0.260 | 0.630 | 0.733 | 6.62E-51 |
| CUTA | -0.260 | 0.530 | 0.647 | 1.63E-43 |
| HIGD2A | -0.263 | 0.505 | 0.645 | 9.35E-51 |
| HLA-E | -0.264 | 0.498 | 0.618 | 1.85E-43 |
| PRELID1 | -0.265 | 0.562 | 0.678 | 2.30E-46 |
| CCT5 | -0.268 | 0.482 | 0.607 | 1.01E-44 |
| SH3BGRL3 | -0.268 | 0.705 | 0.810 | 4.39E-46 |
| ATP5MC1 | -0.269 | 0.532 | 0.659 | 4.05E-45 |
| TRMT112 | -0.277 | 0.517 | 0.656 | 6.14E-57 |
| PRDX6 | -0.279 | 0.611 | 0.742 | 1.08E-62 |
| S100P | -0.280 | 0.080 | 0.184 | 1.12E-43 |
| CREB3L2 | -0.284 | 0.234 | 0.387 | 1.77E-61 |
| PEG10 | -0.285 | 0.293 | 0.425 | 8.56E-40 |
| METRN | -0.285 | 0.365 | 0.520 | 1.77E-60 |
| GTF3A | -0.287 | 0.329 | 0.492 | 1.46E-63 |
| SLCO1B3 | -0.287 | 0.274 | 0.430 | 1.18E-57 |
| GSTO1 | -0.290 | 0.452 | 0.599 | 1.66E-59 |
| PSMB9 | -0.294 | 0.156 | 0.337 | 3.92E-91 |
| PIGR | -0.297 | 0.215 | 0.376 | 3.47E-62 |
| RNASE1 | -0.298 | 0.434 | 0.592 | 7.55E-45 |
| MGST1 | -0.302 | 0.447 | 0.616 | 4.81E-66 |
| RANBP1 | -0.305 | 0.437 | 0.584 | 1.09E-55 |
| TSPO | -0.311 | 0.734 | 0.850 | 1.28E-77 |
| ATP5F1B | -0.315 | 0.704 | 0.807 | 3.03E-72 |
| APRT | -0.319 | 0.669 | 0.783 | 2.30E-68 |
| CYC1 | -0.319 | 0.328 | 0.512 | 1.81E-81 |
| NME1 | -0.321 | 0.402 | 0.551 | 2.91E-61 |
| EPB41L2 | -0.321 | 0.482 | 0.647 | 5.92E-72 |
| TMEM141 | -0.324 | 0.432 | 0.621 | 6.14E-69 |
| SNRPB | -0.330 | 0.609 | 0.734 | 1.28E-66 |
| EIF5A | -0.334 | 0.542 | 0.683 | 7.11E-69 |
| MDH2 | -0.335 | 0.511 | 0.674 | 1.54E-86 |
| EIF4EBP1 | -0.335 | 0.296 | 0.485 | 2.12E-82 |
| ECH1 | -0.335 | 0.493 | 0.668 | 7.07E-86 |
| COX5A | -0.340 | 0.743 | 0.852 | 2.84E-90 |
| C1QBP | -0.351 | 0.562 | 0.693 | 7.72E-73 |
| GUK1 | -0.367 | 0.652 | 0.791 | 3.33E-96 |
| CCDC85B | -0.381 | 0.423 | 0.618 | 1.46E-95 |
| ZNF511 | -0.383 | 0.447 | 0.643 | 1.42E-87 |
| ENO1 | -0.384 | 0.843 | 0.959 | 3.20E-116 |
| HMGA1 | -0.391 | 0.676 | 0.799 | 1.06E-97 |
| RPL22L1 | -0.396 | 0.481 | 0.649 | 1.57E-83 |
| AQP5 | -0.403 | 0.750 | 0.866 | 1.58E-68 |
| UCA1 | -0.414 | 0.344 | 0.493 | 1.05E-54 |
| MTRNR2L1 | -0.419 | 0.026 | 0.243 | 6.83E-203 |
| NQO1 | -0.431 | 0.376 | 0.609 | 7.52E-122 |
| ATP5MC3 | -0.436 | 0.808 | 0.910 | 4.62E-147 |
| PRDX5 | -0.446 | 0.617 | 0.799 | 7.50E-142 |
| TKT | -0.450 | 0.788 | 0.908 | 2.95E-160 |
| SLC14A1 | -0.453 | 0.136 | 0.333 | 5.40E-104 |
| CHCHD10 | -0.461 | 0.387 | 0.651 | 2.08E-163 |
| HLA-C | -0.461 | 0.830 | 0.932 | 1.68E-152 |
| LDHA | -0.462 | 0.771 | 0.925 | 1.69E-123 |
| SLC25A5 | -0.496 | 0.761 | 0.895 | 1.14E-181 |
| LGALS4 | -0.517 | 0.752 | 0.904 | 1.52E-171 |
| CDC42EP5 | -0.521 | 0.240 | 0.500 | 1.46E-165 |
| ATP5F1D | -0.567 | 0.756 | 0.900 | 1.10E-238 |
| THBS2 | -0.595 | 0.090 | 0.374 | 1.96E-226 |
| AGR2 | -0.628 | 0.656 | 0.871 | 2.20E-224 |
| PAX8-AS1 | -0.681 | 0.025 | 0.430 | 0.00E+00 |
| IFI27 | -0.799 | 0.147 | 0.440 | 3.31E-238 |
| HLA-B | -0.843 | 0.775 | 0.952 | 0.00E+00 |
| PLIN2 | -0.937 | 0.674 | 0.843 | 6.41E-209 |
| XIST | -1.073 | 0.010 | 0.384 | 0.00E+00 |
| OLFM4 | -1.238 | 0.184 | 0.607 | 0.00E+00 |

| | | | | |
|---|---|---|---|---|
| CCDC85B | -0.294 | 0.423 | 0.587 | 1.33E-64 |
| EIF4EBP1 | -0.295 | 0.296 | 0.486 | 6.22E-82 |
| EIF5A | -0.298 | 0.542 | 0.687 | 2.84E-64 |
| FABP5 | -0.299 | 0.249 | 0.415 | 2.63E-68 |
| ODC1 | -0.304 | 0.368 | 0.552 | 8.30E-79 |
| SATB2 | -0.306 | 0.018 | 0.205 | 5.80E-180 |
| CASC19 | -0.309 | 0.330 | 0.530 | 3.17E-78 |
| PDIA6 | -0.310 | 0.792 | 0.895 | 1.70E-91 |
| ATP5MC1 | -0.320 | 0.532 | 0.695 | 1.16E-77 |
| GTF3A | -0.323 | 0.329 | 0.524 | 1.63E-91 |
| CHCHD10 | -0.327 | 0.387 | 0.599 | 1.07E-101 |
| S100P | -0.331 | 0.080 | 0.189 | 1.56E-50 |
| PALD1 | -0.331 | 0.205 | 0.421 | 3.31E-118 |
| APRT | -0.331 | 0.669 | 0.799 | 3.70E-90 |
| SLC25A5 | -0.331 | 0.761 | 0.875 | 8.26E-94 |
| PEG10 | -0.334 | 0.293 | 0.485 | 1.89E-82 |
| PRMT1 | -0.335 | 0.455 | 0.638 | 4.70E-92 |
| TMEM141 | -0.340 | 0.432 | 0.618 | 1.71E-73 |
| RETNLB | -0.344 | 0.020 | 0.213 | 2.89E-185 |
| PA2G4 | -0.346 | 0.569 | 0.700 | 3.73E-65 |
| RCN1 | -0.346 | 0.490 | 0.681 | 2.65E-91 |
| PLCB4 | -0.349 | 0.349 | 0.497 | 2.92E-61 |
| TSPAN5 | -0.362 | 0.463 | 0.670 | 6.10E-111 |
| CCT5 | -0.364 | 0.482 | 0.678 | 2.88E-103 |
| ANOS1 | -0.365 | 0.041 | 0.288 | 1.37E-226 |
| PRSS23 | -0.366 | 0.335 | 0.549 | 6.93E-112 |
| SNU13 | -0.371 | 0.581 | 0.739 | 1.02E-101 |
| LGR5 | -0.382 | 0.353 | 0.558 | 6.26E-96 |
| CCND2 | -0.382 | 0.367 | 0.596 | 2.66E-97 |
| SNRPB | -0.389 | 0.609 | 0.763 | 2.37E-108 |
| CKB | -0.398 | 0.233 | 0.437 | 1.42E-108 |
| NME1 | -0.399 | 0.402 | 0.606 | 5.84E-111 |
| ATP5F1D | -0.400 | 0.756 | 0.875 | 5.57E-139 |
| C1QBP | -0.409 | 0.562 | 0.742 | 3.24E-126 |
| RANBP1 | -0.413 | 0.437 | 0.642 | 1.64E-109 |
| TKT | -0.416 | 0.788 | 0.904 | 1.12E-153 |
| HLA-B | -0.419 | 0.775 | 0.901 | 3.91E-127 |
| PRDX5 | -0.420 | 0.617 | 0.800 | 9.80E-144 |
| LDHA | -0.439 | 0.771 | 0.883 | 1.11E-91 |
| PPIB | -0.455 | 0.814 | 0.918 | 4.34E-172 |
| RPL22L1 | -0.459 | 0.481 | 0.675 | 1.34E-121 |
| EPB41L2 | -0.475 | 0.482 | 0.722 | 6.35E-171 |
| GABPB1-AS1 | -0.501 | 0.468 | 0.693 | 9.26E-155 |
| MTRNR2L1 | -0.513 | 0.026 | 0.304 | 7.05E-286 |
| VCAN | -0.539 | 0.136 | 0.366 | 4.75E-144 |
| TGFBI | -0.555 | 0.275 | 0.581 | 2.16E-218 |
| PLIN2 | -0.566 | 0.674 | 0.788 | 9.07E-83 |
| AGR2 | -0.612 | 0.656 | 0.877 | 5.08E-243 |
| THBS2 | -0.903 | 0.090 | 0.518 | 0.00E+00 |
| XIST | -1.514 | 0.010 | 0.465 | 0.00E+00 |
| MTRNR2L8 | -1.550 | 0.426 | 0.762 | 0.00E+00 |
| PLA2G2A | -3.520 | 0.149 | 0.439 | 3.12E-267 |

**Supplementary Table 5  Antibody panel for IMC**

| Mass | Metal | Marker | Clone | Vendor | Dilution |
|---|---|---|---|---|---|
| 141 | Pr | a-SMA | 1A4 | Fluidigm | 1000 |
| 142 | Nd | Mucin 2 | Ccp58 | Santa Cruz | 100 |
| 143 | Nd | Vimentin | RV202 | Fluidigm | 400 |
| 144 | Nd | CD14 | EPR3653 | Fluidigm | 100 |
| 145 | Nd | BCL-6 | LN22 | lovus Biologica | 25 |
| 146 | Nd | CD16 | EPR16784 | Fluidigm | 100 |
| 148 | Nd | Pan Keratin | C11 | Fluidigm | 300 |
| 149 | Sm | CD11b | EPR1344 | Fluidigm | 200 |
| 151 | Eu | CD11c | EP1347Y | Abcam | 200 |
| 152 | Sm | CD45 | 2B11 | Fluidigm | 200 |
| 153 | Eu | CD56 | Rabbit Poly Ab | Proteintech | 300 |
| 154 | Sm | IL-6 | 3154011B | Fluidigm | 100 |
| 155 | Gd | Foxp3 | 236A/E7 | Fludigim | 75 |
| 156 | Gd | CD4 | EPR6855 | Fluidigm | 100 |
| 158 | Gd | E-Cadherin | 2.40E+10 | Fluidigm | 300 |
| 159 | Tb | CD68 | KP1 | Fluidigm | 600 |
| 160 | Gd | Lysozyme C | E5 | Santa Cruz | 200 |
| 161 | Dy | CD20 | H1 | Fluidigm | 200 |
| 162 | Dy | CD8a | C8/144B | Fluidigm | 300 |
| 163 | Dy | CXCR3 | G025H7 | Fluidigm | 300 |
| 165 | Ho | CD69 | sc-373798 | Santa Cruz | 200 |
| 166 | Ho | NFKB | 3166006A | Fluidgim | 100 |
| 167 | ER | Granzyme B | EPR20129-217 | Fluidigm | 200 |
| 168 | Er | Ki67 | B56 | Fluidigm | 600 |
| 169 | Tm | Collagen I | Polyclonal | Fluidigm | 600 |
| 170 | ER | CD3 | Polyclonal (C terminal) | Fluidigm | 200 |
| 171 | Yb | CD27 | EPR8569 | Fluidigm | 100 |
| 173 | Yb | CD45RO | UCHL1 | Fluidigm | 300 |
| 174 | Yb | HLA-DR | TAL 1B5 | Abcam | 600 |
| 175 | Lu | CD38 | H-11 | Santa Cruz | 100 |
| 176 | Ho | Histone | D1H2 | Fluidigm | 1000 |
| 191/193 | Ir | DNA | -- | Fluidigm | 400 |

**Detailed information on image acquired by IMC**

| Subject ID | Slide ID | Condition | Region | Image ID | Acquisition area (ums) | Total no. cells captured per image | No. cells per image, after filtering | No. cells per sample, after filtering |
|---|---|---|---|---|---|---|---|---|
| Variant | SUR 5385 | variant | Right colon | Variant_ROI 001 | 2000 x 1100 | 15514 | 5795 | 11011 |
| | | | | Variant_ROI 002 | 2000 x 2000 | 22585 | 5216 | |
| VEO-IBD-1 | SUR 2644 | VEO IBD | Right colon | VEO-IBD-1_ROI001 | 1100 x 3500 | 18780 | 15544 | 15544 |
| VEO-IBD-2 | SUR 2956 | VEO IBD | Colon | VEO-IBD-2_ROI001 | 600 x 2000 | 6198 | 6198 | 15984 |
| | | | | VEO-IBD-2_ROI002 | 1400 x 1300 | 9790 | 9786 | |
| VEO-IBD-3 | SUR 5419 | VEO IBD | Colon | VEO-IBD-3_ROI002 | 1000 x 2000 | 12867 | 12764 | 25129 |
| | | | | VEO-IBD-3_ROI003 | 900 x 2000 | 12365 | 12365 | |
| Control-4 | SUR 6941 | Healthy | Colon | Control-4_ROI003 | 900 x 1750 | 5721 | 5083 | 12468 |
| | | | | Control-4_ROI004 | 350 x 2500 | 4551 | 4545 | |
| | | | | Control-4_ROI005 | 600 x 1000 | 3066 | 2840 | |
| Control-3 | S15 9148 | Healthy | Left colon | Control-3_ROI001 | 1000 x 2884 | 10051 | 6767 | 6767 |
| Control-2 | SUR 3497 | Healthy | Left colon | Control-2_ROI001 | 800 x 2500 | 9353 | 9108 | 11917 |
| | | | | Control-2_ROI002 | 500 x 1500 | 2809 | 2809 | |
